# A multi-dimensional, time-lapse, high content screening platform applied to schistosomiasis drug discovery

**DOI:** 10.1101/872069

**Authors:** Steven Chen, Brian M Suzuki, Jakob Dohrmann, Rahul Singh, Michelle R Arkin, Conor R Caffrey

**Affiliations:** Department of Pharmaceutical Chemistry and Small Molecule Discovery Center, University of California, San Francisco, CA 94143; Center for Discovery and Innovation in Parasitic Diseases, Department of Pathology, University of California, San Francisco, CA 94158; Center for Discovery and Innovation in Parasitic Diseases, Skaggs School of Pharmacy and Pharmaceutical Sciences, University of California, San Diego, La Jolla, CA 92093; Department of Computer Science, San Francisco State University, San Francisco, CA 94132

## Abstract

Approximately 10% of the world’s population is at risk of schistosomiasis, a disease of poverty caused by the *Schistosoma* parasite. To facilitate drug discovery for this complex flatworm, we developed an automated high-content screen to quantify the multidimensional responses of *Schistosoma mansoni* post-infective larvae (somules) to chemical insult. We describe an integrated platform to process worms at scale, collect time-lapsed, bright-field images, segment highly variable and touching worms, and then store, visualize, and query dynamic phenotypes. To demonstrate the methodology, we treated somules with seven drugs that generated diverse responses and evaluated 45 static and kinetic response descriptors relative to concentration and time. For compound screening, we used the Mahalanobis distance to compare multidimensional phenotypic effects induced by 1,323 approved drugs. Overall, we characterize both known anti-schistosomals and identify new bioactives. Apart from facilitating drug discovery, the multidimensional quantification provided by this platform will allow mapping of chemistry to phenotype.

## Introduction

The *Schistosoma* blood fluke (helminth) causes schistosomiasis, a neglected tropical disease (NTD) ^1–3^ that infects over 200 million people and puts more than 700 million people at risk of infection in 78 countries ^4–6^. Parasite eggs cause chronic inflammatory and fibrotic responses that impair visceral and/or urogenital organ function; co-morbidities include increased risks for bladder cancer and HIV ^7, 8^. Praziquantel (PZQ) is the only available drug for schistosomiasis. Although reasonably active against mature schistosomes, PZQ displays little to no efficacy against developing parasites ^9, 10^. Also, increased utilization of PZQ raises concerns that drug resistance will emerge. Thus, new drugs are needed ^11^.

Anthelmintic drug discovery has traditionally relied on phenotypic screens using parasites in culture or in small animal models ^12^. Primary screening of cultured schistosomes has often used post-infective larvae (called schistosomula or somules) that can be obtained in their thousands to tens of thousands from vector snails for relatively little effort and cost, in contrast to adult worms that can only be harvested in low numbers (hundreds) from small mammals. Single-metric assays, in which somules are scored as alive or dead, have been reported. (*e.g*., ^13^ for review). However, single-metric approaches have a number of drawbacks; in some cases, even the clinically used drugs do not score as active in these assays ^13, 14^. High-content imaging of live somules offers the potential to visualize complex and non-lethal (but potentially therapeutically relevant) phenotypic responses to drug treatment.

As part of our research program to develop methods for anti-schistosomal drug discovery ^15–17^, we report a fully integrated, automated and multiparametric image-analysis platform for high throughput phenotyping of living parasites. Starting with a set of seven drugs known to induce changes in shape and motion ^15, 18^, we describe a set of protocols to quantify those changes as a function of time and concentration. We then demonstrate the utility of the method for high-throughput screening using a set of 1,323 approved drugs. Our approach offers key advances in method integration, including several of general utility to the drug screening/imaging community: a) automated liquid handling of 100 µm-sized organisms, b) manipulation of the focal plane to facilitate identification of low-contrast, variable and touching objects, c) time-lapsed tracking to define frequencies and rates of motion, d) a public system for storage, visualization and querying of the complex phenotypic data, and e) use of a statistical metric (the Mahalanobis distance, *d_M_*) to compare multidimensional phenotypes for high-throughput screening.

## RESULTS

### High-throughput sample handling for *S. mansoni* somules

We selected somules for primary assays as we can obtain 10^4^ – 10^5^ somules/week from freshwater vector snails. As somules (∼300 x 150 µm) rapidly settle out of solution, we used a magnetic tumble stirrer containing eight stirring paddles to maintain worms in suspension (**Fig. 1**) and allow their transfer using 96- or 384- channel pipets from a reservoir containing 200 mL of media and 40,000 somules. Plate geometry and number-of-somules/well were optimized for accurate imaging and counting (**Supplementary Fig. 1a**). In particular, u-bottom 96w plates concentrated the parasite as a monolayer into one central field of view, thus facilitating automated imaging. Forty somules per well allowed us to maximize the number of worms being tested without reducing our ability to count them due to overlapping (**Supplementary Fig. 1a**). Routine robotic protocols then dispensed compound and shuttled plates between an incubator (Cytomat 2C) and a high-content imager (GE IN Cell Analyzer 2000) for data-collection.

**Figure 1.**
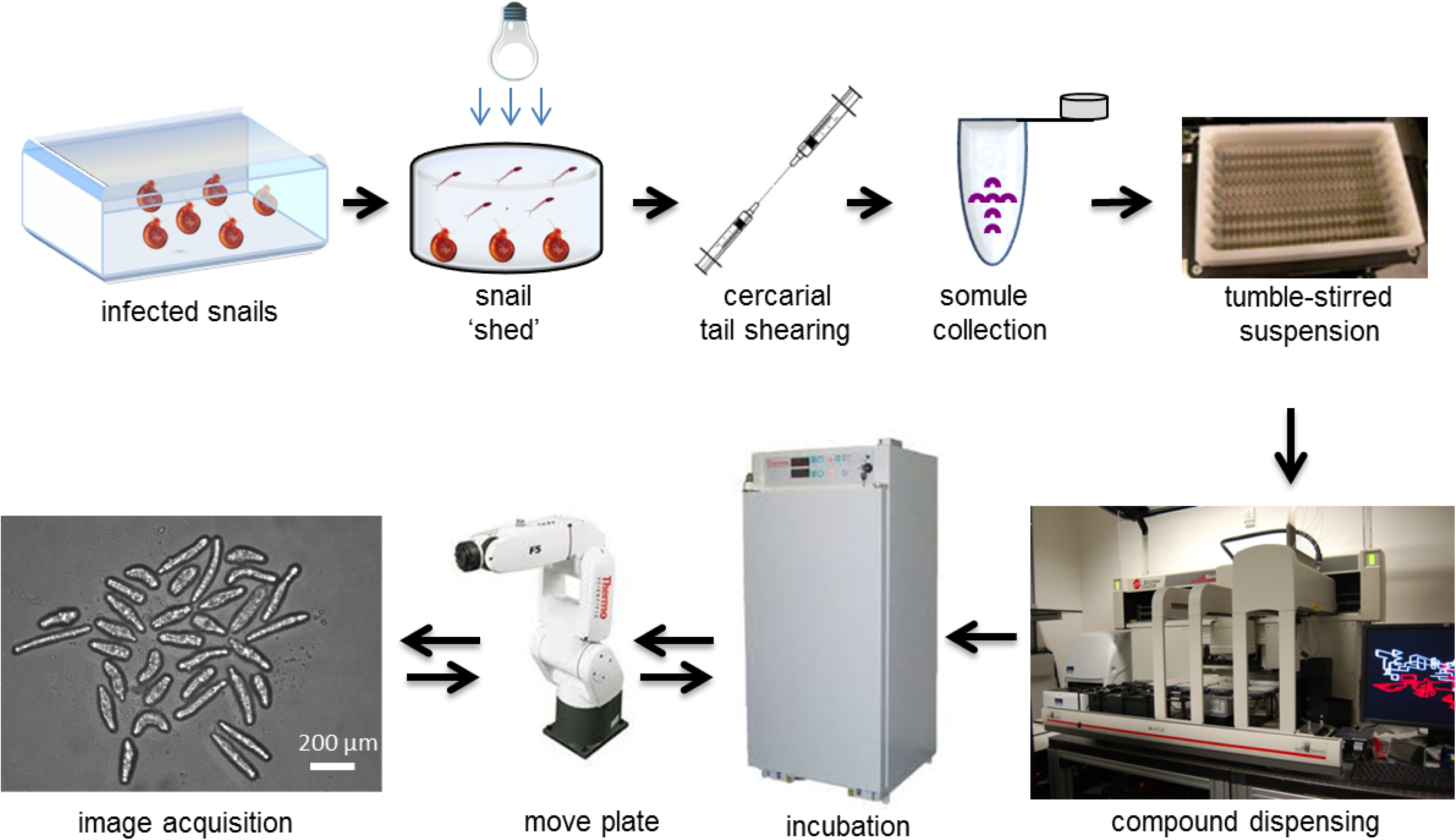
Assay workflow. Infected *Biomphalaria glabrata* vector snails were stimulated under light to release (‘shed’) 10^4^–10^5^ *Schistosoma mansoni* infective larvae (cercariae) per week. Cercariae were mechanically converted to post-infective larvae (schistosomula or somules) using a 20-gauge double-headed syringe needle. After washing to remove cercarial tails, somules were suspended in a paddle-stirred reservoir. Somules were then dispensed into 96-well u-bottom clear plates at 40 units/well/200 µL Basch medium. Compounds were added using a 96-channel pin tool. Plates were maintained at 5% CO_2_ and 37 °C. At specified time points, a 6-axis robotic arm transferred the plates to the high-content imager.

### Imaging schistosomes by automated, bright-field image analysis

Bright-field, time-lapsed images were generated for control and drug-treated somules using a 10x objective. Every 24 h for 3 days, images were collected at 1.66 Hz (the maximum frame rate for the IN Cell Analyzer 2000) to generate 20 s video recordings. We employed bright-field imaging as it is mechanism-agnostic, non-invasive and fast, and because the schistosome is not yet routinely amenable to the transgenic incorporation of fluorescent proteins. However, in bright-field, somules do not present a high-contrast edge relative to background, thus limiting object segmentation (detection of the object’s outline). We, therefore, lowered the focal plane 40 µm below the bottom of the well in order to artificially generate a dark edge that facilitated segmentation without a significant loss of interior density features (texture; **Supplementary Fig. 1b**).

We observed two strong distributions of somules during our studies. Initially, parasites had a translucent body with a discernable outline. However, under the influence of toxic compounds, worms could become progressively opaque, such that the worm outline was indistinguishable from the interior of the worm. This opacity was associated with degenerating/dying parasites, with the transition from “clear” to “opaque” being irreversible. To accurately identify both classes of worms, we segmented the somules using three customized protocols that were optimized to detect somules independently by considering (a) only the worm outline, (b) the worm’s interior, including the inner edge or (c) only the worm’s interior, excluding the edge (**Fig. 2**). The most time-lapse-persistent segmented area obtained from these protocols was selected as a true somule. Each somule was then described using 15 features, including those that define size, shape, texture and color (for terminology, see **Supplementary Table 1**).

**Figure 2.**
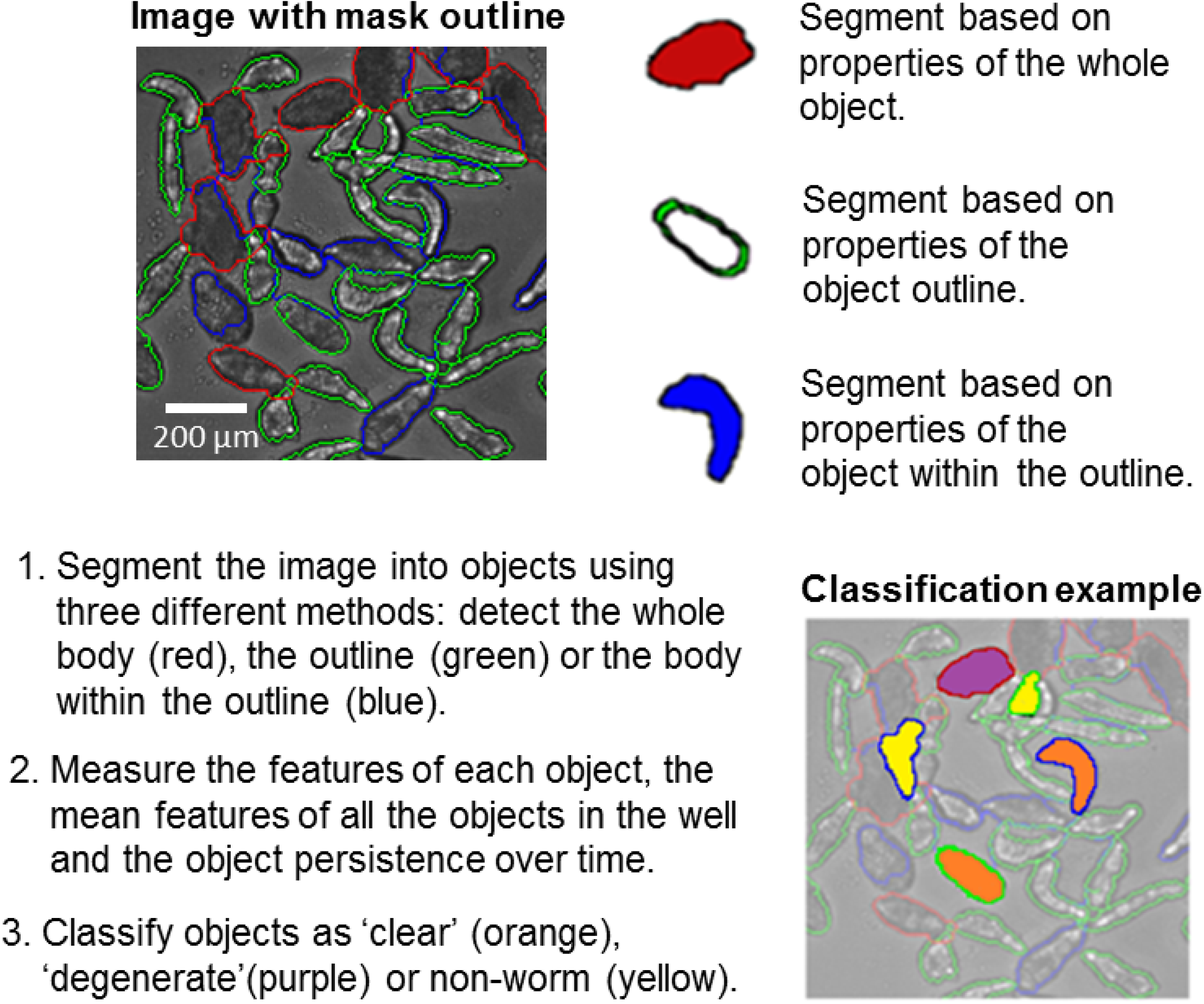
Summary of segmentation and data post-process workflow. Details provided in **Supplementary Information**. Acquired images (30 time-lapse images per well) were loaded into the IN Cell Developer software and segemented. ‘Clear’ worms, ‘opaque’ worms and non-worm objects were classified based on features computed on a subset of the data containing 20,000 worm and non-worm objects.

We classified the segmented objects into groups – ‘clear’ (translucent) worms, ‘opaque’ (degenerate/dying) worms and ‘non-worm’ objects. Compared to the clear worms, degenerate worms had a lower mean intensity level (1,822 ± 49 *vs.* 2,143 +/- 42 levels), a lower standard deviation of levels in the mask (227 ± 50 *vs.* 556 ± 49 levels), a larger area (6,421 ± 599 *vs.* 3,962 ± 447 µm^2^) and a larger form factor (0.66 ± 0.05 *vs.* 0.52 ± 0.06). The percentage of somules that were classified as degenerate yielded a ‘degeneracy’ score.

Our image-segmentation protocols identified the somules with a precision of 88% and a recall of 95% for 58,456 worms in the seven-drug-set experiment described below. These results were confirmed by visual inspection (**Supplementary Fig. 1, Panel C**). Using the plating density of 40 worms/well, we observed very few overlapping worms (< 0.47%; **Supplementary Fig. 1, Panel D**). The precision and recall values, without the need to remove touching worms, represent a significant improvement over the state-of-the-art ^19^. Data collection, segmentation and classification protocols are described in the **Supplementary Information.**

Once worms were classified as clear or degenerate, the 15 calculated features were evaluated across three modes – static, rate and frequency. In the *static* mode, we considered feature measurements in each frame independently, *e.g.,* worm length. Time-dependent changes in these features were measured using *rate* and *frequency*. *Rate* measured the magnitude of a change in a feature, such as worm length per unit time. Motion could also be characterized by how often the time-dependent measurement changes sign or direction, *e.g*., the worm becomes longer, then shorter. To capture this aspect, we defined the *frequency* of a feature to be the number of times the sign of the difference in consecutive feature values changed (*i.e.,* change in sign/unit time). Due to slight offsets in the camera between frames, we set a motion threshold based on a non-moving reference (worms paralyzed by metrifonate). As somules showed little translational movement in the u-bottom wells, we did not record their displacement. When combined, the static, rate and frequency modes for each of the 15 features yielded 45 measurements (which we define as descriptors) for each somule (see next section).

Two statistical approaches are used to evaluate the significance of changes in worm phenotypes in response to drug treatment. First, the mean and standard deviation for each descriptor for all somules within a well are computed and normalized to determine the Glass Effect Size (ES) ^20, 21^:

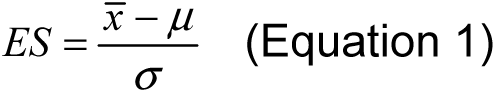

where 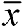 is the mean descriptor due to drug exposure, μ is the DMSO mean and σ is the standard deviation of the descriptor for parasites in DMSO. Being dimensionless, ES is useful in comparing effects across different features. In addition to evaluating individual descriptors, we compared parasites in this descriptor space using the Mahalanobis distance (*d_M_*) ^22^, which measures the multi-dimensional, scale invariant distance between a test well and a standard condition, *e.g.,* DMSO-treated somules. It is calculated by 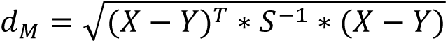 where *X* is the test well vector, *Y* the DMSO vector, *S* the covariance matrix and *T* indicates that the vector should be transposed. The metrics, *d_M_* and ES, are similar in that both measure a distance from the DMSO reference. However, ES measures the distance for just one feature, whereas *d_M_* measures the distance for a group of variables. Moreover, *d_M_* is not dependent on the measurement unit and can identify test wells that have one large difference or multiple small differences compared to DMSO-treated controls.

### SchistoView: query-visualization of phenotypic screening data

We developed SchistoView (**Fig. 3**) ^23^ which comprises a graphical user interface supported by a MySQL database. SchistoView allows users to visualize and query concentration-and time-dependent somule response data, from computed statistics for a given well to features for individual somules. **Figure 3** shows a screenshot of Schistoview and describes the features of the GUI. Further details are provided in the **Supplementary Information**.

**Figure 3.**
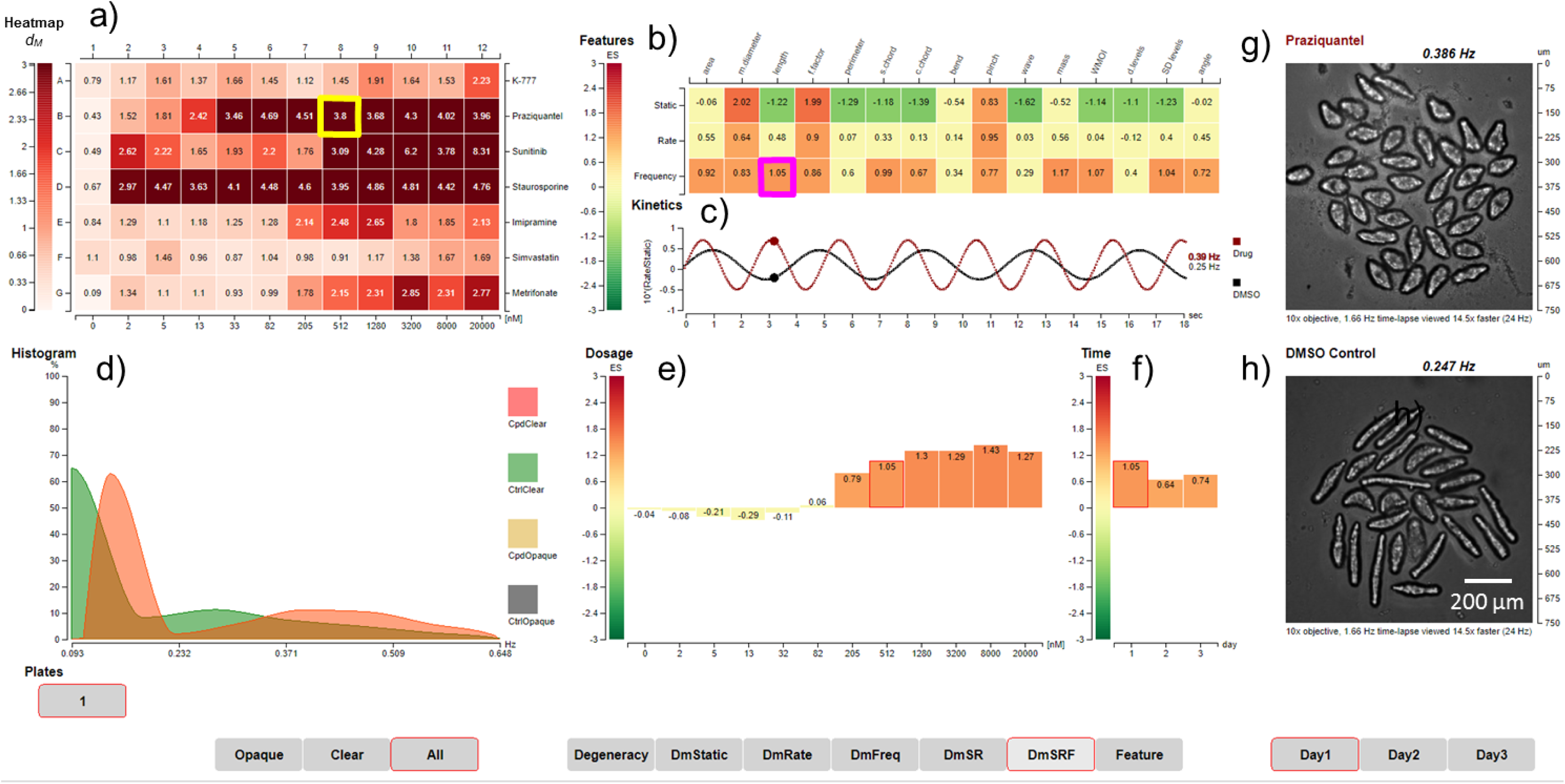
The SchistoView graphical user interface ^23^. Selected data at 2 h illustrate the hierarchical approach to visualization. (**a**) Heat map of Mahalanobis distances (*d_M_*) for seven test drugs over an 11-point, 2.5-fold dilution series (2 nM – 20 μM). The test drugs are K11777, PZQ, sunitinib, staurosporine, imipramine, simvastatin and metrifonate. DMSO controls are in column 1 and the average *d_M_* for each DMSO control is shown. A *d_M_* of 1.61 is 3 SD from the DMSO control. Clicking on well B8 of the heat map (512 nM PZQ – the yellow square) populates panels (**b**) - (**g**) in the GUI. (**b**) Heat map of the effect sizes (ES) for each of 15 features (columns) plotted *vs*. static, rate or frequency modes (rows); a ‘descriptor’ is the combination of a given feature and mode. Clicking on a descriptor, *e.g*., frequency of change-in-length (the magenta square) populates panels (**c**) – (**f**). (**c**) Calculated waveforms (kinetics) defined by the mean length (baseline), range of length (amplitude) and frequency of length contraction (frequency). DMSO control worms (black line) are longer (higher baseline) and slower moving (lower frequency) than those treated with 512 nM PZQ at 2 h (red line). (**d**) Histogram displaying the distribution of frequency-of-length contraction for DMSO controls (green) and PZQ-treated worms (orange). **(e)** Dosage bar graph depicting the ES for the frequency-of-length contraction after 2 h with 11 PZQ concentrations. (**f**) Time bar graph depicting the ES for the frequency-of-length contraction after PZQ (512 nM) treatment across the three days indicated. (**g**) First image from time-lapsed movie of well B8 highlighted in (**a**); in the live SchistoView, the 30-frame movie is shown. (**h**) First image from time-lapsed movie of a DMSO-treated well; in the live SchistoView, the 30-frame movie is shown. Buttons at the bottom are used in SchistoView to select the plate (1), the somules analyzed (Opaque, Clear, All), the mode displayed in panel (**a**) (*i.e.,* degeneracy, *d_M_* for static features, *d_M_* for rate, *d_M_* for frequency, *d_M_* for static and rate, *d_M_* for static, rate and frequency or a single feature), and the day of the experiment (Day 1, 2 or 3).

### Exploring the parasite’s multivariate responses using known anti-schistosomal compounds

We tested the time-lapsed imaging platform with seven compounds that induced diverse changes in the parasite ^15, 18^. Somules were exposed to an 11-point, 2.5-fold dilution series (from 2 nM to 20 μM) of compounds in quadruplicate and images were captured after 2, 24 and 48 h. Raw images, collected after 24 h of the first frame in each well, are shown in **Supplementary Fig. 2**. Images were segmented and data extracted as described above. The results highlight important features of the imaging methodology, the depth of analysis offered and the underlying biology of the schistosome parasite.

In **Fig. 4** and **Supplementary Fig. 3**, the time- and concentration-dependent effects of drugs on worm behavior are visualized using heat maps extracted from SchistoView. Note that the *d_M_* values shown do not necessarily smoothly change with increasing concentration of a drug. This is due to individual features showing maximum changes at different concentrations. For some compounds (*e.g*., sunitinib, staurosporine and PZQ) it is also noteworthy that at later timepoints *d_M_* values can be smaller yet still significant. The biological factors responsible may include compound metabolism and baseline physical changes as the schistosome adapts to its environment.

**Figure 4.**
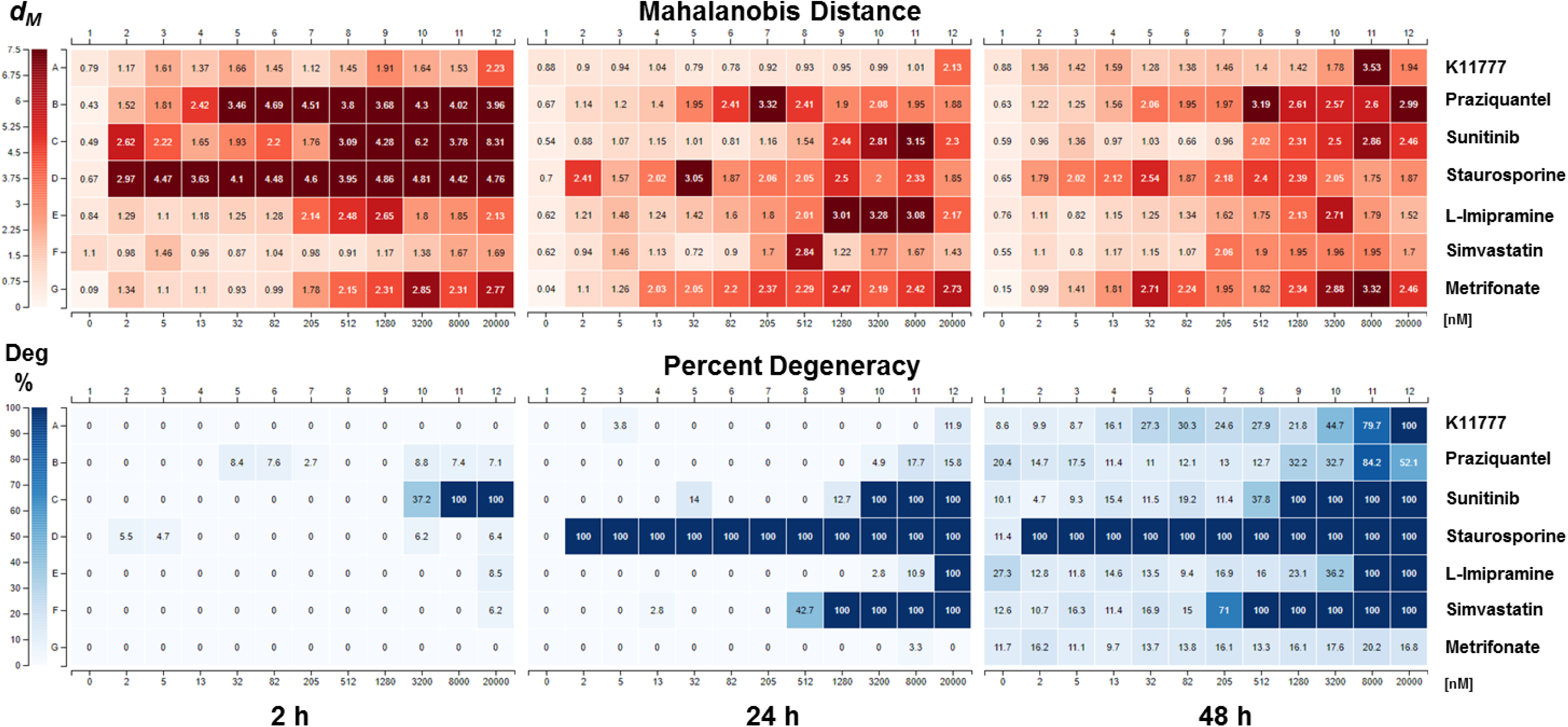
Heat map displaying *d_M_* and degeneracy for seven test drugs arrayed over a 2.5-fold dilution series from 20 μM to 2 nM. *d_M_* values were calculated relative to DMSO controls. *d_M_* values of 1.61, 1.36 and 1.28 are 3 SD from the corresponding controls at 2, 24 and 48 h, respectively. Results presented are aggregated from four wells per treatment group (40 somules/well). For most compounds, phenotypes appeared quickly (within 2 or 24 h), relative to the appearance of degeneracy (pronounced at 48 h). Notably, the two approved anti-schistosomal drugs, PZQ and metrifonate, do not induce significant degeneracy under these conditions. It is also noteworthy that *d_M_* values can be smaller at later time points. Values for *d_M_* were calculated using the eight DMSO control wells collected closest in time (+/-20 min) to the target well. Due to well averaging, the DMSO control well *d_M_* value shown in the figure is > 0 (*d_M_* cannot be negative). This small, positive value allows us to define “activity” as *d_M_* > 3 SD from the mean value for the DMSO control wells.

Using *d_M_* as a summary of overall static and kinetic phenotypic changes, concentration-dependent effects of drug exposure were already significant by the first time point (2 h) for five of the seven drugs (*i.e*., *d_M_* > 1.61, which is 3 SD from the *d_M_* of the DMSO controls) ^24^. These drugs included PZQ, the kinase inhibitors sunitinib and staurosporine, the anticholinergic imipramine, and the acetylcholine esterase inhibitor (and former anti-schistosomal drug) metrifonate. Notably, very little degeneracy was observed at the first time point, except for sunitinib at the two highest concentrations. After 24 h, *d_M_* remained elevated relative to DMSO, and degeneracy became apparent for lower concentrations of sunitinib, staurosporine and the HMG-CoA reductase inhibitor simvastatin. Finally, by 48 h, concentration-dependent degeneracy became apparent for five of the compounds with the notable exceptions of PZQ and metrifonate. The absence of cidal activity for these two drugs was consistent with their primary activity as paralytics (see below). Thus, the ability to capture phenotypic changes by *d_M_* (i) afforded a rapid and deep assessment of anti-schistosomal activity that was independent of degeneracy/death and (ii) added essential value by identifying highly relevant anti-schistosomals that did not induce degeneracy, including the two clinically used drugs, PZQ and metrifonate. Finally, combining static and dynamic descriptors into the *d_M_* provided a more sensitive readout of phenotypic change than either modality alone (**Supplementary Fig. 3**).

Our imaging platform quantified drug-induced increases and decreases in parasite motility. Previously, L-imipramine was visually assessed to induce hypermotility ^15^. We confirmed this finding and, for the first time (to our knowledge), quantified the response. As shown in **Fig. 5a**, imipramine induced a concentration-dependent increase in movement after 2 h between 10 nM and 1 µM (EC_50_ = 100 nM) followed by decreased motility at higher concentrations. This hypermotility was measured as an increasing rate-of-change in length (ES = 1.35; 6 µm/s at ∼ 1 µM *vs.* 2 µm/s for DMSO) and increasing frequency (ES = 1; 0.36 Hz at 1 µM *vs.* 0.21 Hz for DMSO). Importantly, the fitted median length never exceeded the minimum or maximum value of DMSO-treated somules at concentrations <8 µM (within ± 0.31 ES); hence, had we only relied on static image-based analysis, imipramine might have been missed as an active compound. By contrast, metrifonate at 2 h (**Fig. 5b**) caused flacid paralysis (ES_frequency_ = -1; 0.09 Hz at 1.3 µM) that was mirrored by an increase in length (ES_static_ = 1.5; 166 µm for 1.3 µM metrifonate *vs.* 137 µm for DMSO). This paralysis was consistent with the inhibition of acetylcholine esterase by metrifonate^25^. For worms treated with PZQ or metrifonate, *d_M_* remained high at 24 and 48 h at concentrations where degeneracy remained low.

**Figure 5.**
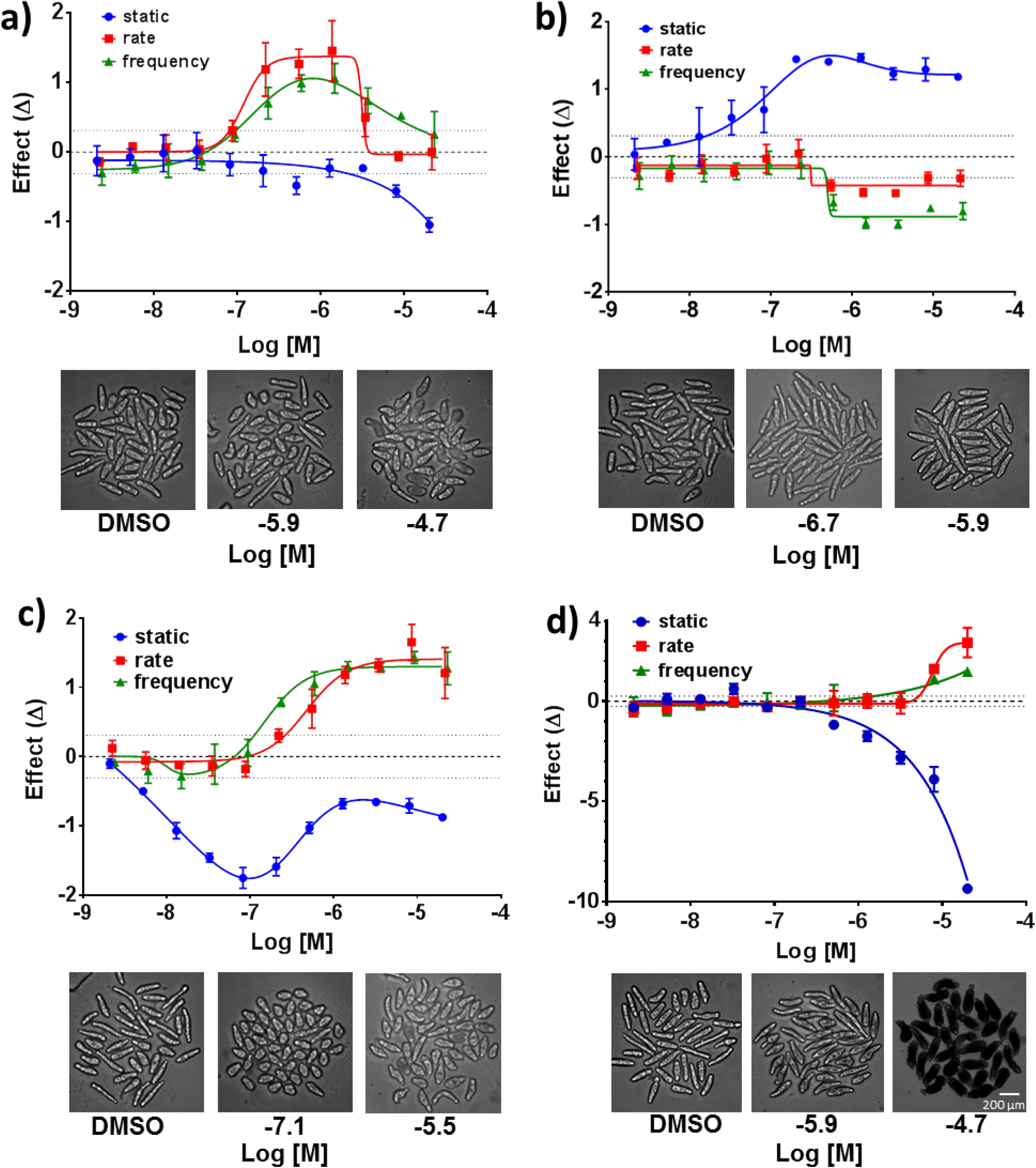
Concentration-dependent phenotypic responses to selected anti-schistosomal drugs from Figure 4 after 2 h of treatment. Compound concentrations (2 nM – 20 µM) are plotted *vs*. effect size (ES). The graphs display changes in mean length (blue), rate-of-change of length (red) and frequency of changes in length/contractions (green) for (**a**) L-imipramine, (**b**) metrifonate and (**c**) PZQ. (**d**) The graph for sunitinib displays changes in mean density levels (blue), rate-of-change of density levels (red) and frequency of changes in density levels (green). Error bars represent the standard deviation from four replicate wells at each concentration. Dotted lines above and below baseline represent 3 SD from the mean of the DMSO-treated wells. Images of the somules at selected drug concentrations are shown below each graph.

Despite the centrality of PZQ for the treatment of schistosomiasis, its mechanism of action is not completely elucidated and is likely complex, leading to Ca^++^ influx and spastic paralysis ^9, 26^. This complexity is reflected in our phenotypic analysis (**Fig. 5c**). After 2 h, PZQ exhibited a concentration-dependent increase in rate of movement and frequency, a phenotype consistent with PZQ’s known spastic paralytic effect ^27^. These measurements peaked at 5 µM (EC_50_ = 389 nM and 196 nM, respectively). Note the shift between frequency and rate, which we attribute to the greater sensitivity of the frequency mode for these shorter somules (see also **Fig. 3**). Also, PZQ showed a concentration-dependent shortening of the parasite that reached a minimum length of 89 ± 3.6 µm at 82 nM (compared to 141 µm for DMSO; ES = -1.7) after which a partial lengthening occurred. The shortening was observed at a 5-fold lower concentration than that needed to increase rate and frequency (EC_50_ = 27 nM for first inflection of the PZQ static-length bell curve and EC_50 =_ 147 nM for frequency-length). The difference in concentrations may point to more than one molecular target /mechanism of action for PZQ.

The time-dependence of the shortening effect by PZQ was also different from that of the spasticity. Although both were observed at 2 h (see above), the shortening effect disappeared by the 48 h time point (**Supplementary Fig. 4f**), whereas spasticity remained unchanged (**Fig. 3f**). These data highlight the *ephemeral* or *transient* nature of some phenotypic responses, a concept that has not yet been considered in anthelmintic screening. Overall, the imaging methodology, as interrogated through SchistoView, allows for the orthogonal identification and quantification of individual concentration- and time-dependent changes.

For the other four members of the seven-drug test set, phenotypic effects at 2 h preceeded degeneracy/death that were recorded at later time points (**Fig. 4**). For example, two known anti-schistosomal agents simvastatin ^18^ and K11777 ^28^ induced gradual increases in degeneracy (simvastatin EC_50_ = 1 µM at 24 h; K11777 EC_50_ = 20 μM at 48 h). Degeneracy caused by staurosporine was apparent by 24 h and extended across the entire concentration range (**Fig. 4**), consistent with this inhibitor’s high affinity for multiple kinases. Other changes included increased area (65% larger than clear worms in DMSO and13% larger than degenerate worms in DMSO) and increased median diameter (51% larger than clear worms in DMSO and 20% larger than degenerate worms in DMSO; Supplementary Fig.2**, row D**). Interestingly, within 2 h, sunitinib produced a gray to jet-black phenotype that was much darker than degenerate worms in DMSO (**Fig. 5d**). This static phenotype (density levels) was significant at lower concentrations than the change in rate. As with PZQ, the complexity of these changes may reflect the time- and concentration-dependent engagement of different targets^29^.

### Using multi-dimensional features for primary screening

In addition to phenotyping based on inspection of individual descriptors, the screening platform and the SchistoView repository are applicable to high-throughput screens using *d_M_*. We prepared 20 x 96-well plates with 40 somules/well that were incubated with DMSO (0.1%) or 10 µM compound from an in-house collection of 1,323 drugs approved for human use. Using the same sample preparation and imaging conditions as for the seven-drug set, plates were robotically handled without manual intervention. Screening proceeded at a maximum rate of one plate/37 min for four scan cycles. Images were automatically processed and analyzed, and data entered into the MySQL database. The screen generated 59,867,820 measurements for 553,492 segmented worms. The *d_M_* values, calculated from the combined static, rate and frequency data, were then extracted and plotted in **Fig. 6a**.

**Figure 6.**
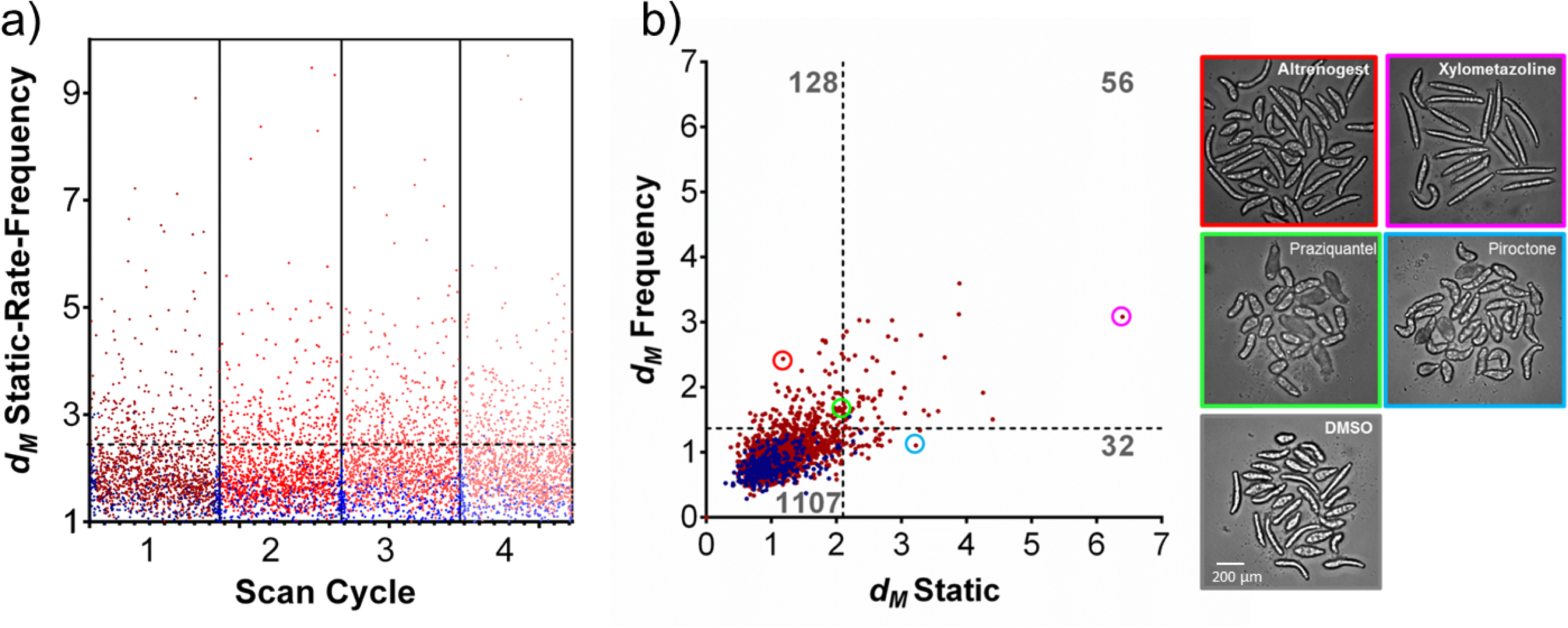
Scatterplot of data from a primary screen of 1,323 approved drugs. (**a**) Twenty plates were imaged over four scan cycles, each representing approximate intervals of 24 DMSO controls are shown in blue and the tested drugs (10 µM) are in red. *d_M_* values combine static, rate and frequency modes. The horizontal dashed line represents the *d_M_* value (2.47) that is 3 SD from the mean of the DMSO controls across the four scan cycles. This *d_M_* value differed by < 10% between individual scan cycles. The number of drugs with *d_M_* ≥ 2.47 are 237, 263, 326 and 309 compounds for scan cycles 1 through 4, respectively. (**b**) Scatter plot showing contributions of *d_M_* values based on static (x-axis) *vs.* frequency (y-axis) modes for the first scan cycle. The dashed lines represent the *d_M_* values that are 3 SD from the DMSO mean (2.1 for *d_M_* for static and 1.4 for *d_M_* for frequency). The number of drugs in each quadrant is indicated in grey font: 1,107 drugs were inactive; 32 drugs induced static phenotypes only, 128 induced only kinetic phenotypes and 56 compounds induced significant changes in both modes. The frames of the images to the right are color-matched with the highlighted compounds in the plot: note the remarkable range of phenotypes presented by this parasite.

Because *d_M_* does not have an inherent maximum value, the typical screening metric, Z’, cannot be calculated. However, ‘hits’ are usually picked based on standard deviations from the mean of the DMSO controls. For the data in **Fig. 6a** (combining static, rate and frequency), a *d_M_* value of 2.47 represents 3 SD from the mean of the DMSO-treated controls. Using this cutoff value, a total of 237, 263, 326 and 309 hits were identified for scan cycles 1 through 4, respectively. Notably, three drugs (PZQ, simvastatin and sunitinib) from the seven-drug test were also present in this screening set, and were identified as active, with *d_M_* values greater than 2.47.

The individual contributions of the static and frequency measurements to the identification of hits were illustrated by calculating a *d_M_* for each measurement in scan cycle 1 (**Fig. 6b**). Of the 216 total hits identified using these individual *d_M_* values, 128 (59%) of the active compounds differed from DMSO-treated worms only in frequency, 32 (15%) compounds differed only by static measurements and 56 (26%) showed a significant response by both static and frequency measures. Likewise, when considering rate *vs*. static *d_M_*, the results also point to the importance of motion-based descriptors. Together, 68% of active compounds differed from DMSO-treated worms based on changes in rate but were not significantly different from DMSO based on static features (**Supplementary Fig. 5**). Finally, 80 compounds had statistically significant changes in frequency but not rate and 116 compounds had statistically significant changes in rate but not frequency (**Supplementary Fig. 6**). In summary, quantifying changes in motion was critical for identifying the majority of the active compounds in the drug set.

Phenotypes among the 1,323 drug set were remarkably varied. For example, somules exposed to the hormone analog altrenogest displayed static features similar to those of DMSO controls (**Fig. 6b**), but exhibited a 144% increase in frequency-of-movement by length and a 147% increase in frequency-of-movement by area. Hence, altrenogest was ‘active’ based on the *d_M_* for frequency. By contrast, the antifungal piroctone only yielded changes in static features with a decreased variation in internal texture (63%) and an increase in form factor (110%, *i.e*., a more rounded phenotype) relative to DMSO controls. Somules treated with the adrenergic agonist xylometazoline were altered in both static and dynamic descriptors, *i.e*., a greater mean length of 211 µm *vs.* 130 µm for controls and a 160% increase in frequency of movement by length, respectively. Consistent with the data in **Fig. 3**, PZQ-treated somules were shorter (82%) with an increased form factor (140%) compared to DMSO controls (**Fig. 6b)**.

The data obtained for the 1,323-member drug set were compared with results from another drug screen that employed an observation-based scoring system (**Supplementary Data**) ^30^. Of the 235 compounds for which data were described ^30^, 143 were also screened by us. To compare the screens, we considered the third time point (closest to the 72 h time point used in ^30^) and defined ‘actives’ as those for which *d_M_* was >2.47 and/or degeneracy was >50%. Of the 143 compounds, 66 compounds were identified as active in both screens, 36 only by us and 11 only in ^30^; 30 compounds were inactive in both screens. Overall, there was a 67% agreement between the screens, *i.e*., considering both actives and inactives. Interestingly, the two screens differed the most in the category of compounds that we identified by the *d_M_* metric only: out of 30 actives in this category, only 12 were also identified in ^30^ (40% concordance). Thus, the present screen extends our ability to identify compounds by quantifying live phenotypic changes in the parasite.

## DISCUSSION

The screening platform described here comprises an integrated suite of solutions that solve the bottlenecks hampering drug discovery for global disease pathogens like the schistosome. Issues addressed include: a) producing and dispensing 10^4^-10^5^ parasites/week to enable automated screening; b) developing a robust image-collection and segmentation protocol and c) designing a system – SchistoView – to store, visualize, query and explore the multivariate data. Using the method, we captured the complexity of the schistosome’s response to seven drugs and identified previously unreported screening hits from a drug library. The data highlight the importance of quantifying changes in motion on the seconds timescale. To our knowledge, this scale and quantitative depth have not been achieved before for schistosomes or other parasitic helminths and the method is, in principle, adaptable to other organisms.

Somules are difficult to image due to their (variability in) movement and because they have a low contrast in bright field. We solved the image-collection challenge using round-bottom wells to constrain worms into one visual field and thus limit translation. We then addressed their low contrast by focusing slightly below the worm to enhance its outline. From there, we observed two basic classes of worms – clear and opaque (degenerate), and optimized segmentation protocols for each. The resulting segmentation accuracy (precision of 88% and a recall of 95%) is an improvement on a previous report that employed bright-field analysis for HTS screening (24.5±7% segmentation accuracy) where touching somules could not be evaluated ^19^. Also, our imaging approach economizes on the number of parasites needed by 3-4 fold and measures *how* worms move rather than a simple classification of *whether* movement has occurred ^19^. Finally, our methodology provides a solution to the critical issue of segmentation of touching objects in the analysis of bright-field images generally ^31–33^.

Our live imaging platform can be incorporated into a drug discovery pipeline upstream of *ex vivo* phenotypic screens of adult schistosomes and rodent models of infection^15^. Recent advances in the image-based quantification of adult schistosome motility^34, 35^ have demonstrated the ability to quantify motion and could mesh seamlessly with the workflow described here. The platform will also complement other advances relating to schistosome biology, including gene expression profiling ^36^, metabolomics ^37^, and CRISPR/Cas9 ^38^, that together will improve our ability to holistically quantify this globally important parasite’s responses to a range of drug-induced, environmental and developmental phenotypes. To facilitate such discoveries, the database and SchistoView interface are available online.

## METHODS

### Compounds

K11777 was synthesized via a contract research organization ^28^; simvastatin (S1792) was purchased from Selleckchem. PZQ (racemic; P4668), metrifonate (45698) and L-imipramine (I0899, HCl) were purchased from Sigma Aldrich. Sunitinib (S-8803 as the malate salt) and staurosporine (S-9300 as the free base) were purchased from LC laboratories. The library of approved drugs included 1,129 compounds from the Microsource Discovery Systems Drug Collection and an additional 194 compounds donated by Iconix or purchased from commercial vendors. The set included drugs approved by the FDA (85%) and by the analogous European and Japanese agencies (15%).

### Ethics Statement

Vertebrate animal maintenance and use were performed in accordance with UCSF’s Institutional Animal Care and Use Committee protocol AN086607.

### Somule preparation and plating

The *S. mansoni* life-cycle (NMRI isolate) is maintained by intraperitoneal injections of up to 600 infective larvae (cercariae) into 4-6 week-old, female Golden Syrian hamsters. Eggs are harvested from hamster livers six weeks later to generate miracidia which are then used to infect the *Biomphalaria glabrata* (NMRI strain) snail vector. The platform design (**Fig. 1**) involves the intensive propagation of snails to produce 10^4^ – 10^5^ infective larvae (cercariae) per week which are sufficient for up to twenty-five 96-well assay plates. Cercariae are then mechanically converted ^15^ into the post-infective schistosomula (somules, ∼ 200 x 100 µm) that are relevant to infection in humans. These somules are used within 2 h of their transformation from cercariae.

Each well of a 96-well u-bottom polystyrene assay plate (Corning, Costar 3799) was pre-wetted with 200 µL ddH20 to prevent the formation of bubbles at the well surface. After aspirating the ddH20, each well received an average of 40 somules in 200 µL Basch medium ^39^ supplemented with 4% heat-inactivated FBS, 100 U/ml penicillin and 100 mg/ml streptomycin. Somules (40,000) were suspended in 200 mL of medium using a magnetic tumble stirrer (V&P Scientific, VP 710C1-ALPFX) rotating at 45 rpm and then dispensed with a 96-channel pipette head (Beckman Coulter Biomek FXp) loaded with sterile 165 µL wide bore filter tips (Axygen, FXF-165-WB-R-S). Eight 96w plates could be prepared per tumbler volume of 200 mL (including a 40 mL dead volume). All assay plates were dispensed in less than 5 min to minimize tumbling damage to the parasite. Compound in neat DMSO was added to the well at the required final assay concentration in 0.1% DMSO using a 96-channel pin tool fitted with 200 nL slotted hydrophobic pins (V&P Scientific, AFIXFX96FP3). Compound was added while shaking the plate at 1,000 rpm in a 0.5 mm radius using a Teleshake 1536 (Variomag) which dispersed the compound from the pin into the surrounding media.

### Robotic handling, imaging and analysis

The Momentum 2.0 automation scheduler moved each assay plate from the automated tissue culture incubator (Thermofisher C2, 37⁰C, 5% CO2) to the barcode reader, then to the automated microscope (GE IN Cell Analyzer 2000), and back to the tissue culture incubator. Each iteration took approximately 35 min.

A high-content imager (InCell Analyzer 2000; GE Healthcare) was used to collect 20 seconds of time-lapse images of parasites under one field of view with a 10x objective. The 4 megapixel CCD sensor was binned 4x4 and the bright-field/DAPI channel was set to a 3 ms exposure. The focal plane was offset 40 µm from the bottom of the well to thicken the edge (surface) of the worm. The “sit-and-stare” time-lapse schedule began with a 3.5 sec delay to allow time for auto-focusing followed by 30 image acquisitions 0.66 sec apart.

Images were then analyzed and segmented as described in **Supplementary Information, Extended Methods**. Features were extracted from the optimal mask chosen from multiple segmentation attempts for each worm and were stored in a custom MYSQL database for visualization in SchistoView.

### Statistics and reproducibility

Microsoft VBA was used to perform all statistical analysis. The method monitors each worm as a separate object. *Glass* effect size ^20, 21^ and *Mahalanobis d_M_* ^22^ for the seven drug set were were based on the average values from 160 somules (four experimental wells, each with approximately 40 worms). Error bars shown in **Fig. 5** represent standard deviations in effect size from four wells, each containing approximately 40 worm objects. Dotted lines above and below baseline represent three standard deviations from the mean of the DMSO-treated wells. Data shown are representative of at least three independent experiments. *Glass* effect size and *Mahalanobis d_M_* for the 1,323-compound drug set were calculated from approximately 40 worms/test compound in one well. Dotted lines in **Fig. 6** represent three standard deviations from the mean of the DMSO-treated wells.

### Data Availability

**Supplementary materials** include **Supplementary Information** (Supplementary Table 1, Supplementary figures, and Extended Methods detailing data collection, segmentation, and database protocols) and **Supplementary Data** (1,323-drug screen, hits and non-hits). The data of the seven drug dose response screen and the 1,323-compound drug screen are available through SchistoView ^23^. Any remaining data can be obtained from the corresponding author upon reasonable request.

## Supporting information

1323 drug comparison

## Acknowledgements

We thank the following funding sources: NIH AI146719, NIH AI089896 and NSF IIS-1817239.

## Author Information

### Contributions

C.R.C., B.M.S. and S.C. performed the screens. S.C. designed the scripts for image analysis. S.C., J.D. and R.S developed SchistoView and all authors contributed to its design. S.C., C.R.C., M.R.A. and R.S interpreted the data. The paper was written by S.C., C.R.C., M.R.A. and R.S.

## Supplementary Information

**Supplementary Table 1.**
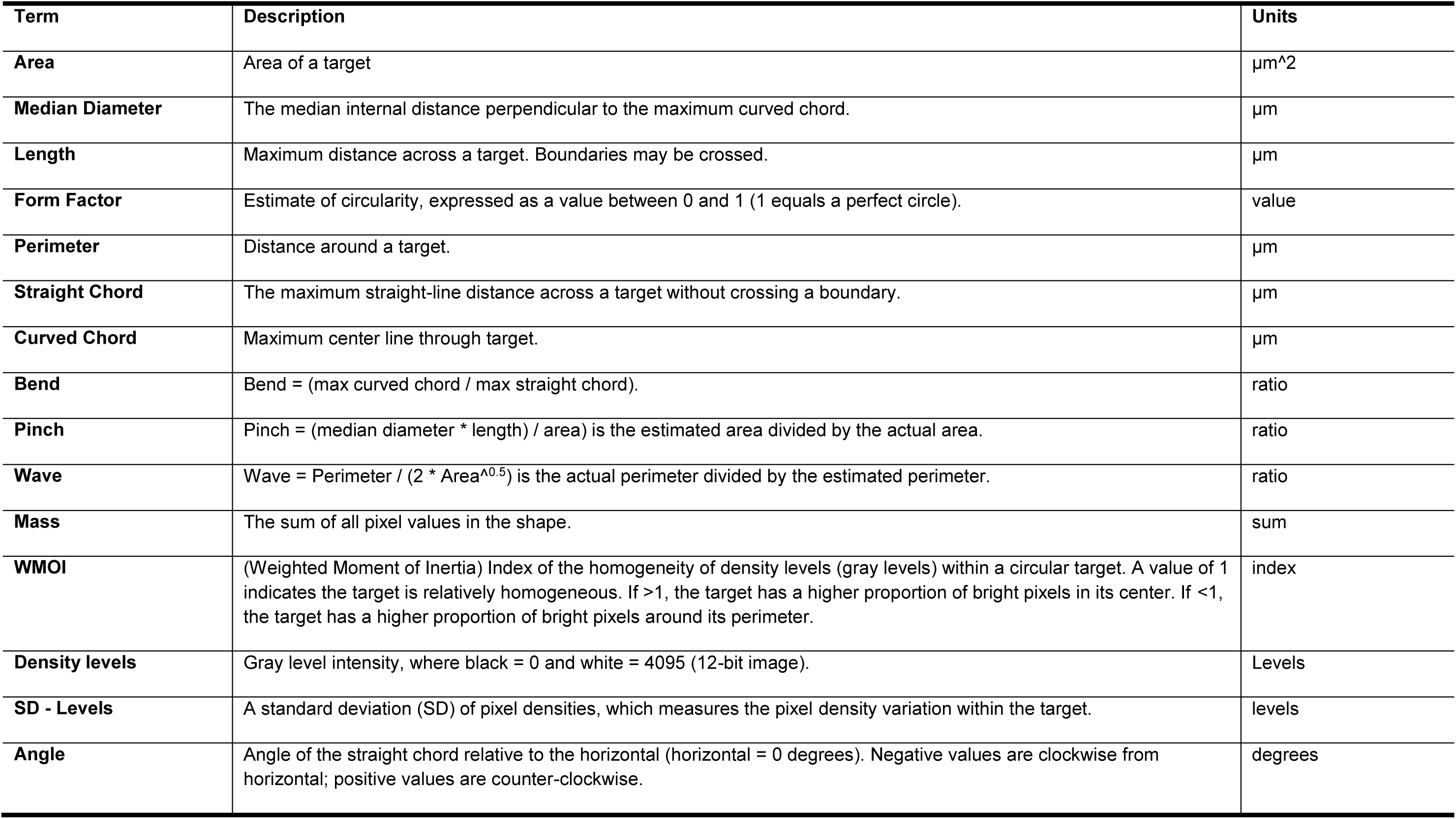
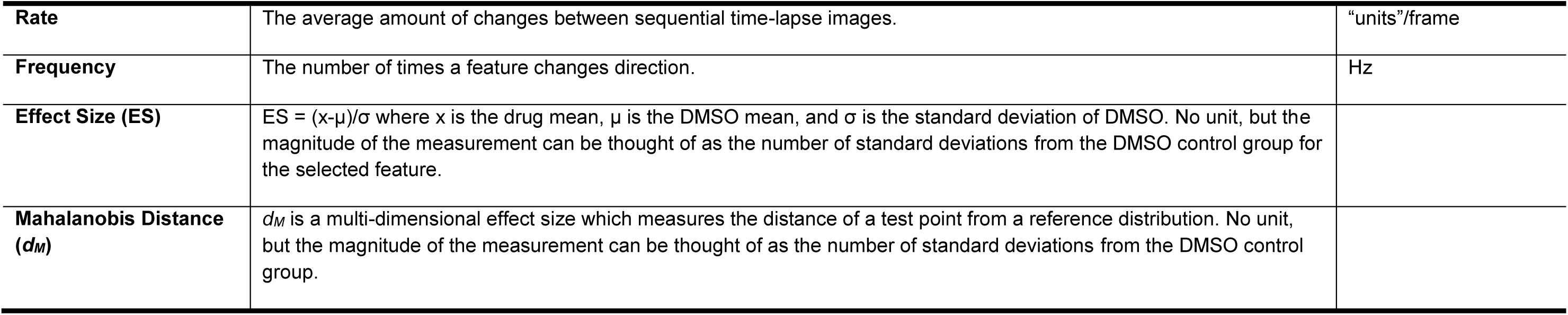
Glossary of Terms.

| Term | Description | Units |
| --- | --- | --- |
| **Area** | Area of a target | $\mu\text{m}^2$ |
| **Median Diameter** | The median internal distance perpendicular to the maximum curved chord. | $\mu\text{m}$ |
| **Length** | Maximum distance across a target. Boundaries may be crossed. | $\mu\text{m}$ |
| **Form Factor** | Estimate of circularity, expressed as a value between 0 and 1 (1 equals a perfect circle). | value |
| **Perimeter** | Distance around a target. | $\mu\text{m}$ |
| **Straight Chord** | The maximum straight-line distance across a target without crossing a boundary. | $\mu\text{m}$ |
| **Curved Chord** | Maximum center line through target. | $\mu\text{m}$ |
| **Bend** | $\text{Bend} = (\text{max curved chord} / \text{max straight chord})$. | ratio |
| **Pinch** | $\text{Pinch} = (\text{median diameter} * \text{length}) / \text{area}$ is the estimated area divided by the actual area. | ratio |
| **Wave** | $\text{Wave} = \text{Perimeter} / (2 * \text{Area}^{0.5})$ is the actual perimeter divided by the estimated perimeter. | ratio |
| **Mass** | The sum of all pixel values in the shape. | sum |
| **WMOI** | (Weighted Moment of Inertia) Index of the homogeneity of density levels (gray levels) within a circular target. A value of 1 indicates the target is relatively homogeneous. If $>1$, the target has a higher proportion of bright pixels in its center. If $<1$, the target has a higher proportion of bright pixels around its perimeter. | index |
| **Density levels** | Gray level intensity, where black = 0 and white = 4095 (12-bit image). | Levels |
| **SD - Levels** | A standard deviation (SD) of pixel densities, which measures the pixel density variation within the target. | levels |
| **Angle** | Angle of the straight chord relative to the horizontal (horizontal = 0 degrees). Negative values are clockwise from horizontal; positive values are counter-clockwise. | degrees |
| **Rate** | The average amount of changes between sequential time-lapse images. | “units”/frame |
| **Frequency** | The number of times a feature changes direction. | Hz |
| **Effect Size (ES)** | ES = $(x-\mu)/\sigma$ where $x$ is the drug mean, $\mu$ is the DMSO mean, and $\sigma$ is the standard deviation of DMSO. No unit, but the magnitude of the measurement can be thought of as the number of standard deviations from the DMSO control group for the selected feature. | |
| **Mahalanobis Distance ($d_M$)** | $d_M$ is a multi-dimensional effect size which measures the distance of a test point from a reference distribution. No unit, but the magnitude of the measurement can be thought of as the number of standard deviations from the DMSO control group. | |

**Supplementary Figure 1.**
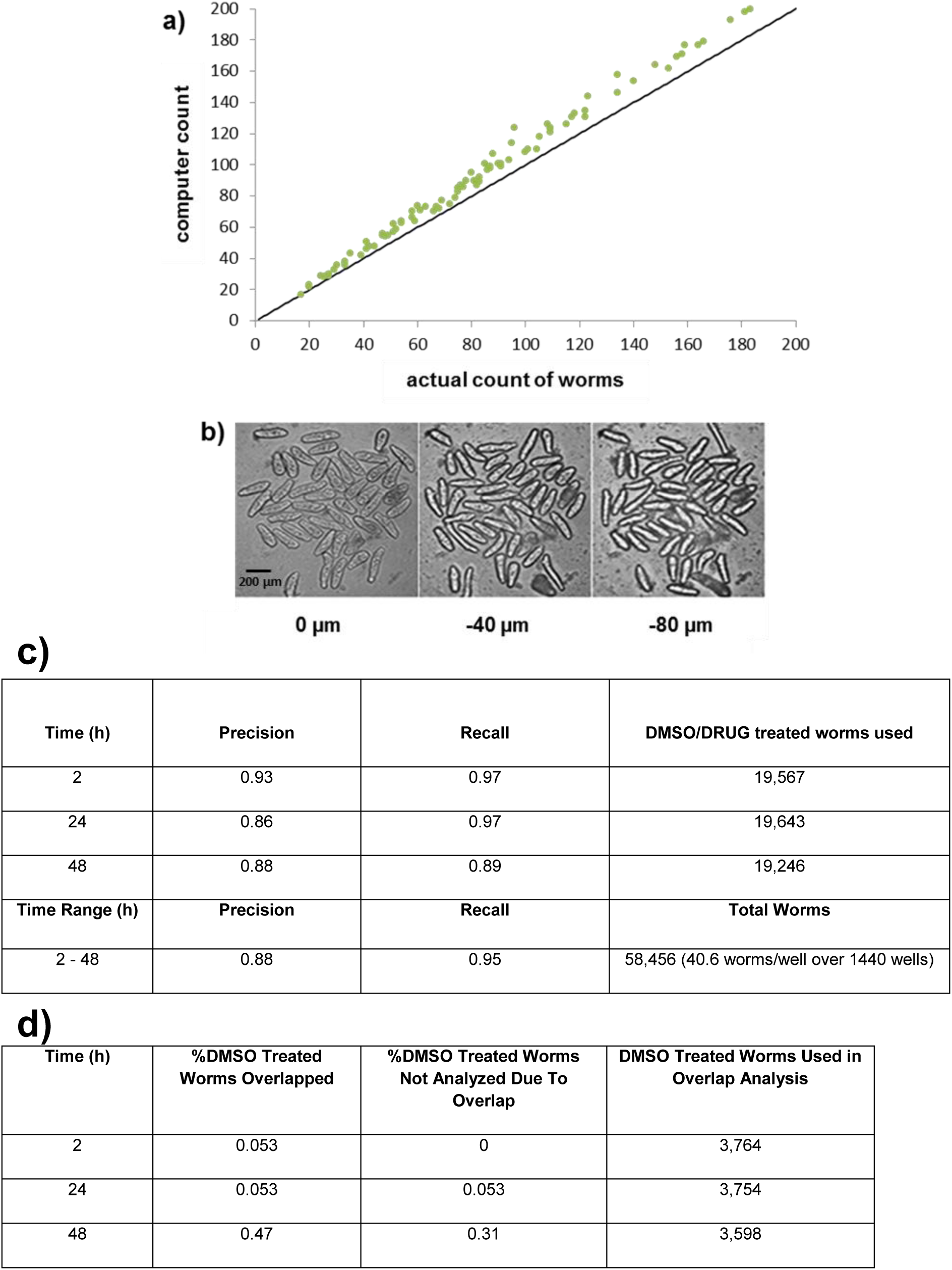
Optimizing parasite handling and segmentation. (a) Comparison of actual somules (counted by visual inspection) and the number of somules identified by the object classifier algorithm ‘computer count’ (see Extended Methods, below). Computational inclusion of non-worm objects with worm-like features leads to a systematic 10% increase in object count. (b) Images from a single sample well imaged at three focal planes (0, -40, and -80 µm from the outside bottom of the well). Lowering the focal plane improves the contrast of the somule outline (‘edge’); at -40 µm the appearance of the outline is improved while some of the internal texture detail is preserved. (c) Precision and recall were determined by manual inspection of the 58,456 somules that were screened in the seven-drug set. (d) Somule overlap frequency and data removal due to overlap events. The overlap increases from 0.05% to 0.3-0.47% between 24 and 48 h, potentially reflecting growth of the somules, increased degeneracy or increased motility.

**Supplementary Figure 2.**
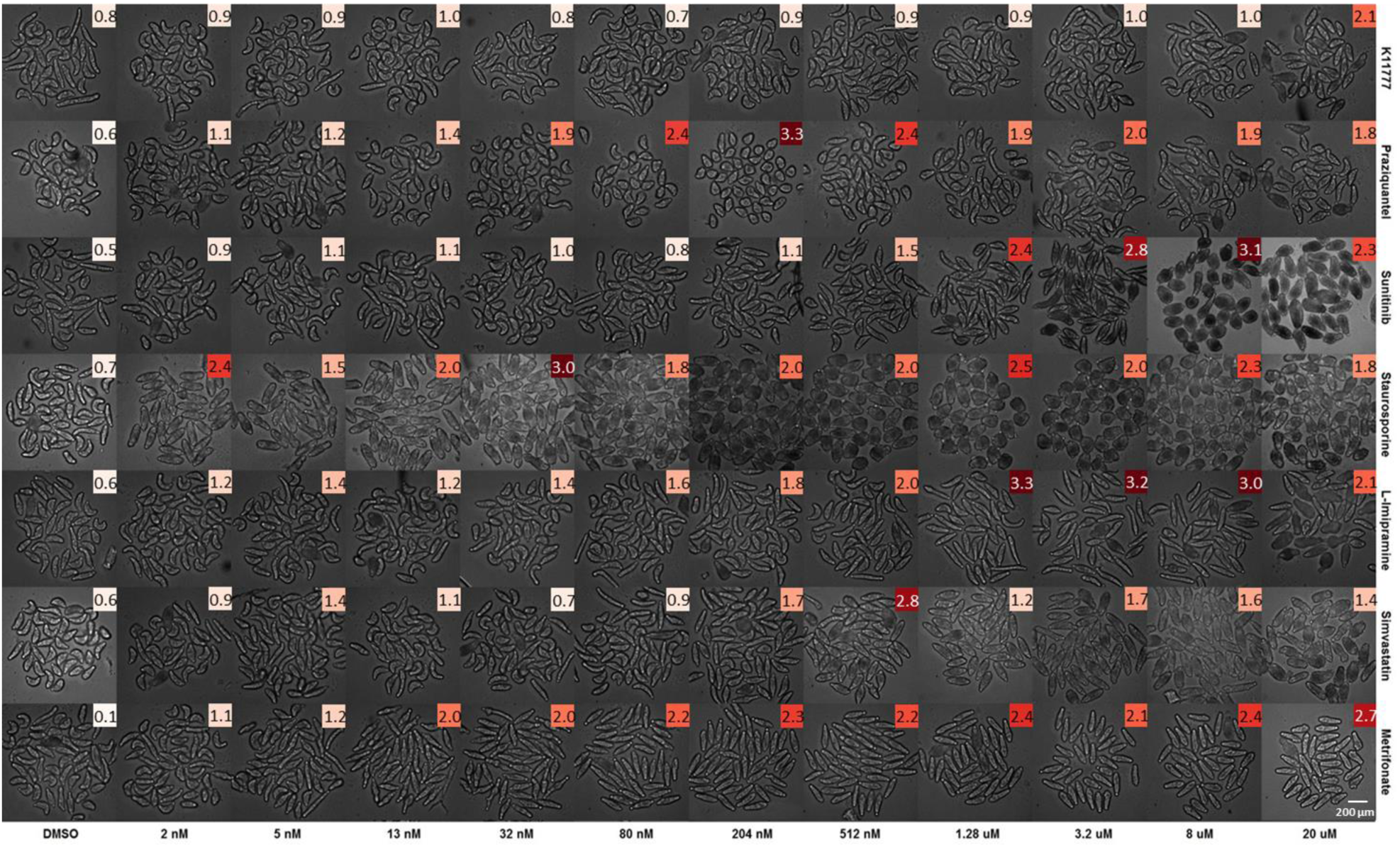
Montage of somule images 24 h after treatment with seven test drugs. Drug names are to the right and concentrations are at the bottom. Each image in the montage is labeled with the corresponding *d_M_* value and scaled color.

**Supplementary Figure 3.**
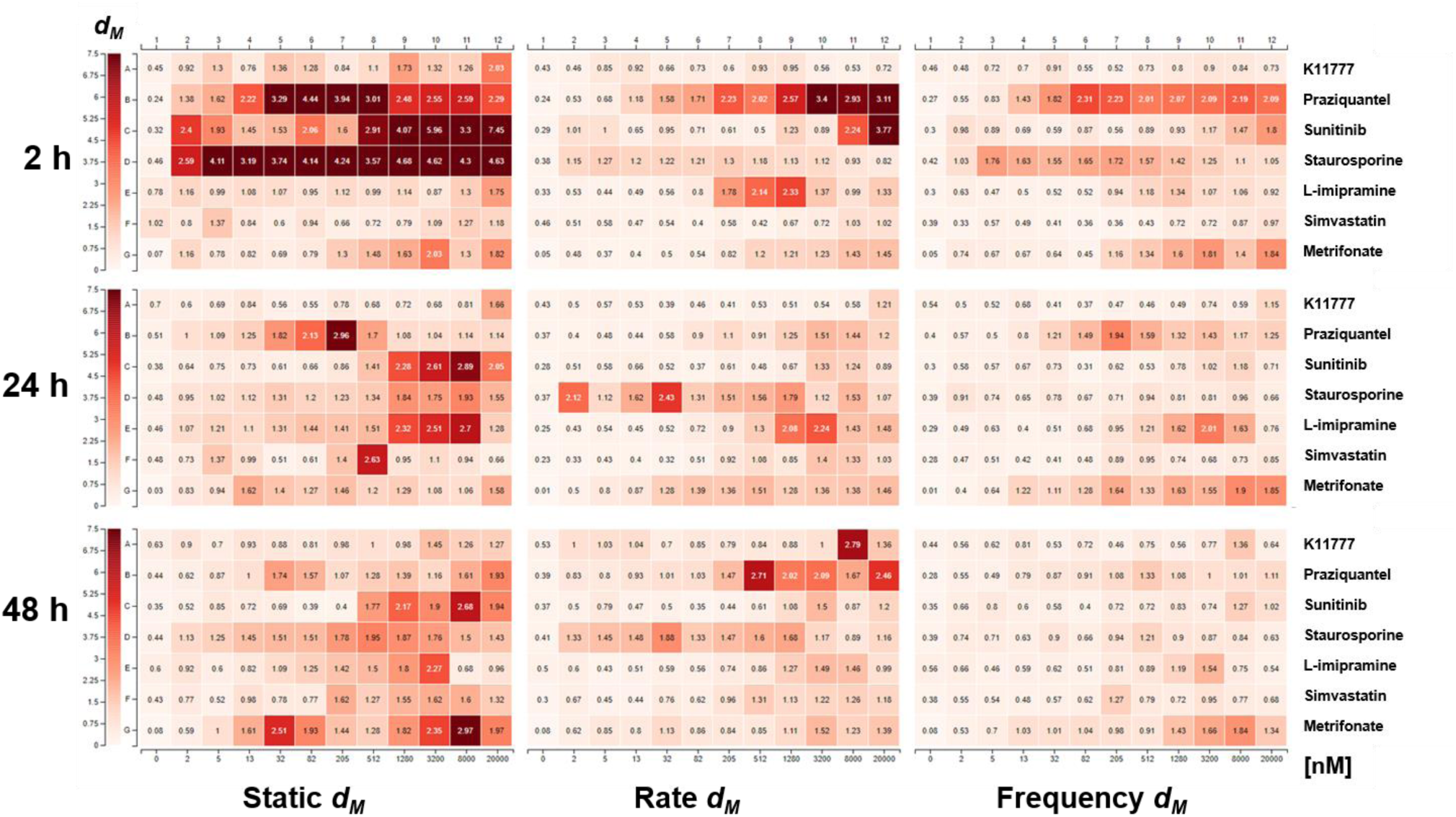
Differing sensitivities of *d_M_* measurements in the static, rate and frequency modes. *d_M_* values at 2, 24 or 48 h after treatment were measured using only static features (left panels), rate (center panels) or frequency for seven test drugs. Drugs were arrayed over an 11-point 2.5-fold dilution range from 2 nM to 20 μM. Values were determined from the aggregation of four wells per treatment. Note that the *d_M_* values shown do not necessarily smoothly change with increasing dose of drug. This complexity reflects the observation that multiple parameters show maximum changes at different concentrations. Each mode offers a differential sensitivity to measuring changes in the somule; *e.g*., the static *d_M_* for staurosporine across all concentrations after 2 h, the rate *d_M_* for imipramine (205 – 1280 nM) after 2 h and the frequency *d_M_* for metrifonate at 8 and 20 µM after 24 h.

**Supplementary Figure 4.**
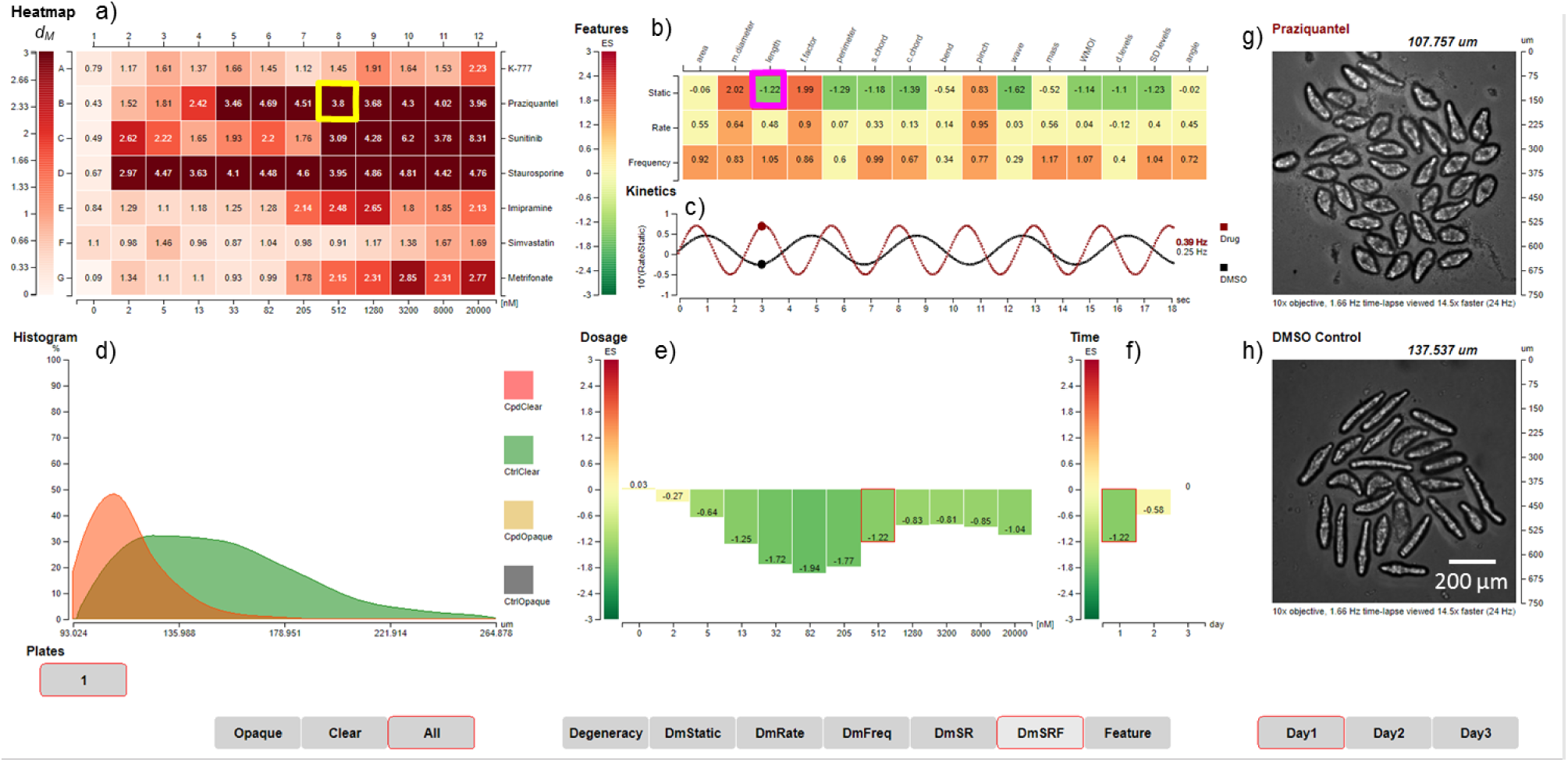
Screenshot of the SchistoView graphical user interface. The figure is analogous to **Fig 3**, but highlights the length of PZQ-treated somules whereas **Fig 3** shows the frequency of changes in length. Selected data are shown to illustrate the hierarchical approach to visualization. **(a)** Heat map of Mahalanobis distances (*d_M_*) for seven test drugs arrayed over an 11-point 2.5-fold dilution series from 2 nM in column 2 to 20 μM in column 12. Drugs, from top to bottom, are, K11777, PZQ, sunitinib, staurosporine, imipramine, simvastatin and metrifonate. DMSO controls are arrayed in column 1 and are shown as the average *d_M_* (0.77) for all DMSO controls. A *d_M_* of 1.61 is significantly different (3 SD) from control. Clicking on coordinate B8 (identified by the yellow square: 512 nM PZQ) populates panels (**b**) and (**g**) (see below). **(b)** Heat map showing the effect sizes (ES) for static, rate and frequency, after exposure to 512 nM PZQ for 2 h, *i.e*., the selected well from (**a**). Three sets of 15 features are arrayed in rows and columns, respectively. Clicking on the intersection of the length feature and static mode (magenta box) in (**b**) populates panels (**c**) through (**f**) and the underlying data. (**c**) Calculated waveforms defined by the range of length (amplitude) and frequency of length contraction (frequency). DMSO control worms are slower moving (lower frequency) than those treated with 512 nM PZQ (red line). (**d**) Histogram displaying the distribution of static length for DMSO control worms (green) and PZQ-treated worms (orange). (**e**) Bar graph depicting the ES for static length after PZQ treatment across 11 concentrations (second row in (**a**)). (**f**) Bar graph depicting the ES for static length in the 512 nM PZQ treatment across the three days of measurement. **(g)** First image from time-lapsed movie of the well highlighted in (**a**); in the live SchistoView, the 30-frame movie is looped. (**h**) as for (**g**) except for the DMSO control.

**Supplementary Figure 5.**
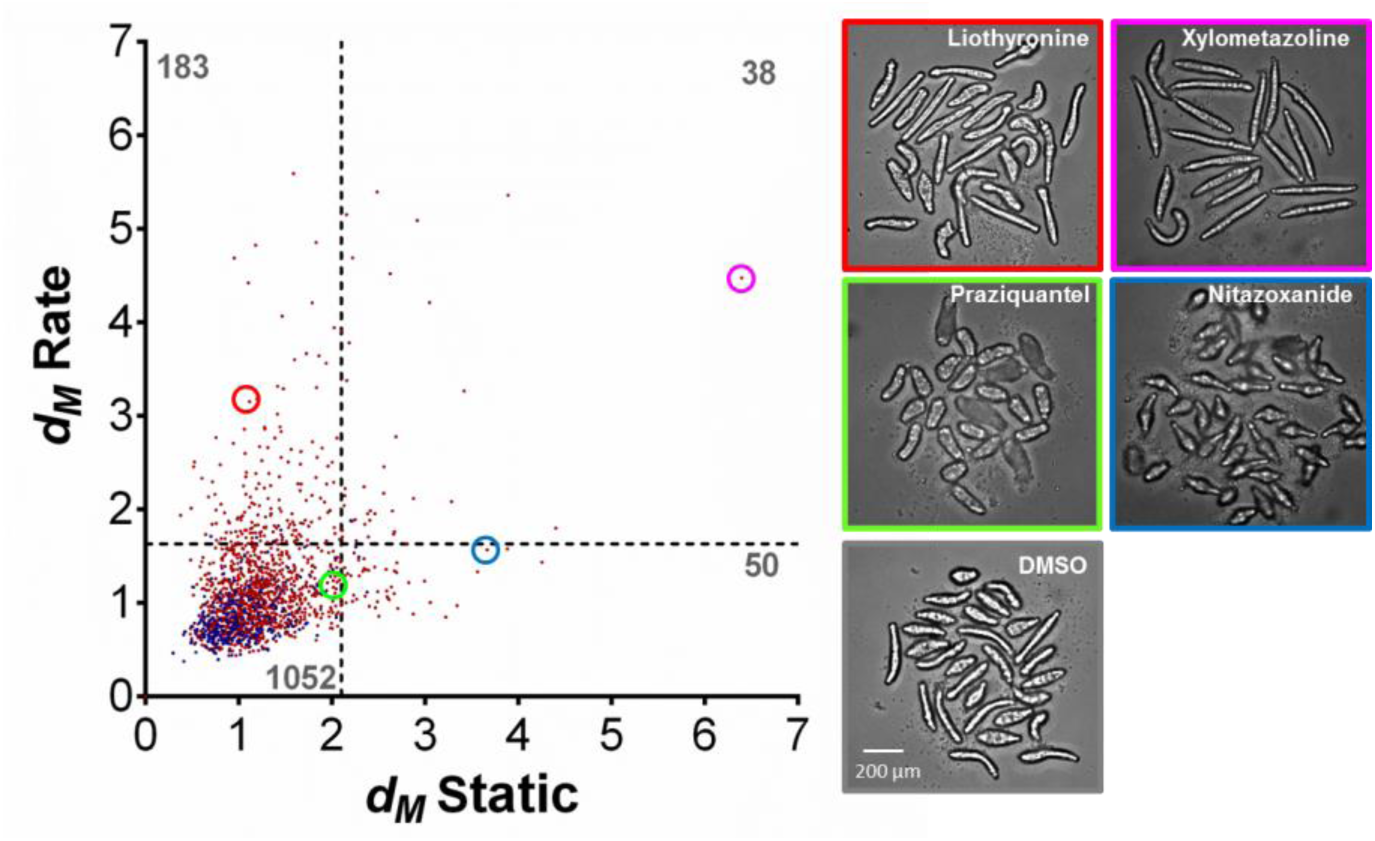
Scatterplot of *d_M(rate)v_s*. *d_M(static)_* for a primary screen of 1,323 approved drugs. Like Fig 6b in the main text, which shows a scatterplot of *d_M(frequency)_ vs. d_M(static)_*, the screen was performed at 10 µM. The data shown are from the first scan cycle approximately 24 h after the addition of drug. The dashed lines represent the *dM* values that are 3 SD from the DMSO mean (2.1 for *d_M(static)_* and 1.6 for *d_M(rate)_*). The number of drugs in each quadrant is indicated in dark grey for both static and rate modes: 1,052 drugs were inactive, 50 drugs induced static phenotypes only, 183 induced only kinetic phenotypes and 38 compounds induced changes in both modes. The frames of the images to the right are color-matched with the highlighted compounds in the plot: note the remarkable range of phenotypes presented by this parasite.

**Supplementary Figure 6.**
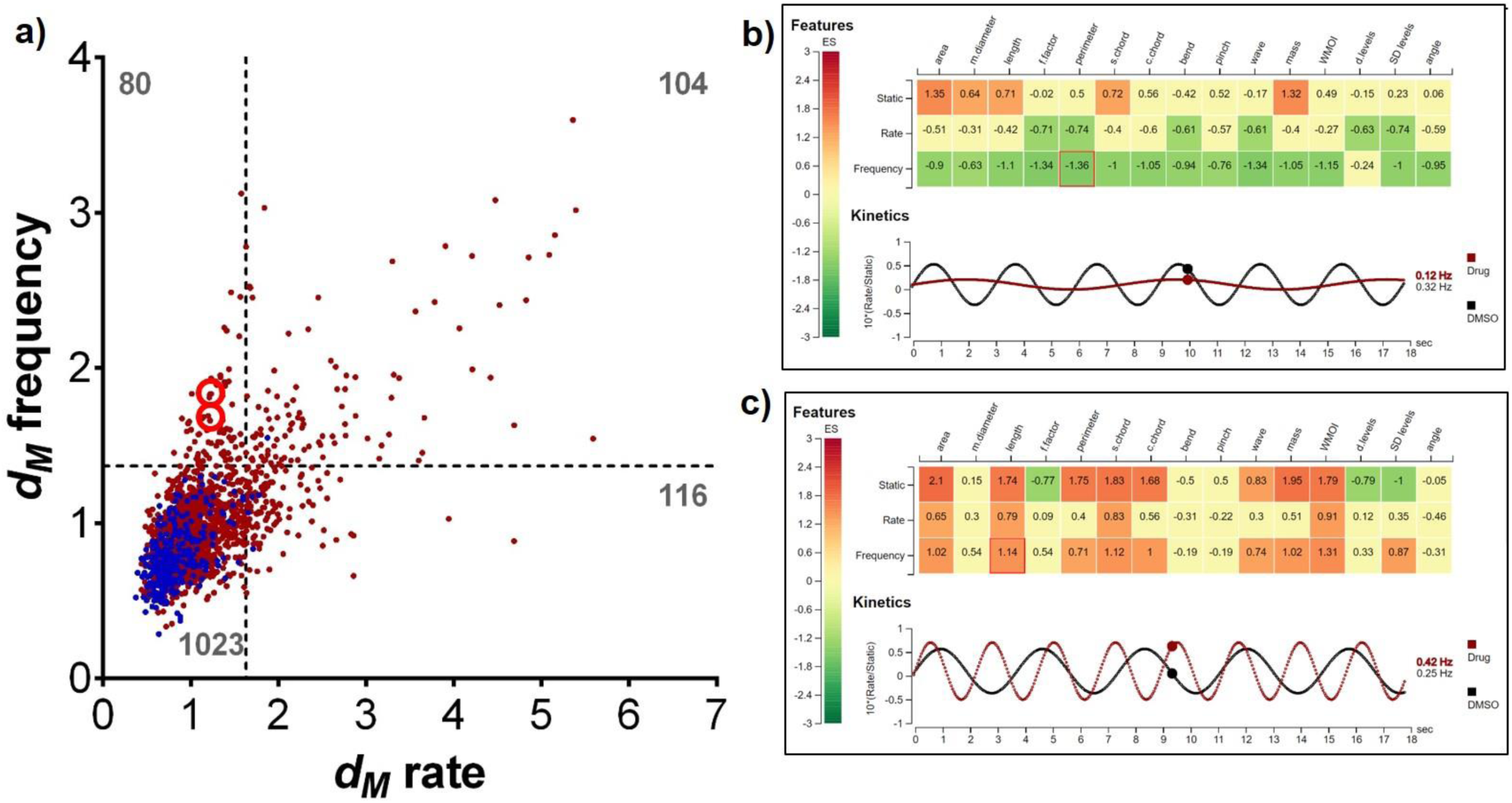
(**a**) Scatterplot of *d_M(frequency)_vs. d_M(rate)_* for a primary screen of 1,323 approved drugs. Like Fig. 6b in the main text which shows a scatterplot of *d_M(frequency)_ vs. d_M(static)_*, the screen was performed at 10 µM. The data shown are from the first scan cycle approximately 24 h after the addition of drug. The dashed lines represent the *dM* values that are 3 SD from the DMSO mean (1.6 for *dM* for rate and 1.4 for *dM* for frequency. The number of drugs in each quadrant is indicated in dark grey for both frequency and rate modes: 1,023 drugs were inactive, 116 drugs induced significant rate-based phenotypes only, 80 induced phenotypes associated with frequency only and 104 compounds induced changes in both modes. Two drugs, vilazodone and apomorphine, are marked with upper and lower red circles, respectively. (**b, c**) Examples of drugs that induce more changes in frequencies than changes in rates as shown by the table of Features (effect sizes) and Kinetics (red = drug; black = DMSO). Apomorphine (**b**) induces a hypomotile phenotype, whereas vilazodone (c) generates hypermotility. Images taken from SchistoView.

## Extended Methods

### Time-lapse Image Acquisition

Open the automation scheduler (Momentum 2.0) with instructions to move each assay plate from the automated tissue culture incubator (Thermofisher C2, 37⁰C, 5% CO_2_) to the barcode reader, then to the automated microscope (GE IN Cell Analyzer 2000), and then back to the tissue culture incubator. One cycle takes approximately 35 minutes or the time it takes to scan one assay plate in the automated microscope.

1. Place the 96-well round bottom polystyrene assay plate with lid (Costar 3799) into the nest of a GE IN Cell Analyzer 2000 (software version 3.0.0.43).
2. Open the acquisition protocol with the following settings (from XDCE):

a. Objective

i. Focal length = 20.0
ii. Id = 12111
iii. Lineartype = 7
iv. Magnification = 10
v. Numerical Aperture = 0.45
vi. Objective Name = 10X/0.45, Plan Apo, CFI/60
vii. Pixel height = 2.96
viii. Pixel width = 2.96
ix. Refractive index = 1.0
x. Unit = μm
b. Polychroic

i. QUAD1 (any polychroic will do)
c. CCD Camera

i. Size; height = 2048, width = 2048
ii. Flat Field Correction = False
iii. Binning value = 4 x 4
iv. Bias value = 144.21
v. Gain value = “”
d. Wavelength

i. Imaging mode = 2-D
ii. Excitation Filter = TL-Brightfield, 473 nM
iii. Emission Filter = DAPI, 455 nM
iv. Exposure time = 3 ms
v. HWAFOffset = 0 μm
vi. FocusOffset = -50 μm
e. Software Auto Focus = False
f. Laser Auto Focus = True
g. Plate dimensions as entered in software

i. Columns = 12, Rows = 8
ii. Plate height = 14.16 mm
iii. Bottom thickness = 1310 μm
iv. Bottom interface = plastic
v. Bottom height = 2.15 mm
vi. Well volume = 360 uL
vii. Well parameters = Round, Size = 6.35 mm
viii. Top Left Well Center Offset ; horizontal = 13.8 mm, vertical = 11.7 mm
ix. Well Spacing; horizontal = 9.0 mm, vertical = 9.0 mm
h. Plate heater use = false
i. Plate map, acquisition mode

i. Layout; columns = 1, rows = 1
ii. Direction value = horizontal snake
j. Time schedule = enabled

i. Incubate between time points = false
ii. Mode = spit and stare
iii. Time points in ms = 0, 3740, 4340, 4940, 5540, 6140, 6740, 7340, 7940, 8540, 9140, 9740, 10340, 10940, 11540, 12140, 12740, 13340, 13940, 14540, 15140, 15740, 16340, 16940, 17540, 18140, 18740, 19340, 19940, 20540, 21140

1. This first time point = 0 is the point at which the autofocus operation occurs. To allow time for this operation to complete, the next time point occurs at 3740 ms and continues at a rate of 600 ms between time lapse images. In total, there were 1 autofocus image and 30 time lapse images.
3. Write the image stack to an E-SATA connected storage device (preferably RAID) capable of storing 1.44 GB per assay plate per day.

The exposure time of 3 ms (step 2.d.iv) is the shortest exposure time the GE IN Cell Analyzer 2000 will accept. With the CCD camera binning value set to 4 x 4 (step 2.c.iii), some of the pixels in the image may be saturated (>= 4095). The binning value of 4 x 4 allowed for the fastest time lapse acquisition of 600 milliseconds since a smaller array is faster to readout. The possibility of saturated pixels was accepted in return for a faster frame rate.

### Image Segmentation (*Protocol: Schisto94*)

Open IN Cell Developer Toolbox 1.9 which can be found in IN Cell Investigator 1.6, a suite of software that comes with the GE IN Cell Analyzer 2000 automated microscope. From the “analysis” tab, select “batch analysis manager…” and “add…”, select the “Schisto94_ba” protocol, and image folders for analysis. Then select “Run batch analysis…”

There are three main workflows: clear body outliner, tegument outliner, dark body outliner. These workflows are imaging preprocessing macros which transform the image prior to segmentation. Each workflow produces 4 target types: threshold 1, threshold 2, threshold 3, and a merge of the results from threshold 1 to 3. Each target set records 17 features: area, x, y, diameter, length, form factor, perimeter, straight chord, curved chord, bend (a user defined feature), pinch (a user defined feature), wave (a user defined feature), mass, weighted moment of inertia, density-levels, sd-levels, and angle. The data is recorded at the cell level, or in this case, per organism.

The purpose of this segmentation is to cast a large net around a large variety of objects where up to 50% could be artefactual (i.e. inter-organism objects, intra-organism objects). This inefficiency is by design in order to lower the false negative rate in segmentation. The false positives (artifacts) are detected and removed in a subsequent data processing step external to IN Cell Developer 1.9

Before any of the workflows are run, the raw image is processed with a custom built flat field correction (FFC) image preprocessing macro. All subsequent processing and analysis will be based on the flat field corrected image. (The flat field correction is the first step in the Macro: *schisto94_n_master*.)

1. **Flat Field Correction** (Macro: *Schisto94_a_FFC*) (**Figure 1**). Divide the raw image by an estimate of the background of each image to produce a flat field correction. The steps below outline how an estimate was generated. (Code enumerated in roman numerals) (8 operations per image)

a. Load raw image.
b. Transform result from step “1.a” with transform filter = median, kernel size = 5
c. Transform result from step “1.b” with transform filter = local arithmetic

i. kernel = 99;
ii. src = lAve;
d. Apply a transform point operation = arithmetic (two src) where source1 = result from step “1.b” and source2 = result from step “1.c”.

i. Sel(abs(src1-src2)/src2&lt;0.025,src1,0);
e. Transform result from step “1.d” with transform filter = max, kernel size = 33
f. Transform result from step “1.e” with transform filter = local arithmetic

i. kernel = 51;
ii. src = lAve;
g. Transform result from step “1.f” with transform filter = local arithmetic

i. kernel = 67;
ii. src = lAve;
h. Apply a transform point operation = arithmetic (two src) where source1 = result from step “1.a” and source2 = result from step “1.g”.

i. (src1/src2)*2048; The round wells of the 96-well assay plate produce varying backgrounds due to the position of the well in the plate, the artefacts in the plastic, and changes in illumination due to the grayness of the sample. Therefore, an *in situ* background estimate was used. Flat field correction and centering of the pixel values in each image allows for the universal application of object detection thresholds based on fixed values.

**Figure 1.**
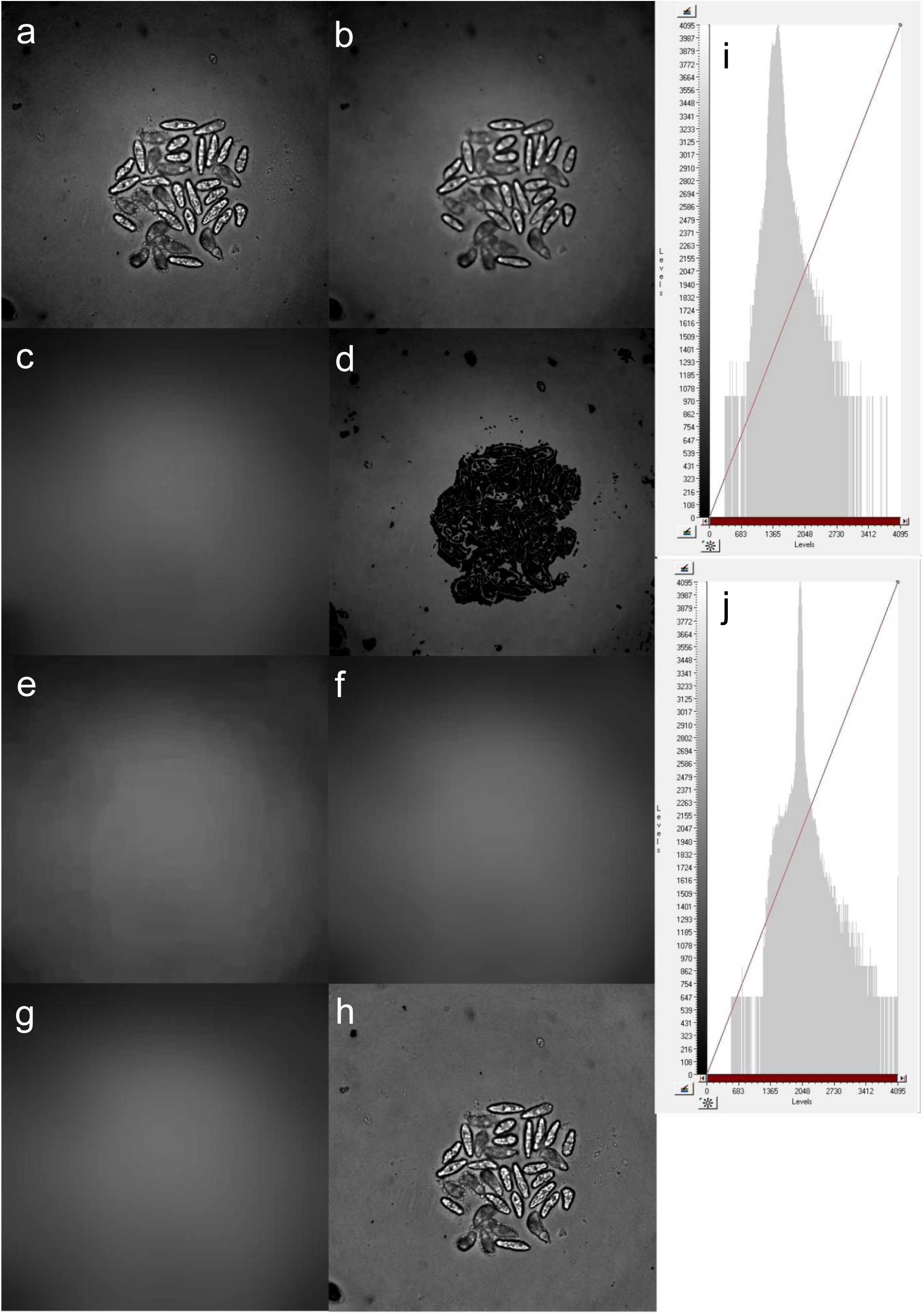
Flat Field Correction using an *in situ* background estimate. a) raw image, b) median transform of raw image (light smoothing), c) local arithmetic transform of raw image (heavy smoothing), transform point operation which sets pixel in panel "b" to zero if pixel is > 2.5% different from the same pixel in panel "c", e) max transform promotes background pixels between and around zeroed pixels in panel "d", f) local arithmetic transform smooths result from panel "e", g) local arithmetic transform smooths result from panel "f", h) transform point operation which divides the raw image by the background estimate in panel "g" and then multiplies the result by 2048 to center the image pixel values within the 12-bit range, i) histogram of the raw image, j) histogram of the flat field corrected image. The next macro is called “clear body outliner” because the outlines produced are based on image thresholding which target the white translucent area of the organism.
2. **Clear Body Outliner Preprocessing** (Macro: *schisto94_n_master*) (Figure 2) (53 operations per image)

Transform FFC image with transform filter = local arithmetic (Macro: *schisto94_n_1950)*:

a. sel(src<1950,4095,0);
b. Transform result from step “2.a” with transform filter = sieve (binary), retain objects > 500 μm^2
c. Transform result from step “2.b” with transform filter = closing (binary), kernel = 3
d. Transform result from step “2.c” with transform filter = thinning, passes = 3
e. Transform result from step “2.d” with transform filter = pruning, passes = 3
f. Transform result from step “2.e” with transform filter = inversion
g. Transform result from step “2.f” with transform filter = sieve (binary), retain objects > 2600 μm^2
h. Repeat steps “2.a” through “2.g” with following code for step “2.a”, (Macro:

i. schisto94_n_1800):
ii. sel(src<1800,4095,0);
i. Repeat steps “2.a” through “2.g” with following code for step “2.a”, (Macro: *schisto94_n_1700*):

i. sel(src<1700,4095,0);
j. Reset a channel with transform filter = local arithmetic

i. src = 0;
k. Transform result from “Macro: *schisto94_n_1950*” with transform filter = dilation (binary), kernel = 3
l. Apply a transform point operation = arithmetic (two src) where source1 = result from threshold “Macro: schisto94*_n_1950*” and source2 = result from step “2.k”.

i. src2-src1;
m. Apply a transform point operation = arithmetic (two src) where source1 = result from step “2.j” and source2 = result from step “2.l”. The result is added to the channel used in step “j”.

i. i. src2+src1;
n. Repeat steps “2.k” through “2.m” with result from “Macro: *schisto94_n_1800*”.
o. Repeat steps “2.k” through “2.m” with result from “Macro: schisto94_n_1700”.
p. Transform merge result (channel in step “j”) with transform filter = inversion
q. Transform result from step “2.p” with transform filter = sieve (binary), retain objects > 100 μm^2
Transform result from step “2.q” with transform filter = inversion
s. Transform result from step “2.r” with transform filter = erosion, kernel = 3
t. Transform result from step “2.s” with transform filter = sieve (binary), retain objects > 100 μm^2
u. Transform result from step “2.t” with transform filter = pruning, passes = 3
v. Apply a transform point operation = arithmetic (two src) where source1 = result from step “2.q” and source2 = result from step “2.u”.

i. src2+src1;
w. Apply a transform point operation = arithmetic (two src) where source1 = result from step “2.v” and source2 = result from step “1.h”.

i. sel(src1==4095,src2,0); Steps “2.a” through “2.i” generate the preliminary target data for threshold1 to threshold3. (Figure 2A) The refinement of this preliminary data and consolidation into a fourth target begins with step “2.j”. (Figure 2B) The fourth target is based on a merge of the data from the three thresholds. The resulting outline will have extra objects produced by the intersections. The extra objects are filtered to reduce complexity of the merged result and this process starts with step “2.p”. (Figure 3B) In addition, the merge result represents all possible objects all clear body target sets and these results are used to patch holes in the results from threshold 1 to 3. A complete set of overlapping objects is a requirement for target linking.

**Figure 2A.**
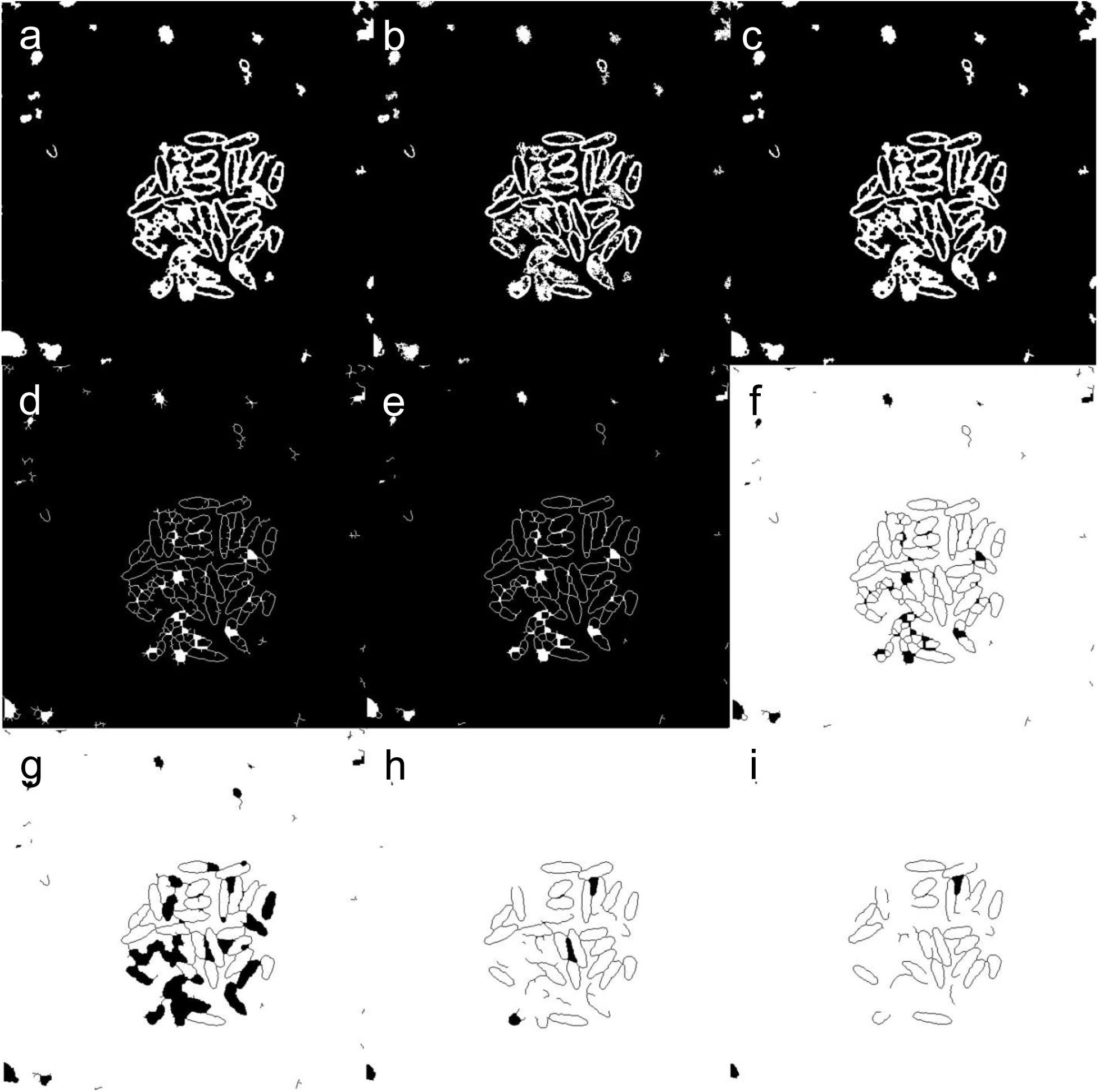
Clear body workflow for thresholds 1 to 3. Panels “a” through “g” describe the clear body workflow for threshold1 (“Macro: *schisto94_n_1950*”). a) binarization of the flat field corrected image, b) sieve to remove small objects, c) closing to close gaps, d) thinning to reduce outline to a skeleton, e) pruning to prune back branches, f) inversion, g) sieve to retain large objects (this is the threshold1 “Macro: *schisto94_n_1950*” result), h) threshold2 “Macro: *schisto94_n_1800*” result, i) threshold3 “Macro: *schisto94_n_1700*” result.

**Figure 2B.**
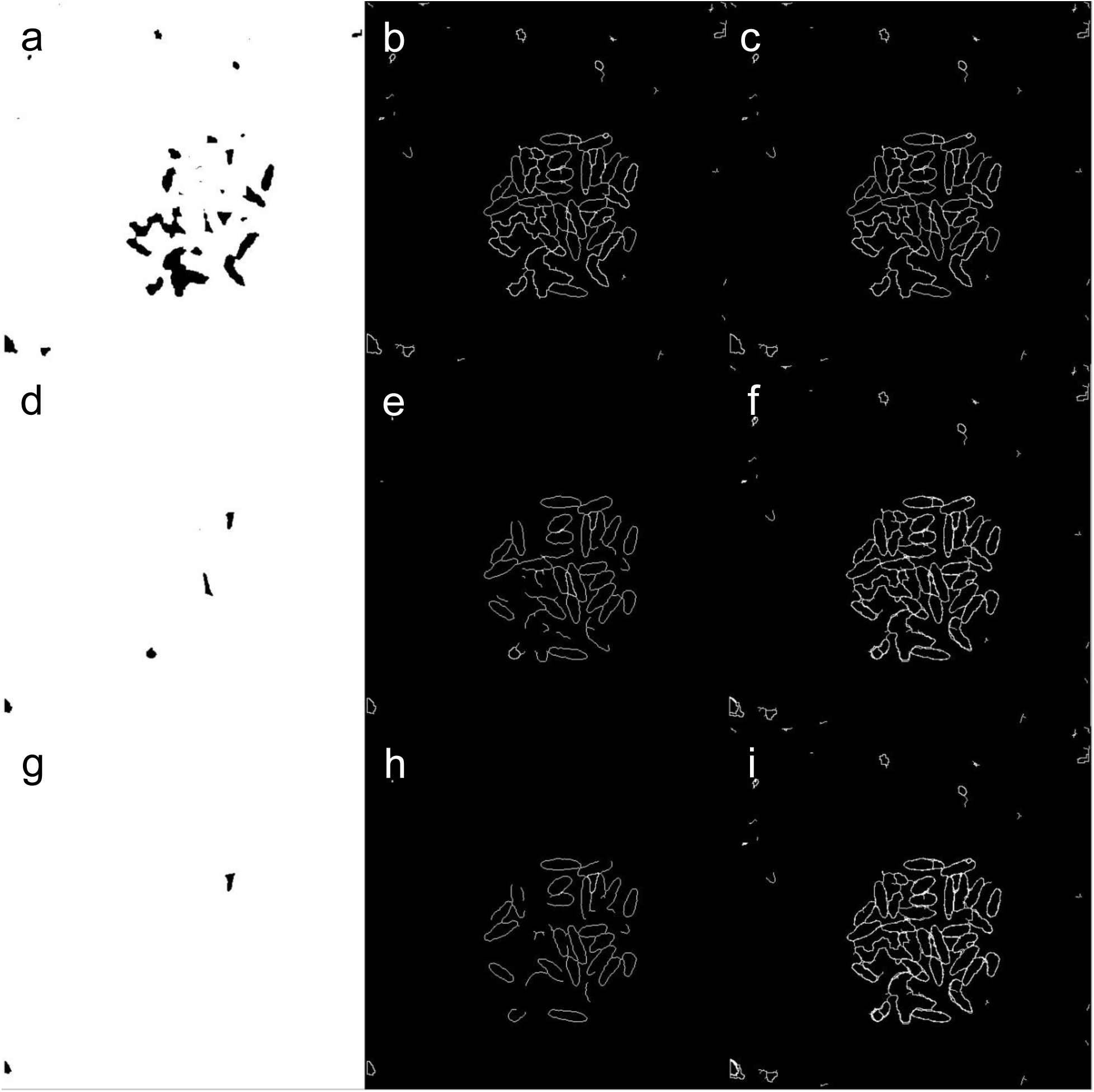
Merging thresholds 1 to 3 into the fourth target set. Some targets, seen as black blobs in panel “a,d,g”, are discovered through the negative of the clear body workflow. Panel series “a,b,c”, “d,e,f”, and “g,h,i” (continues threshold 1,2,3 respectively) show three steps: 1) dilation of the result to expand the white area and erode the black blobs, 2) subtraction of the result from the dilation which adds black blobs as outlines to our set of outlines, 3) addition of the updated result to the merge channel. As we work through the second and third series, additional data is added to the merge channel until we have a rough merge result in panel “i”.

**Figure 2C.**
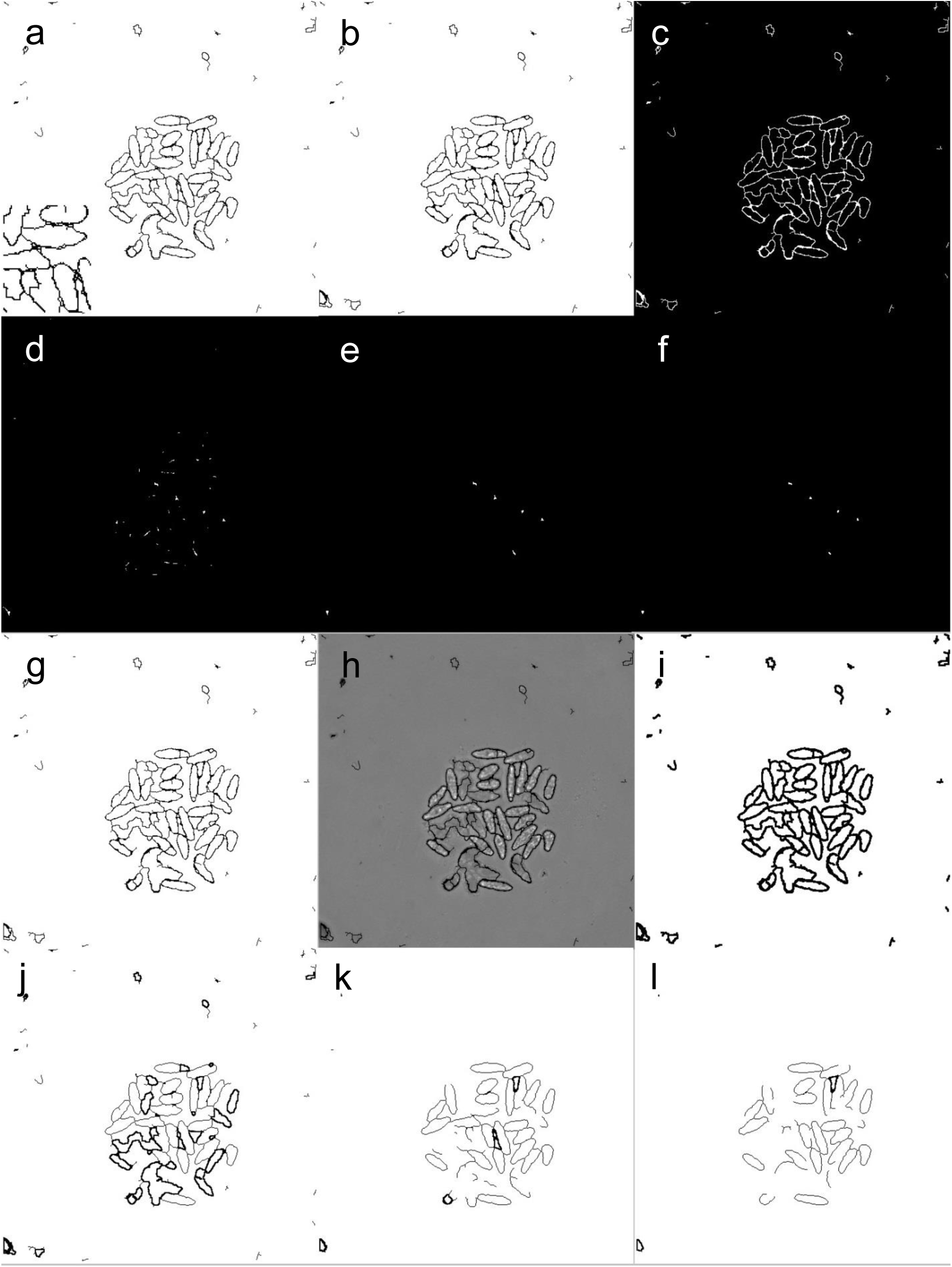
Refining merge result and preparing for target linking. The merge result has tiny little objects around the perimeter of the large objects which are produced from the intersection of three sets of outlines from thresholds 1 to 3. These small objects are filtered and added back to the image if passed (panel “a” to “g”). The finished merge result in panel “g” is assigned pixel values from the flat-field corrected image (panel “h”). The merge result is eroded to provide objects that can patch the black spaces (a requirement for target linking) in the images for the results for threshold 1 to 3. These operations are shown in panels “i” to “l”. a) inversion of the merge result with a zoomed-in region shown in the left corner to show the tiny little objects, b) sieve to remove small objects, c) inversion, d) erosion e) sieve to remove small objects, f) prune branches, g) add the result from panel “f” to the result from panel “b”, h) assign the pixel values from the flat field corrected image to any non-zero pixels in panel “g”, i) erode the result from panel “g”, j) add the result from “j” to the result from “Fig2A.g”, k) add the result from “j” to the result from “Fig2A.h”, l) add the result from “j” to the result from “Fig2A.i”. The merge result and results for threshold 1 to 3 are segmented using intensity segmentation with a minimum threshold of 1.00 and a maximum threshold of 4095.00. These settings select all non-zero pixels. The outlines of the objects produced by the preprocessing macro have pixel values of zero. The merge result segmentation has additional post processing steps which use sieve (binary) to retain objects between 1500 μm^2 and 50,000 μm^2 and border object removal to remove any objects within 5 pixels of the image border. Results from threshold 1 to 3 do not use additional post processing steps to remove objects as this could remove objects needed for target linking. Target linking uses object overlap from different images to determine which objects should be analyzed as a group. Target linking removes objects that do not overlap between all images. Target linking simplifies reporting by putting all feature data for each object found across four images on one row. To make the linking schema easier to read, the four target sets (merge, threshold1, threshold2, threshold3) are named “a”, “b”, “c”, “d”, respectively. Set “a” is linked to set “b”, “c”, and “d” using a “one to one link” to form links “ab”, “ac”, and “ad”. An object in set “a” must be within 75% of the object it is linking to otherwise the link is for those two objects are rejected. Link “ab” and “ac” are joined into link “abc” using a “composed one to one link”. Link “ab” and “ad” are joined into link “abd” using a “composed one to one link”. Link “abc” and “abd” are joined into link “abcd” (which was renamed to “e”) using a “composed one to one link”. (Figure 2D)

**Figure 2D.**
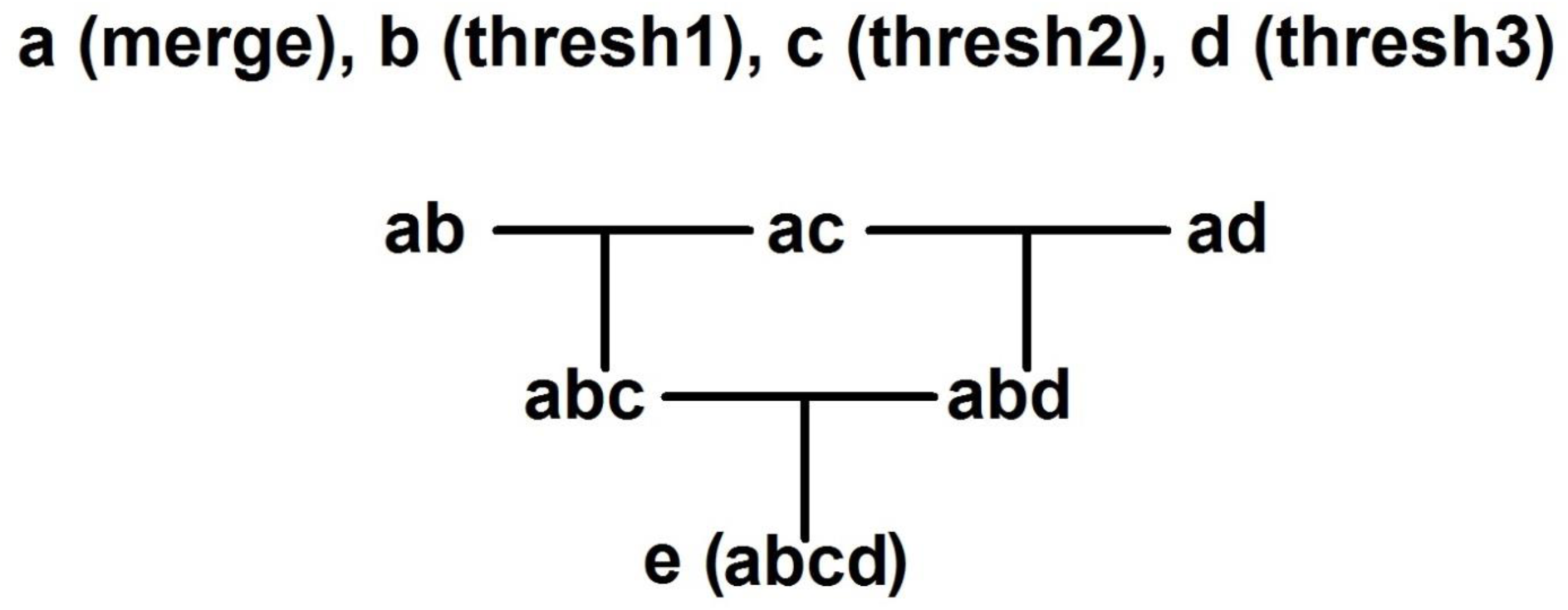
Target linking schematic. The target sets are represented by letters “a”, “b”, “c”, and “d” with “a” representing the merge data. The merge data contains all possible objects and is the root data that links to “b”, “c”, and “d” forming “ab”, “ac”, and “ad” on the first row. “Composed one to one linking” is used to link “ab” and “ac” using “matching path” data from “a” to form “abc”. “Composed one to one linking” is used to link “ab” and “ad” using “matching path” data from “a” to form “abd”. “Composed one to one linking” is used to link “abc” and “abd” using “matching path” data from “ab” to form “abcd” which is renamed to “e”. Target set “e” contains all the linking data that relates objects found at similar positions across data sets “a”, “b”, “c”, and “d”. The next macro is called “tegument outliner” because the outlines produced are based on image thresholding which target the tegument of the organism.
3. **Tegument Outliner Preprocessing** (Macro: *schisto94_g_master*) (Figure 3) (49 operations per image)

a. Transform FFC image with transform filter = median, (Macro*: schisto94_g_med1_97),* kernel = (do not apply transform)
b. Transform result from step “3.a” with local arithmetic (Macro*: schisto94_g_med1_97)*:

i. kernel = 5;
ii. sel(src<0.97*lAve,4095,0);
c. Transform result from step “3.a” with transform filter = sieve (binary), retain objects > 1500 μm^2
d. Transform result from step “3.b” with transform filter = inversion
e. Transform result from step “3.c” with transform filter = sieve (binary), retain objects > 100 μm^2
f. Transform result from step “3.b” with transform filter = inversion
g. Transform result from step “3.c” with transform filter = thinning, passes = 3
h. Transform result from step “3.d” with transform filter = pruning, passes = 3
i. Repeat steps “3.a” through “3.h” with following setting for step “3.a”, (Macro: *schisto94_g_med3_97*), kernel = 3 **AND** with the following code for step “3.b”, (Macro: *schisto94_g_med3_97*):

i. kernel = 5;
ii. sel(src<0.97*lAve,4095,0);
j. Repeat steps “3.a” through “3.h” with following setting for step “3.a”, (Macro: *schisto94_g_med3_99*), kernel = 3 **AND** with the following code for step “3.b”, (Macro: *schisto94_g_med3_99*):

i. kernel = 5;
ii. sel(src<0.99*lAve,4095,0);
k. Reset a channel with transform filter = local arithmetic

i. src = 0;
l. Apply a transform point operation = arithmetic (two src) where source1 = result from “Macro: schisto94_g_med1_97” and source2 = result from step “3.k”. The result is added to the channel used in step “3.k”.

i. src2+src1;
m. Repeat step “2.l” with source1 = result from “Macro: *schisto94_g_med3_97*”.
n. Repeat step “2.l” with source1 = result from “Macro: *schisto94_g_med3_99*”.
o. . Transform merge result (channel in step “k”) with transform filter = inversion
p. Transform result from step “3.o” with transform filter = sieve (binary), retain objects > 100 μm^2
q. Transform result from step “3.p” with transform filter = inversion
r. Transform result from step “3.q” with transform filter = erosion, kernel = 3
s. Transform result from step “3.r” with transform filter = sieve (binary), retain objects > 100 μm^2
t. Transform result from step “3.s” with transform filter = pruning, passes = 3
u. Apply a transform point operation = arithmetic (two src) where source1 = result from step “3.p” and source2 = result from step “3.t”.

i. src2+src1;
v. Apply a transform point operation = arithmetic (two src) where source1 = result from step “3.u” and source2 = result from step “1.h”.

i. sel(src1==4095,src2,0);
w. Transform result from “Macro*: schisto94_g_med1_97”* with transform filter = inversion
x. Transform result from “Macro*: schisto94_g_med3_97”* with transform filter = inversion
y. Transform result from “Macro*: schisto94_g_med3_99”* with transform filter = inversion Steps “3.a” through “3.j” generate the preliminary target data for threshold1 to threshold3. (Figure 3A) The refinement of this preliminary data and consolidation into a fourth target begins with step “3.k”. (Figure 3B) The fourth target is based on a merge of the data from the three thresholds. The resulting outline will have extra objects produced by the intersections. The extra objects are filtered to reduce complexity of the merged result and this process starts with step “3.o”. (Figure 3B) In addition, the merge result represents all possible objects for all tegument outliner target sets. Steps “3.w” through “3.y” invert the results in preparation for segmentation.

**Figure 3A.**
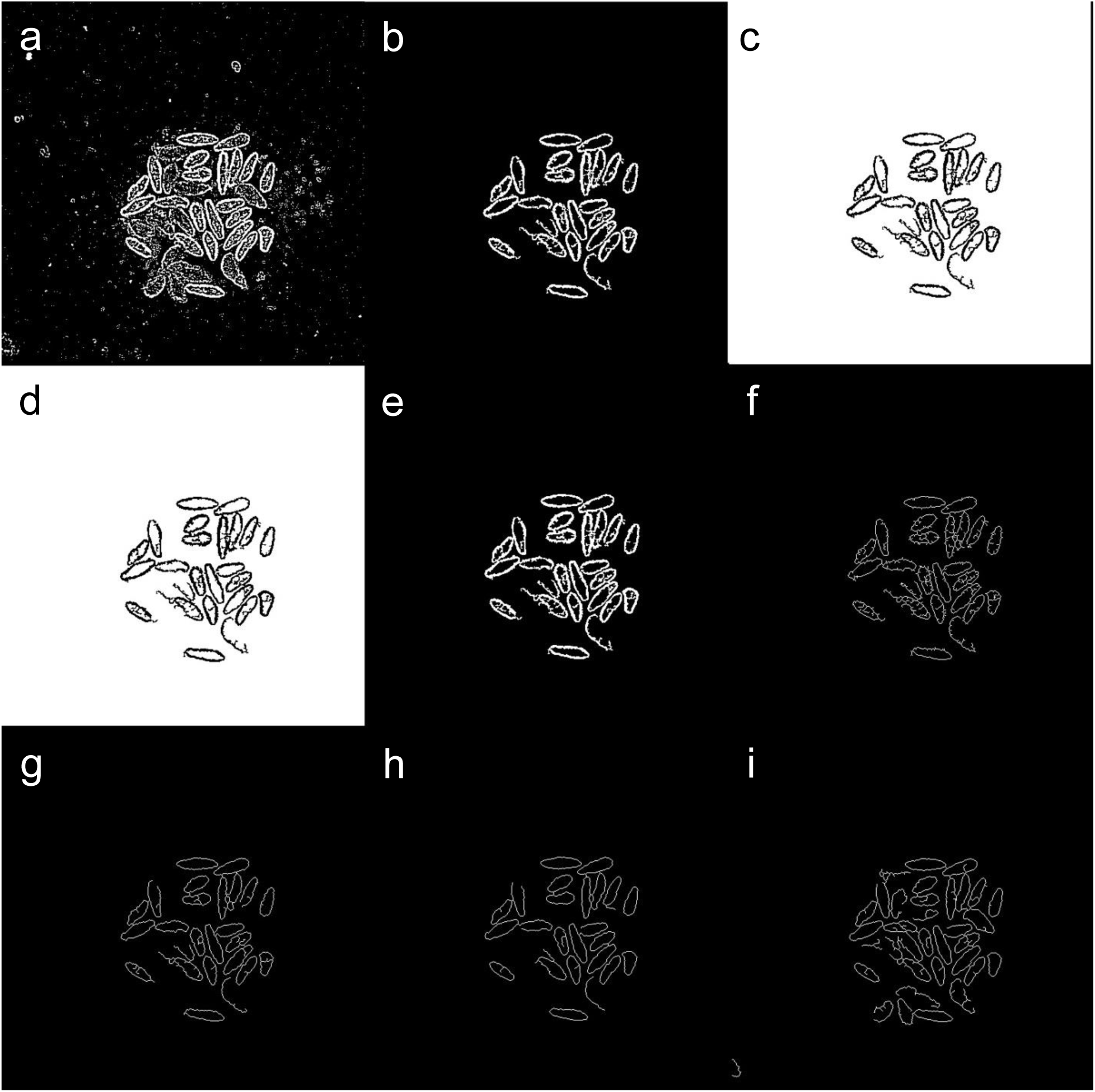
Tegument outliner workflow for thresholds 1 to 3. Panels “a” through “g” describe the tegument outliner workflow for threshold1 (“Macro*: schisto94_g_med1_97”)*. a) binarization of the flat field corrected image, b) sieve to remove small objects, c) inversion, d) sieve to remove small objects, inversion, f) thinning to reduce outline to a skeleton, g) pruning to prune back branches (this is the “Macro*: schisto94_g_med1_97”* result), h) threshold2 “Macro*: schisto94_g_med3_97*” result, i) threshold3 “Macro*: schisto94_g_med3_99*” result

**Figure 3B.**
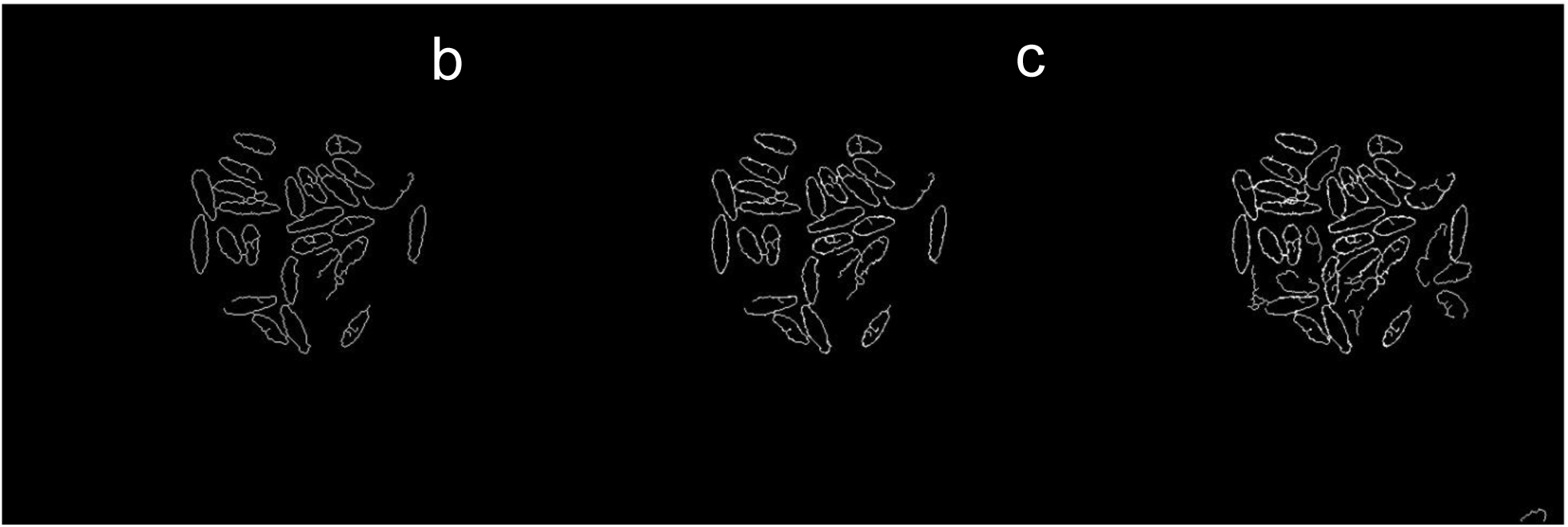
Merging thresholds 1 to 3 into the fourth target set. The results from each threshold are progressively added together into a rough merge result. a) threshold 1 result, b) threshold 1 and 2 results added together, c) threshold results 1, 2, and 3 added together. The merge result and results for threshold 1 to 3 are segmented using intensity segmentation with a minimum threshold of 1.00 and a maximum threshold of 4095.00. These settings select all non-zero pixels. The outlines of the objects produced by the preprocessing macro have pixel values of zero. The merge result segmentation has additional post processing steps which use sieve (binary) to retain objects between 1500 μm^2 and 50,000 μm^2 and border object removal to remove any objects within 5 pixels of the image border. Results from threshold 1 to 3 do not use additional post processing steps to remove objects as this could remove objects needed for target linking. Target linking uses object overlap from different images to determine which objects should be analyzed as a group. Target linking removes objects that do not overlap between all images. Target linking simplifies reporting by putting all feature data for each object found across four images on one row. To make the linking schema easier to read, the four target sets (merge, threshold1, threshold2, threshold3) are named “f”, “g”, “h”, “i”, respectively. Set “f” is linked to set “g”, “h”, and “i” using a “one to one link” to form links “fg”, “fh”, and “fi”. An object in set “f” must be within 75% of the object it is linking to otherwise the link is for those two objects are rejected. Link “fg” and “fh” are joined into link “fgh” using a “composed one to one link”. Link “fg” and “fi” are joined into link “fgi” using a “composed one to one link”. Link “fgh” and “fgi” are joined into link “fghi” (which was renamed to “j”) using a “composed one to one link”. (Figure 3D)

**Figure 3C.**
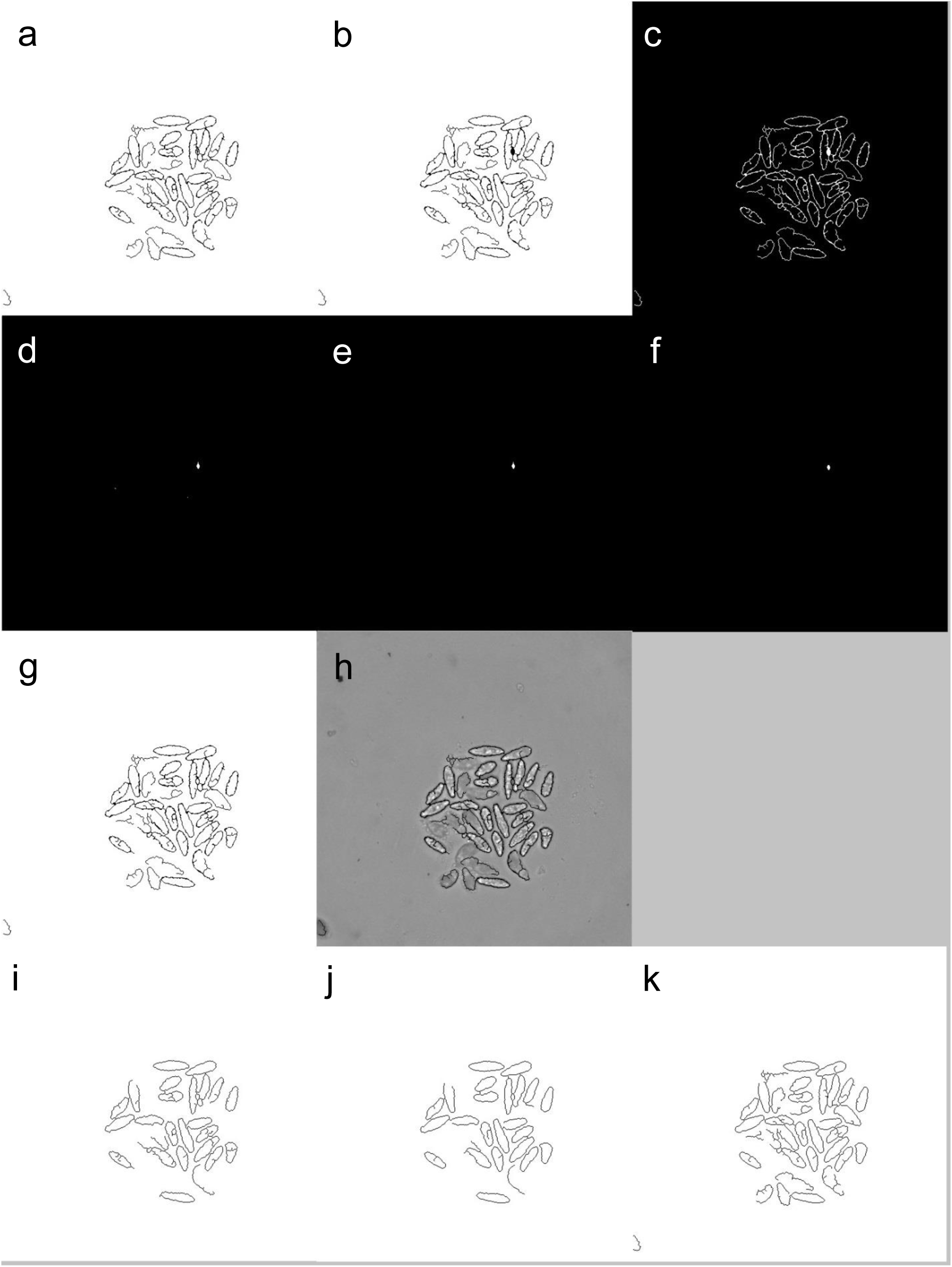
Refining merge result and preparing for target linking. The merge result has tiny little objects around the perimeter of the large objects which are produced from the intersection of three sets of outlines from thresholds 1 to 3. These small objects are filtered and added back to the image if passed (panel “a” to “g”). The finished merge result in panel “g” is assigned pixel values from the flat-field corrected image (panel “h”). The final results for the three sets of outlines from thresholds 1 to 3 are shown in panels “i” to “k” respectively. a) inversion of the merge result, b) sieve to remove small objects, c) inversion, d) erosion e) sieve to remove small objects, f) prune branches, g) add the result from panel “f” to the result from panel “b”, h) assign the pixel values from the flat field corrected image to any non-zero pixels in panel “g”, i) inversion of “Fig2A.g”, j) inversion of “Fig2A.h”, k) inversion of “Fig2A.i”

**Figure 3D.**
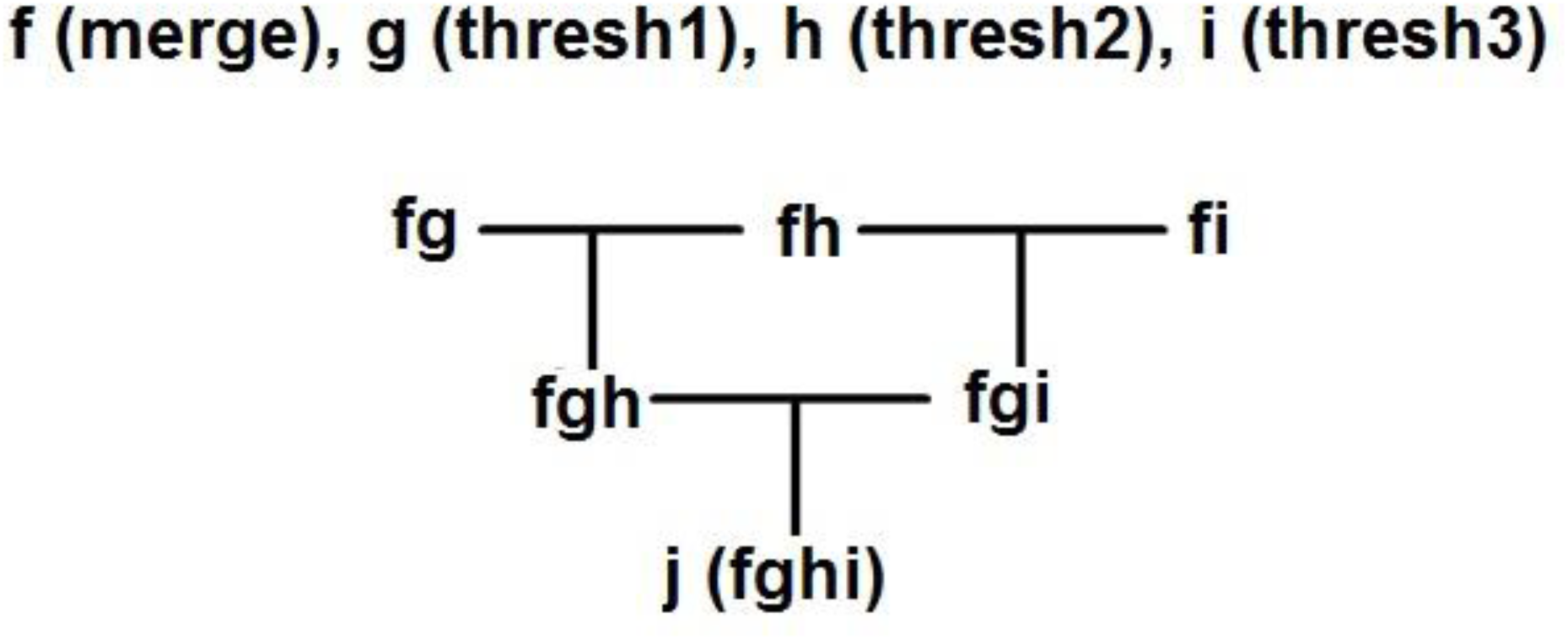
Target linking schematic. The target sets are represented by letters “f”, “g”, “h”, and “i” with “f” representing the merge data. The merge data contains all possible objects and is the root data that links to “g”, “h”, and “i” forming “fg”, “fh”, and “fi” on the first row. “Composed one to one linking” is used to link “fg” and “fh” using “matching path” data from “f” to form “fgh”. “Composed one to one linking” is used to link “fg” and “fi” using “matching path” data from “f” to form “fgi”. “Composed one to one linking” is used to link “fgh” and “fgi” using “matching path” data from “fg” to form “fghi” which is renamed to “j”. Target set “j” contains all the linking data that relates objects found at similar positions across data sets “f”, “g”, “h”, and “i”.
4. **Dark Body Outliner Preprocessing** (Macro: *schisto94_d_master*) (Figure 4)

a. Reset a channel with transform filter = local arithmetic

i. src = 0; (ch15)
b. (Macro: *schisto94_d_2050*), Transform FFC image with local arithmetic i. sel(src<2050,4095,0)
c. Transform result from step “4.b” with transform filter = sieve (binary), retain objects > 250 μm^2
d. Transform result from step “4.c” with transform filter = inversion
e. Transform result from step “4.d” with local arithmetic

i. kernel = 3
ii. sel(src>2*lAve,0,src)
f. Transform result from step “4.e” with transform filter = sieve (binary), retain objects >500 μm^2
g. Transform result from step “4.f” with transform filter = inversion (ch1)
h. Apply a transform point operation = copy result “4.g” (ch13… Macro: *schisto94_d_2050 RESULT*)
i. (Macro: *schisto94_d_fill*), Transform result from step “4.h” with local arithmetic

i. kernel =7
ii. sel(src>0,sel(src>lAve,0,4095),0)
j. Transform result from step “4.i” with transform filter = sieve (binary), retain objects > 1500 μm^2
k. Transform result from step “4.j” with transform filter = dilation, kernel = 5 (ch14… Macro: *schisto94_d_fill RESULT*)
l. Transform FFC image with local arithmetic, (Macro*: schisto94_d_1950)*

i. sel(src<1950,4095,0)
m. Transform result from step “4.l” with transform filter = sieve (binary), retain objects > 250 μm^2
n. Transform result from step “4.m” with transform filter = inversion
o. Transform result from step “4.n” with local arithmetic

i. kernel = 3
ii. sel(src>2*lAve,0,src)
p. Transform result from step “4.o” with transform filter = sieve (binary), retain objects > 500 μm^2
q. Transform result from step “4.p” with transform filter = inversion (ch1)
r. Transform result from step “4.q” with local arithmetic, (Macro*: schisto94_c17)*

i. kernel = 17
ii. sel(src>0,sel(src==lAve,4095,0),0)
s. Transform result from step “4.r” with transform filter = dilation, kernel = 3
t. Transform result from step “4.s” with transform filter = sieve (binary), retain objects < 2500 μm^2 (see step “4.r” for reason, kernel “seeds” will be small)
u. Apply a transform point operation = arithmetic (two src) where source1 = result from “Macro: *schisto94_c17*” and source2 = result from step “4.a”. The result is added to the channel used in step “4.a”. (ch15)

i. src2+src1;
v. Repeat step “4.r” through “4.v” with kernel = 15 from “Macro: *schisto94_c15*”.
w. Repeat step “4.r” through “4.v” with kernel = 13 from “Macro: *schisto94_c13*”.
x. Repeat step “4.l” through “4.x” with the following code for step “4.l” (Macro: *schisto94_d_1450*).

i. sel(src<1450,4095,0);
y. Repeat step “4.l” through “4.x” with the following code for step “4.l” (Macro: *schisto94_d_1150*). After all loops completed this is: (ch15… Merge Seed *RESULT*) i. sel(src<1150,4095,0);
z. Apply a transform point operation = arithmetic (two src) where source1 = result from step “3.u” and source2 = result from step “4.k”.

i. sel(src1<1,src1,src2); (ch16… Macro: schisto94_d_fill RESULT combined with the Tegument Outliner Merge Result)
a. Transform result from step “4.aa” with transform filter = sieve (binary), retain objects > 1500 μm^2 (ch16)
b. Apply a transform point operation = copy result “4.z” (ch1)

**Figure 4A.**
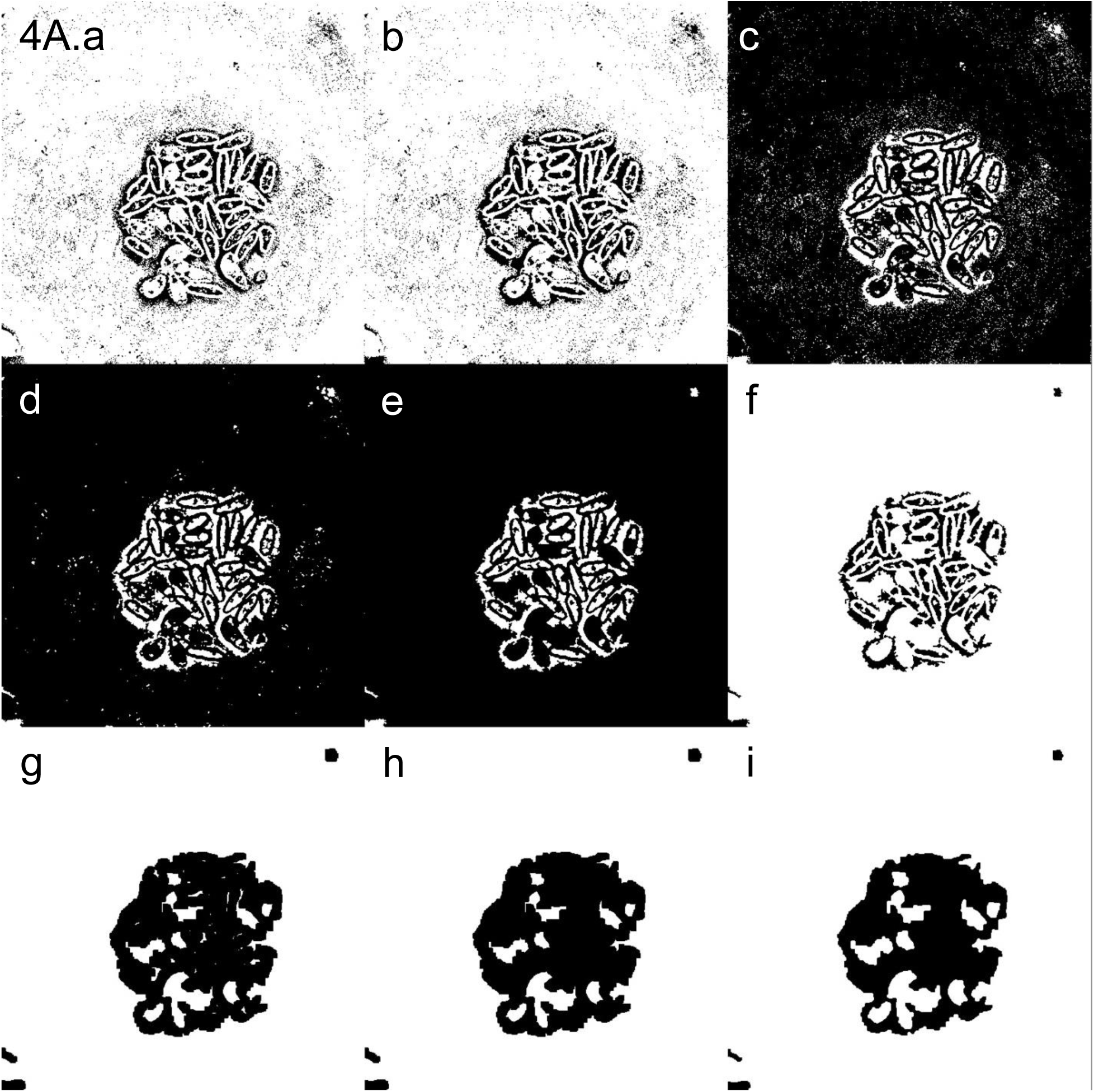
Dark body outliner part 1. Steps “4.a” through “4.k”. a) binarization, b) sieve to remove small objects, c) inversion, d) enhance dark body with local adaptive thresholding, e) sieve to remove small objects, f) inversion to get 1^st^ target image (Macro: schisto94_d_2050), g) local arithmetic to isolate dark bodies (white blobs), h) sieve to remove small objects, i) dilation to get 2^nd^ target image (Macro: schisto94_d_fill)

**Figure 4B.**
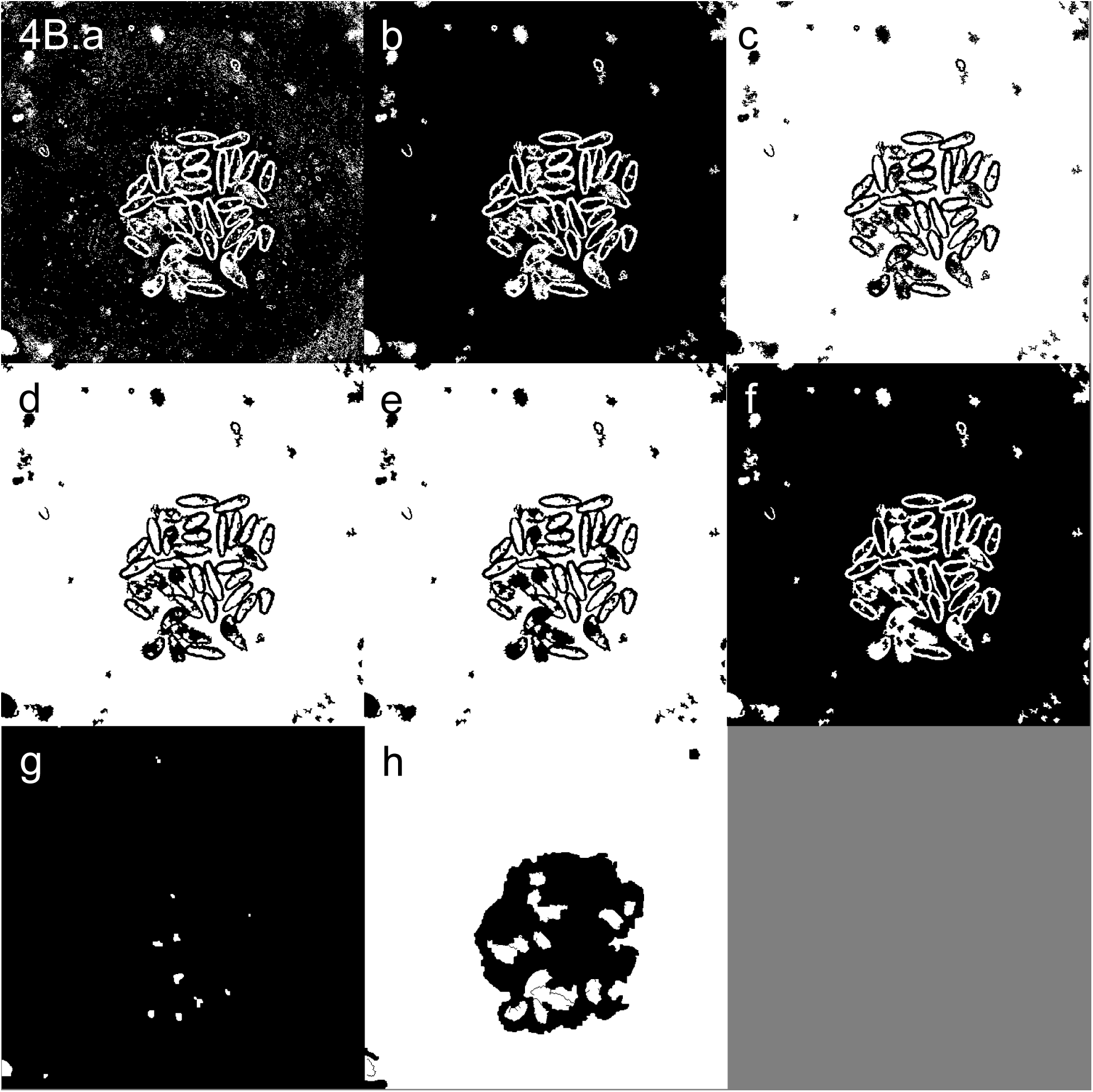
Dark body outliner part 2. Steps “4.l” through “4.v”. (Macro: *schisto94_d_1950*). a) binarization, b) sieve to remove small objects, c) inversion, d) enhance dark body with local adaptive thresholding, e) sieve to remove small objects, f) inversion, g) dark body “seeds” generated after iterative processing to get 3^rd^ target image (Macro: (see step 4.z)), h) Macro: *schisto94_d_fill* result “4.k” combined with the tegument outliner merge result “3.u” to get the 4^th^ target set (4B.h).

To detect dark bodies a variety of approaches were used. The image was scanned with varying thresholds (density = 1950, 1450, 1150) and kernels (k = 13, 15, and 17) to generate “seeds” of possible dark bodies. The “seed” target set (4B.g) is similar to previously discussed merge results for clear body and tegument outliner in that available target across multiple treatments are represented by a single object. Dark bodies are found (4A.f) by simply taking the inverse of previously described clear body outlining methods. Rather than produce an outline, as with a clear body, dark bodies produce large filled objects with the complication that these objects are sometimes surrounded by other outlines. The surrounding outlines can be diminished to leave the larger dark bodies behind, but at a cost of some distortion to the dark body shape (4A.i). Finally, the tegument outliner merge result (3C.g) is combined with the result from Macro: schisto94_d_fill (4A.h) to separate touching objects (4B.h).

The four binarized dark body target images are transformed back to original FFC image values with the following transform. Apply a transform point operation = arithmetic (two src) where source1 = binarized target image and source2 = FFC image. Code = “sel(src1==4095,src2,0);”.

*Dark body “seed” segmentation (K)*

The “seed” objects (4B.g) are segmented with intensity segmentation where objects with a pixel value > 0 are masked. The masks are post-processed with a sieve which removes objects less than 250 μm^2.

*Dark body “fill” segmentation part I (L)*

The objects from the Macro: *schisto94_d_fill* result (4.k) are segmented with intensity segmentation where objects with a pixel value > 0 are masked. Objects within 5 pixels from the border are removed using “border object removal”. The remaining masks are further post-processed with clump breaking using masks segmented in the *dark body “seed” segmentation* as seeds. The masks are post-processed with a sieve which removes objects less greater than 50,000 μm^2.

*Dark body “combo” segmentation (seed generation)*

The target image generated from Macro: *schisto94_d_fill* result “4.k” combined with the tegument outliner merge result “3.u” are segmented with intensity segmentation where objects with a pixel value > 0 are masked. The masks are post-processed with an erosion (kernel = 3) and a sieve which removes objects less greater than 500 μm^2.

*Dark body “fill” segmentation part II (N)*

The objects from the Macro: *schisto94_d_fill* result (4.k) are segmented with intensity segmentation where objects with a pixel value > 0 are masked. Objects within 5 pixels from the border are removed using “border object removal”. Clump breaking is applied to the remaining masks with masks segmented in the *dark body “combo” segmentation* as seeds. The masks are further post-processed with watershed clump breaking and a sieve which removes objects less greater than 2,000 μm^2.

*Dark body “fill” segmentation part III (O)*

The objects from the Macro: *schisto94_d_fill* result (4.k) are segmented with intensity segmentation where objects with a pixel value > 0 are masked. Objects within 5 pixels from the border are removed using “border object removal”. The masks are further post-processed with watershed clump breaking and a sieve which removes objects less greater than 2,000 μm^2.

**Figure 4D.**
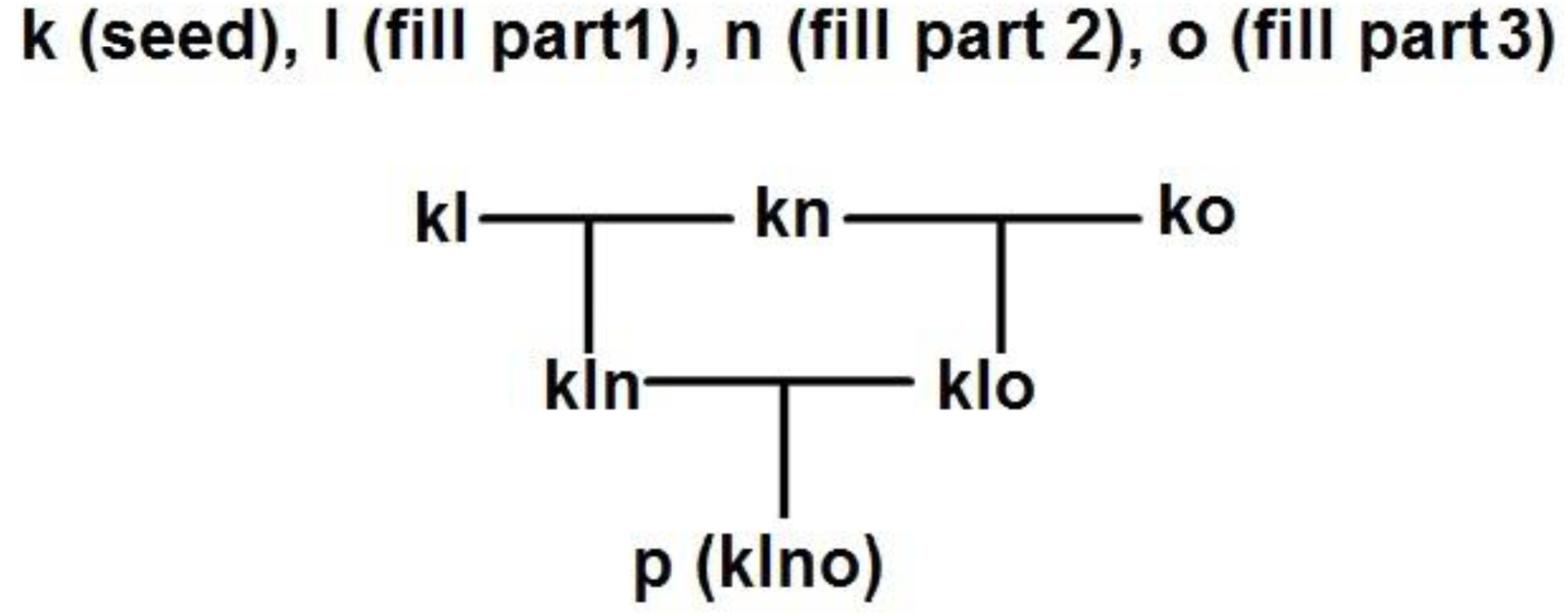
Target linking schematic. The target sets are represented by letters “k”, “l”, “n”, and “o” with “k” representing the seed data. The merge data contains all possible objects and is the root data that links to “l”, “n”, and “o” forming “kl”, “kn”, and “ko” on the first row. “Composed one to one linking” is used to link “kl” and “kn” using “matching path” data from “k” to form “kln”. “Composed one to one linking” is used to link “kl” and “ko” using “matching path” data from “k” to form “klo”. “Composed one to one linking” is used to link “kln” and “klo” using “matching path” data from “kl” to form “klno” which is renamed to “p”. Target set “p” contains all the linking data that relates objects found at similar positions across data sets “k”, “l”, “n”, and “o”.

**Figure 5A** shows various masking results among the 12 target sets generated. It appears that “**Fig5A.a**” captured most of the objects using a clear body merge set. Missing, incomplete, joined, or broken masks have more or less complete counterparts found in other panels. In total, the 12 sets of masks form a more complete set of objects that any one set can provide alone.

**Figure 5A.**
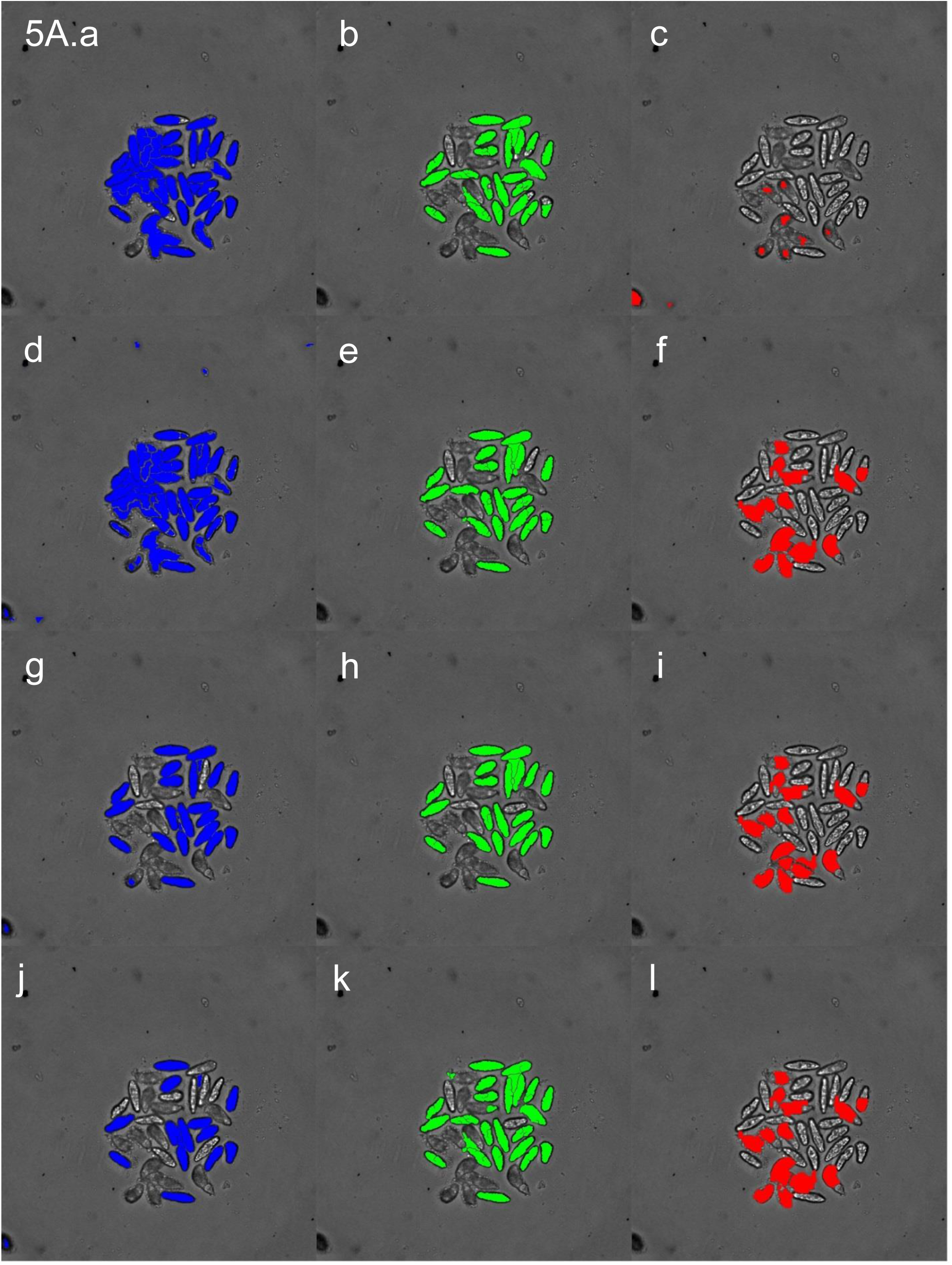
Target set masks for a mixed clear and dark body example. Clear body targets are shown in blue, tegument outliner targets are shown in green and dark body targets in red. The top row of the figure show results from the merge data sets for clear body and tegument outliner with the “seeds” from dark body shown in red. In subsequent rows, the binarization threshold becomes darker for the blue clear body target set, increase in smoothing and decrease in stringency for outline detection in green tegument outliner, and different clump breaking approaches using different seeds shown in the red dark body target set. For the clear body set: a) merge set, d) threshold 1, g) threshold 2, j) threshold 3. For the tegument outline set: b) merge set, e) threshold 1, h) threshold 2, k) threshold For the dark body target set: c) “seeds”, f) fill part 1, i) fill part 2, l) fill part 3.

**Figure 5B.**
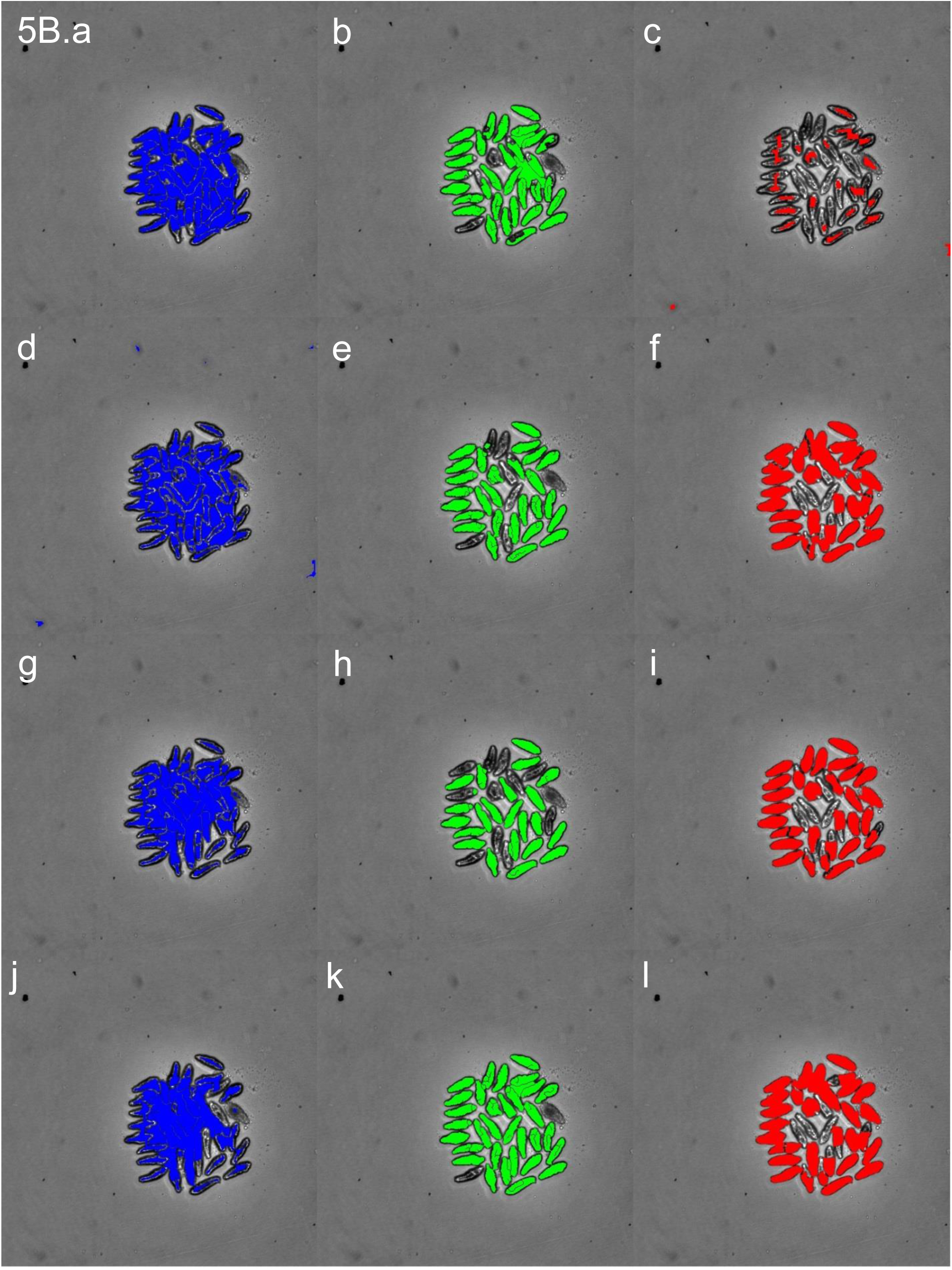
shows various masking results among the 12 target sets generated. It appears that the clear body workflow captured most of the objects using a clear body merge set but most of the masks are joined to other masks leading to a poor segmentation result. The dark body workflow performs better but still suffers from objects that are masked together. The tegument outliner performed the best in terms of finding individual objects with accurate masks. Missing, incomplete, joined, or broken masks have more or less complete counterparts found in other panels. In total, the 12 sets of masks form a more complete set of objects that any one set can provide alone.

**Figure 5C** shows various masking results among the 12 target sets generated. It appears that the clear body workflow captured most of the objects using a clear body merge set but most of the masks are joined to other masks leading to a poor segmentation result. The tegument outliner workflow performs better but still suffers from objects that are masked together or missing altogether. The dark body workflow performed the best in terms of finding individual objects with accurate masks. Missing, incomplete, joined, or broken masks have more or less complete counterparts found in other panels. In total, the 12 sets of masks form a more complete set of objects that any one set can provide alone.

**Figure 5C.**
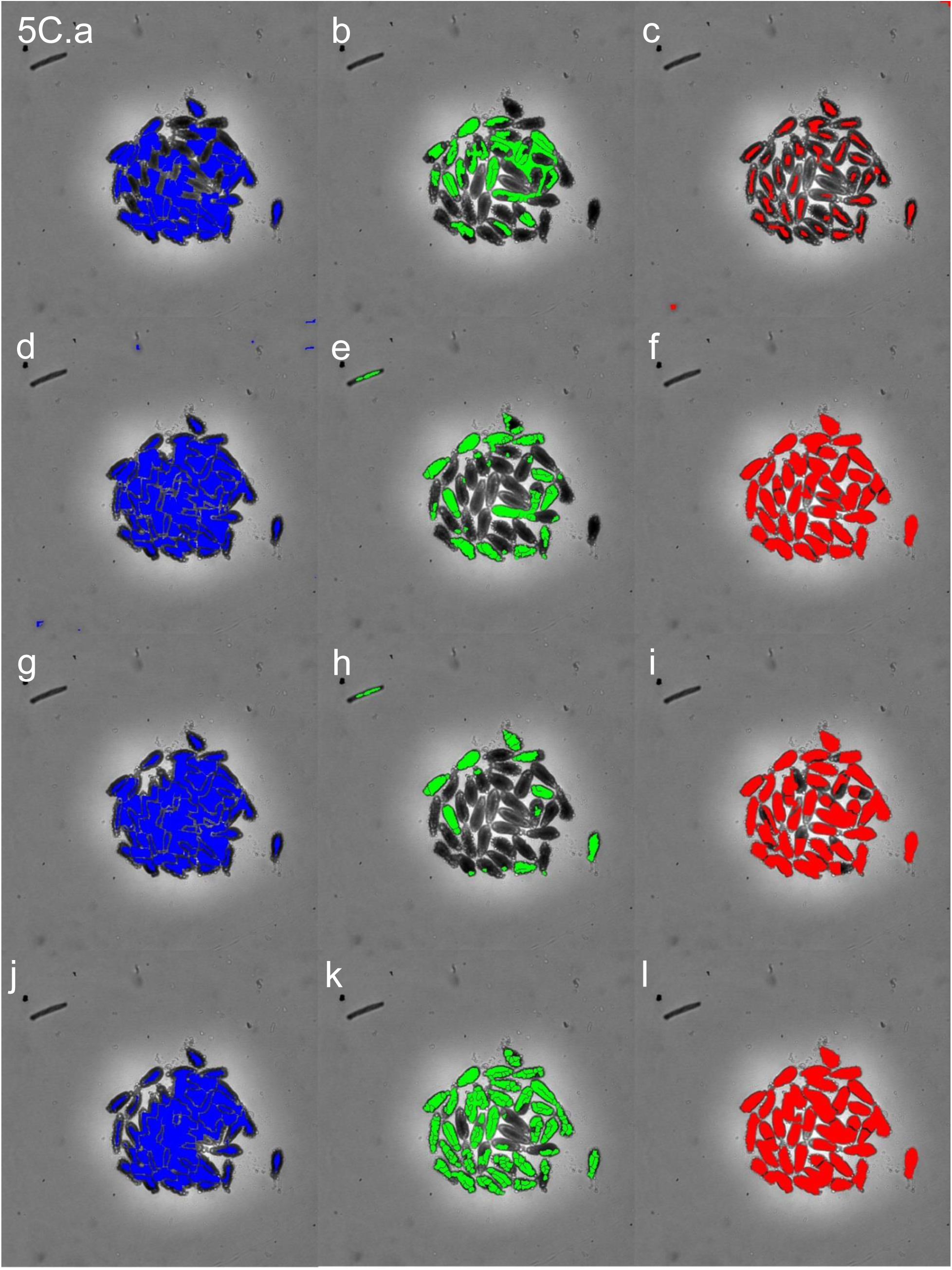
Target set masks for dark body. Clear body targets are shown in blue, tegument outliner targets are shown in green and dark body targets in red. The top row of the figure show results from the merge data sets for clear body and tegument outliner with the “seeds” from dark body shown in red. In subsequent rows, the binarization threshold becomes darker for the blue clear body target set, increase in smoothing and decrease in stringency for outline detection in green tegument outliner, and different clump breaking approaches using different seeds shown in the red dark body target set. For the clear body set: a) merge set, d) threshold 1, g) threshold 2, j) threshold 3. For the tegument outline set: b) merge set, e) threshold 1, h) threshold 2, k) threshold 3. For the dark body target set: c) “seeds”, f) fill part 1, i) fill part 2, l) fill part 3.

Features (area, length, color, etc) are recorded for every object in all 12 target sets with an IN Cell Developer analysis time of about 5 hours per plate. The data is exported to a comma separated file (CSV) typically weighing in at approximately100 megabytes and containing more than 40 million data points to describe 200,000 objects. Macro parameters can be easily modified to extend the range or resolution of this approach, or even to adapt segmentation to different organisms.

### Data Pre-Processing

Before the data from IN Cell Developer can be analyzed and turned into descriptors for database entry, the large data file requires treatment for the following:

1. Correct for segmentation feature offset (bias introduced from differences in segmentation)
2. Choose the best mask out of the twelve possible masks to represent the object. There may be no best object if all fail to be within certain size limits.
3. Use organism-level and well-level data in both appearance and motion-based descriptors to classify objects as either “clear body”, “dark body”, or non-organism objects. Non-organism objects are removed from further analysis.

**Figure 6.**
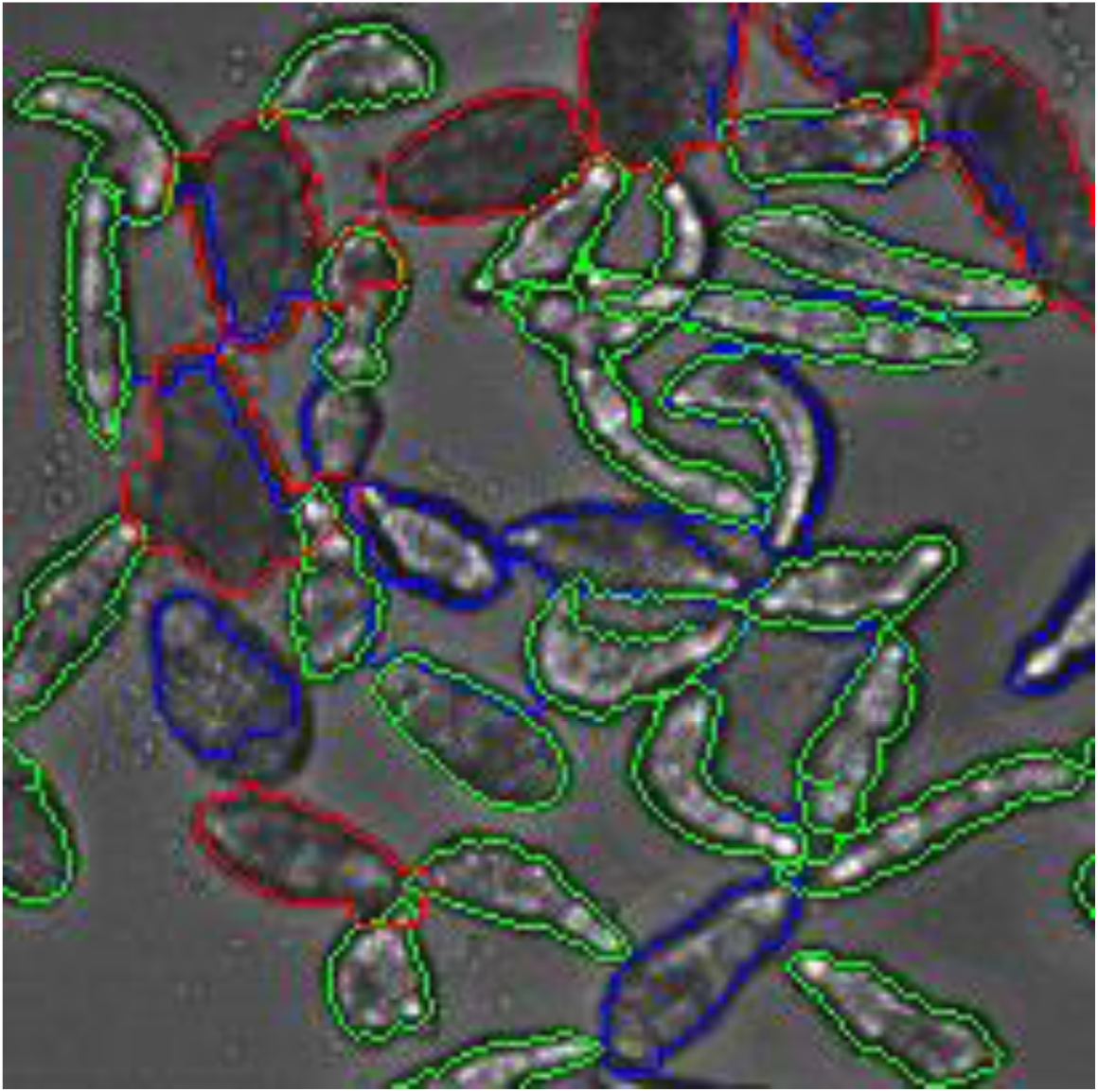
MySQL database. A model of the database is shown with tables represented by blue boxes. Lines connecting boxes describe the relationship of the tables to each other.

### Choosing the Best Mask

In the process of choosing the best mask, the objects are first filtered and reorganized. Objects that are too small or too large are removed from analysis. The next step is to link objects within each workflow by x and y position over time. Objects are linked to the coordinates of the last time-point or to an average of all the previous time-points when there are gaps. The workflow that generated masks with the most persistence over time was selected as the best series of masks to move forward with analysis. In the case where workflows generated series with the same amount of persistence, the series with the lower x,y variability over time was selected.

### Object Classification

The object classifier uses organism-level data and well-level data to classify objects as clear body, dark body, intra-body artifact, or inter-body artifact. First, the objects were linked through the time-lapse images by x,y position. The time-linked objects were then used to generate feature descriptors (such as mean, standard deviation) at both levels. Descriptors were then entered into Model_1 to classify an object as a clear object or a dark object. Each additional level of classification refines a part of the previous result. Model_2_Clear then classifies a clear object as a clear organism or a clear non-organism (e.g. an object formed between organisms). Model_3_Clear then classifies a clear organism as a complete clear organism or a partial clear organism (e.g. a partial masking of the organism due to an internal boundary). Model_2_Dark and Model_3_Dark perform the same operations but for the dark objects from Model_1.

1. Set objects in the first available time point as reference positions.
2. Set objects in the next available time points at test positions.
3. If a test position is less than 35 μm^2 from a reference position than record a time linkage.
4. Set test positions as reference positions.
5. Repeat steps 2 through 4 until all time points have been processed.
6. Calculate mean and standard deviation at the organism level and at the well level.
7. Calculate the persistence at the organism level and at the well level. Persistence is the count of time-linkages divided by the total number of time-lapse points for a given object. Objects with < 5% persistence are removed from analysis.
8. Classify an object as a clear object or a dark object (model_1: clear < 0.62)

a. Classify a clear object as a clear organism or a clear non-organism object (model_2_clear: 0.75 < non-organism < 2.5)

i. Classify a clear organism as a complete clear organism or a partial clear organism (model_3_clear: 0.48 < partial < 2.5)
b. Classify a dark object as a dark organism or a dark non-organism object (model_2_dark:0.8 < non-organism < 2.5)

i. Classify a dark organism as a complete dark organism or a partial dark organism (model_3_dark: 0.54 < partial < 2.5)
9. Write data for clear and dark complete organisms to a new CSV file.

### Model_1

n(0) = percent persistence (organism level); n(1) = density – levels mean (organism level); n(2) = SD – levels mean (organism level); n(3) = form factor mean (well level); n(4) = density – levels mean (well level)

R1 = 18.25203487215 + 1.161068183069 * n(0) + 0.75542095675251 * n(1) + -0.68305180301445 * n(2) + -1781.6761055724 * n(3) + -0.045006141550507 * n(4) + 3.2365931882399E-04 * n(0) * n(1) + - 4.1740004117079E-04 * n(0) * n(2) + -4.6310458346469 * n(0) * n(3) + -5.7890475949427E-04 * n(0) * n(4) + -9.8420513221299E-05 * n(1) * n(2) + -2.1884015714641 * n(1) * n(3) + -2.7848992087403E-04 * n(1) * n(4)

R2 = 1.3369398769791 * n(2) * n(3) + 8.9582262126079E-04 * n(2) * n(4) + 3.3329037457293 * n(3) * n(4) + -1.8232199977951E-03 * n(0) ^ 2 + -9.4790505473488E-05 * n(1) ^ 2 + -2.2142492356085E-04 * n(2) ^ 2 + 2134.9935522591 * n(3) ^ 2 + -4.7102296313054E-04 * n(4) ^ 2 + 1.2266056885224E-06 * n(0) * n(1) * n(2) + -9.2354913779362E-04 * n(0) * n(1) * n(3) + -1.6491057509021E-07 * n(0) * n(1) * n(4) + 1.5522216710331E-04 * n(0) * n(2) * n(3) + -8.2092941369132E-07 * n(0) * n(2) * n(4)

R3 = 2.9992856931552E-03 * n(0) * n(3) * n(4) + -3.5217569712729E-04 * n(1) * n(2) * n(3) + 6.5107500960996E-08 * n(1) * n(2) * n(4) + 2.6791800633611E-04 * n(1) * n(3) * n(4) + - 9.0380771936169E-04 * n(2) * n(3) * n(4) + 3.1247731735274E-06 * n(0) ^ 2 * n(2) + 5.6240376267275E-03 * n(0) ^ 2 * n(3) + -1.4549471230916E-06 * n(0) ^ 2 * n(4) + -7.2417177105256E-08 * n(0) * n(1) ^ 2 + -6.1772369143247E-07 * n(0) * n(2) ^ 2 + 3.781429940739 * n(0) * n(3) ^ 2 + 1.0418245071802E-07 * n(0) * n(4) ^ 2 + 1.0219248176663E-07 * n(1) ^ 2 * n(2)

R4 = 4.5564375868527E-04 * n(1) ^ 2 * n(3) + -3.2106682445134E-08 * n(1) ^ 2 * n(4) + - 1.8650850919366E-07 * n(1) * n(2) ^ 2 + 1.9206238869076 * n(1) * n(3) ^ 2 + 1.1738445506832E-07 * n(1) * n(4) ^ 2 + 3.7442327867001E-05 * n(2) ^ 2 * n(3) + 3.1503030609951E-07 * n(2) ^ 2 * n(4) + - 0.24033752947827 * n(2) * n(3) ^ 2 + -4.3786761514222E-07 * n(2) * n(4) ^ 2 + -3.4671132225482 * n(3) ^ 2 * n(4) + -6.8418530427196E-04 * n(3) * n(4) ^ 2 + 7.6446830499591E-06 * n(0) ^ 3 + - 1.0108280682996E-08 * n(1) ^ 3

R5 = 1.2530435647321E-07 * n(2) ^ 3 + -682.50789300831 * n(3) ^ 3 + 2.3776592427786E-07 * n(4) ^ 3 + -8.3535696931061E-07 * n(0) * n(1) * n(2) * n(3) + -1.5257479853204E-10 * n(0) * n(1) * n(2) * n(4) + 4.0724584996769E-07 * n(0) * n(1) * n(3) * n(4) + 4.604799340958E-07 * n(0) * n(2) * n(3) * n(4) + - 8.7746281304306E-10 * n(0) ^ 2 * n(2) ^ 2 + -3.3302009301847E-06 * n(0) ^ 2 * n(2) * n(3) + - 4.756929644064E-10 * n(0) ^ 2 * n(2) * n(4) + -3.4228762882309E-03 * n(0) ^ 2 * n(3) ^ 2 + 6.5899847343375E-10 * n(0) ^ 2 * n(4) ^ 2 + -1.1829905701319E-10 * n(0) * n(1) ^ 2 * n(2)

R6 = -6.5159315821518E-08 * n(0) * n(1) ^ 2 * n(3) + 5.2720741951882E-04 * n(0) * n(1) * n(3) ^ 2 + 4.9534714604476E-07 * n(0) * n(2) ^ 2 * n(3) + 4.1269226932606E-10 * n(0) * n(2) ^ 2 * n(4) + 6.3407631003489E-04 * n(0) * n(2) * n(3) ^ 2 + 1.494970625121E-10 * n(0) * n(2) * n(4) ^ 2 + - 1.6907505821672E-03 * n(0) * n(3) ^ 2 * n(4) + -5.1365605396076E-07 * n(0) * n(3) * n(4) ^ 2 + - 3.6155421452363E-11 * n(1) ^ 2 * n(2) ^ 2 + -4.8126433937627E-08 * n(1) ^ 2 * n(2) * n(3) + - 1.7625953726279E-11 * n(1) ^ 2 * n(2) * n(4) + -3.2351132584444E-04 * n(1) ^ 2 * n(3) ^ 2 + - 7.9530874313422E-08 * n(1) ^ 2 * n(3) * n(4)

R7 = -2.669952144001E-11 * n(1) ^ 2 * n(4) ^ 2 + 1.4503329430718E-07 * n(1) * n(2) ^ 2 * n(3) + 1.4365500750249E-10 * n(1) * n(2) ^ 2 * n(4) + 4.2874540494959E-04 * n(1) * n(2) * n(3) ^ 2 + - 3.6294914163709E-11 * n(1) * n(2) * n(4) ^ 2 + -3.3805927443037E-04 * n(2) ^ 2 * n(3) ^ 2 + - 1.4210910495358E-10 * n(2) ^ 2 * n(4) ^ 2 + 2.3003285709267E-07 * n(2) * n(3) * n(4) ^ 2 + 4.9777029628825E-04 * n(3) ^ 2 * n(4) ^ 2 + 2.97760253571E-09 * n(0) ^ 3 * n(2) + - 4.3123259790407E-09 * n(0) ^ 3 * n(4) + 2.8293207850737E-11 * n(0) * n(1) ^ 3 + - 2.6346389605596E-10 * n(0) * n(2) ^ 3

R8 = -0.67260141510363 * n(0) * n(3) ^ 3 + 1.8752589917427E-08 * n(1) ^ 3 * n(3) + 3.5136792540332E-11 * n(1) ^ 3 * n(4) + -1.7714456049114E-11 * n(1) * n(2) ^ 3 + - 0.46479901154684 * n(1) * n(3) ^ 3 + -3.4875673740552E-08 * n(2) ^ 3 * n(3) + -2.5841449832186E- 11 * n(2) ^ 3 * n(4) + -0.22134015982785 * n(2) * n(3) ^ 3 + 8.7853677440858E-11 * n(2) * n(4) ^ 3 + 0.86618600900044 * n(3) ^ 3 * n(4) + -9.5374757135764E-12 * n(1) ^ 4 + -3.6583886794751E-11 * n(4) ^ 4 + 0

Model_1 = R1 + R2 + R3 + R4 + R5 + R6 + R7 + R8

### Model_2_Clear

n(0) = percent persistence (organism level); n(1) = density – levels mean (organism level); n(2) = SD – levels mean (well level); n(3) = density – levels mean (well level) ; n(4) = pinch mean (organism level) ; n(5) = model_1 (organism level) ; n(6) = area mean (organism level)

R1 = 3050.53971 + -15.56494 * n(0) + -0.83465 * n(1) + -4.70861 * n(2) + -1.83172 * n(3) + - 1087.22384 * n(4) + -1798.96904 * n(5) + 0.00388693 * n(6) + 0.00701913 * n(0) * n(1) + 0.00621377 * n(0) * n(2) + 0.012155 * n(0) * n(3) + 1.18447 * n(0) * n(4) + 3.5422 * n(0) * n(5) + 0.000836635 * n(0) * n(6)

R2 = 0.00167653 * n(1) * n(2) + 0.000382736 * n(1) * n(3) + 0.82585 * n(1) * n(4) + 0.48752 * n(1) * n(5) + -0.0000204636 * n(1) * n(6) + 0.00409758 * n(2) * n(3) + 0.22639 * n(2) * n(4) + 0.50064 * n(2) * n(5) + 0.00000930964 * n(2) * n(6) + -0.2123 * n(3) * n(4) + 1.34517 * n(3) * n(5) + -0.000049354 * n(3) * n(6) + 467.08521 * n(4) * n(5) + 0.065277 * n(4) * n(6)

R3 = 0.022834 * n(5) * n(6) + -0.032424 * n(0) ^ 2 + -0.00016082 * n(1) ^ 2 + 0.000283988 * n(2) ^ 2 + 0.0000706305 * n(3) ^ 2 + 665.10179 * n(4) ^ 2 + 297.72877 * n(5) ^ 2 + -0.00000288682 * n(6) ^ 2 + 0.000000576592 * n(0) * n(1) * n(2) + -0.00000550441 * n(0) * n(1) * n(3) + -0.00344384 * n(0) * n(1) * n(4) + -0.000418887 * n(0) * n(1) * n(5) + -0.00000000266297 * n(0) * n(1) * n(6) + -0.00000537496 * n(0) * n(2) * n(3)

R4 = 0.00123966 * n(0) * n(2) * n(4) + -0.000539946 * n(0) * n(2) * n(5) + -0.000000269514 * n(0) * n(2) * n(6) + 0.000685879 * n(0) * n(3) * n(4) + -0.00295768 * n(0) * n(3) * n(5) + -0.000000600777 * n(0) * n(3) * n(6) + 0.45772 * n(0) * n(4) * n(5) + -0.0000773276 * n(0) * n(4) * n(6) + -0.0000149499 * n(0) * n(5) * n(6) + -0.00000121635 * n(1) * n(2) * n(3) + 0.0000187735 * n(1) * n(2) * n(4) + -0.000393188 * n(1) * n(2) * n(5) + -0.00000000836334 * n(1) * n(2) * n(6) + 0.000237552 * n(1) * n(3) * n(4)

R5 = -0.000170248 * n(1) * n(3) * n(5) + 0.0000000424461 * n(1) * n(3) * n(6) + -0.13473 * n(1) * n(4) * n(5) + -0.0000398496 * n(1) * n(4) * n(6) + -0.00000609553 * n(1) * n(5) * n(6) + -0.000155192 * n(2) * n(3) * n(4) + -0.000135105 * n(2) * n(3) * n(5) + -0.00000000770149 * n(2) * n(3) * n(6) + 0.12141 * n(2) * n(4) * n(5) + 0.00000677018 * n(2) * n(4) * n(6) + -0.0000221139 * n(2) * n(5) * n(6) + -0.17437 * n(3) * n(4) * n(5) + -0.0000342233 * n(3) * n(4) * n(6) + 0.00000269916 * n(3) * n(5) * n(6)

R6 = -0.014 * n(4) * n(5) * n(6) + -0.00000661836 * n(0) ^ 2 * n(1) + -0.00000267154 * n(0) ^ 2 * n(2) + 0.0000291939 * n(0) ^ 2 * n(3) + 0.00558697 * n(0) ^ 2 * n(4) + 0.00128265 * n(0) ^ 2 * n(5) + 0.00000138321 * n(0) ^ 2 * n(6) + 0.00000042153 * n(0) * n(1) ^ 2 + -0.000001289 * n(0) * n(2) ^ 2 + - 0.00000234632 * n(0) * n(3) ^ 2 + 1.26238 * n(0) * n(4) ^ 2 + -0.17282 * n(0) * n(5) ^ 2 + - 0.0000000408076 * n(0) * n(6) ^ 2 + -0.000000130828 * n(1) ^ 2 * n(2)

R7 = 0.000000125061 * n(1) ^ 2 * n(3) + -0.000289823 * n(1) ^ 2 * n(4) + -0.000023126 * n(1) ^ 2 * n(5) + -0.0000001232 * n(1) * n(2) ^ 2 + -0.000000121779 * n(1) * n(3) ^ 2 + -0.23081 * n(1) * n(4) ^ 2 + - 0.16611 * n(1) * n(5) ^ 2 + 0.000000158732 * n(2) ^ 2 * n(3) + -0.0000495972 * n(2) ^ 2 * n(4) + 0.000141303 * n(2) ^ 2 * n(5) + 0.0000000097208 * n(2) ^ 2 * n(6) + -0.00000110647 * n(2) * n(3) ^ 2 + -0.10298 * n(2) * n(4) ^ 2 + 0.026119 * n(2) * n(5) ^ 2

R8 = 0.00000000227912 * n(2) * n(6) ^ 2 + 0.000183837 * n(3) ^ 2 * n(4) + -0.000396325 * n(3) ^ 2 * n(5) + 0.0000000245561 * n(3) ^ 2 * n(6) + -0.31387 * n(3) * n(4) ^ 2 + -0.033724 * n(3) * n(5) ^ 2 + 0.000000000433223 * n(3) * n(6) ^ 2 + -126.01624 * n(4) ^ 2 * n(5) + -0.00723282 * n(4) ^ 2 * n(6) + - 88.82786 * n(4) * n(5) ^ 2 + 0.00000545089 * n(4) * n(6) ^ 2 + -0.012727 * n(5) ^ 2 * n(6) + - 0.000000923552 * n(5) * n(6) ^ 2 + 0.000027338 * n(0) ^ 3

R9 = 0.0000000615183 * n(1) ^ 3 + -0.00000021971 * n(2) ^ 3 + 0.000000176908 * n(3) ^ 3 + - 60.56525 * n(4) ^ 3 + -13.42882 * n(5) ^ 3 + 0.000000000125105 * n(6) ^ 3 + 8.83587E-11 * n(0) * n(1) * n(2) * n(6) + 0.00000163142 * n(0) * n(1) * n(3) * n(4) + 0.000000287771 * n(0) * n(1) * n(3) * n(5) + - 0.000188377 * n(0) * n(1) * n(4) * n(5) + -0.000000867302 * n(0) * n(2) * n(3) * n(4) + 0.000000000132956 * n(0) * n(2) * n(3) * n(6) + 0.000514007 * n(0) * n(2) * n(4) * n(5) + - 0.0000000938365 * n(0) * n(2) * n(4) * n(6)

R10 = -0.000256585 * n(0) * n(3) * n(4) * n(5) + 0.000000144641 * n(0) * n(3) * n(4) * n(6) + 0.00000017782 * n(1) * n(2) * n(3) * n(5) + -0.000033291 * n(1) * n(2) * n(4) * n(5) + 0.00000000921168 * n(1) * n(2) * n(5) * n(6) + -0.0000684562 * n(1) * n(3) * n(4) * n(5) + 0.0000000189874 * n(1) * n(3) * n(4) * n(6) + -0.00013601 * n(2) * n(3) * n(4) * n(5) + -0.00000000626452 * n(2) * n(3) * n(5) * n(6) + 0.0000154375 * n(2) * n(4) * n(5) * n(6) + 0.0000000022209 * n(0) ^ 2 * n(1) ^ 2 + -0.00000000030416 * n(0) ^ 2 * n(1) * n(6) + -0.00000000197509 * n(0) ^ 2 * n(2) ^ 2 + 0.00000301416 * n(0) ^ 2 * n(2) * n(4)

R11 = -0.00000000426287 * n(0) ^ 2 * n(3) ^ 2 + -0.00000500198 * n(0) ^ 2 * n(3) * n(4) + - 0.00000142532 * n(0) ^ 2 * n(3) * n(5) + -0.000000000528475 * n(0) ^ 2 * n(3) * n(6) + 0.00075716 * n(0) ^ 2 * n(4) ^ 2 + 0.00120644 * n(0) ^ 2 * n(4) * n(5) + 0.000000207233 * n(0) ^ 2 * n(4) * n(6) + - 0.000000000324208 * n(0) * n(1) ^ 2 * n(2) + -0.000000000244258 * n(0) * n(1) ^ 2 * n(3) + 0.000000000461462 * n(0) * n(1) * n(2) ^ 2 + 0.00000000112886 * n(0) * n(1) * n(3) ^ 2 + 0.000000635517 * n(0) * n(2) ^ 2 * n(4) + -0.000000000036482 * n(0) * n(2) ^ 2 * n(6) + 0.000000001308 * n(0) * n(2) * n(3) ^ 2

R12 = -0.000206579 * n(0) * n(2) * n(4) ^ 2 + -1.17651E-11 * n(0) * n(2) * n(6) ^ 2 + -0.000000833799 * n(0) * n(3) ^ 2 * n(4) + 0.000000636733 * n(0) * n(3) ^ 2 * n(5) + 0.000000000061738 * n(0) * n(3) ^ 2 * n(6) + 0.0000662669 * n(0) * n(3) * n(5) ^ 2 + 2.40122E-11 * n(0) * n(3) * n(6) ^ 2 + -0.0000818538 * n(0) * n(4) ^ 2 * n(6) + 0.043108 * n(0) * n(4) * n(5) ^ 2 + -0.00000000647759 * n(0) * n(4) * n(6) ^ 2 + 0.00000000165957 * n(0) * n(5) * n(6) ^ 2 + -0.00000000014381 * n(1) ^ 2 * n(2) ^ 2 + - 0.000000000318973 * n(1) ^ 2 * n(2) * n(3) + 0.0000000115171 * n(1) ^ 2 * n(2) * n(4)

R13 = 0.000000119935 * n(1) ^ 2 * n(3) * n(4) + -0.000000100809 * n(1) ^ 2 * n(3) * n(5) + 0.0000705524 * n(1) ^ 2 * n(4) * n(5) + 0.00000619885 * n(1) ^ 2 * n(5) ^ 2 + 0.000000000267632 * n(1) * n(2) ^ 2 * n(3) + -0.000000051141 * n(1) * n(2) ^ 2 * n(4) + 0.000000000524427 * n(1) * n(2) * n(3) ^ 2 + -0.000000268552 * n(1) * n(3) ^ 2 * n(4) + 0.000000121555 * n(1) * n(3) ^ 2 * n(5) + - 1.48346E-11 * n(1) * n(3) ^ 2 * n(6) + 0.000110143 * n(1) * n(3) * n(4) ^ 2 + 0.0000487937 * n(1) * n(3) * n(5) ^ 2 + 0.021921 * n(1) * n(4) * n(5) ^ 2 + 0.00000346283 * n(1) * n(5) ^ 2 * n(6)

R14 = -0.000000000177114 * n(2) ^ 2 * n(3) ^ 2 + 0.0000000210209 * n(2) ^ 2 * n(3) * n(4) + - 0.0000000818784 * n(2) ^ 2 * n(3) * n(5) + 0.0000376644 * n(2) ^ 2 * n(4) * n(5) + -9.31503E-13 * n(2) ^ 2 * n(6) ^ 2 + 0.000106985 * n(2) * n(3) * n(4) ^ 2 + 0.055031 * n(2) * n(4) ^ 2 * n(5) + -0.035459 * n(2) * n(4) * n(5) ^ 2 + 0.0000755324 * n(3) ^ 2 * n(4) * n(5) + -0.0000189566 * n(3) ^ 2 * n(5) ^ 2 + 0.040084 * n(3) * n(4) ^ 2 * n(5) + -0.00000000234695 * n(3) * n(4) * n(6) ^ 2 + 17.20136 * n(4) ^ 2 * n(5) ^ 2 + 0.004504 * n(4) * n(5) ^ 2 * n(6)

R15 = 0.000000561702 * n(4) * n(5) * n(6) ^ 2 + 0.0000000479781 * n(5) ^ 2 * n(6) ^ 2 + - 0.00000000944116 * n(0) ^ 3 * n(1) + 0.0000000115184 * n(0) ^ 3 * n(2) + -0.0000000158032 * n(0) ^ 3 * n(3) + 0.00000536878 * n(0) ^ 3 * n(4) + 0.00000362103 * n(0) ^ 3 * n(5) + 0.00000000106752 * n(0) ^ 3 * n(6) + -0.31887 * n(0) * n(4) ^ 3 + -0.00358755 * n(0) * n(5) ^ 3 + 0.000000000155012 * n(1) ^ 3 * n(2) + 0.000000025693 * n(1) ^ 3 * n(5) + 8.47252E-11 * n(1) * n(2) ^ 3 + 0.0000000269769 * n(2) ^ 3 * n(4)

R16 = -0.040793 * n(2) * n(4) ^ 3 + 0.013965 * n(2) * n(5) ^ 3 + 0.0000000679358 * n(3) ^ 3 * n(4) + 0.00494237 * n(4) ^ 3 * n(6) + 0.000439059 * n(5) ^ 3 * n(6) + 3.26886E-11 * n(5) * n(6) ^ 3 + 0.0000000400428 * n(0) ^ 4 + -1.89276E-11 * n(1) ^ 4 + -3.18337E-11 * n(3) ^ 4 + 25.12615 * n(4) ^ 4 + 2.44964 * n(5) ^ 4 + -7.13436E-15 * n(6) ^ 4 + 0 + 0

Model_2_Clear = R1 + R2 + R3 + R4 + R5 + R6 + R7 + R8 + R9 + R10 + R11 + R12 + R13 + R14 + R15 + R16

### Model_3_Clear

n(0) = percent persistence (organism level); n(1) = form factor mean (organism level); n(2) = density – levels mean (well level) ; n(3) = model_1 (organism level) ; n(4) = model_2_clear (organism level)

R1 = 32.52402 + -0.54831 * n(0) + 55.81306 * n(1) + -0.029513 * n(2) + 13.34291 * n(3) + 13.05986 * n(4)

R2 = 0.16126 * n(0) * n(1) + 0.000562305 * n(0) * n(2) + 0.11982 * n(0) * n(3) + -0.08123 * n(0) * n(4) + -0.062883 * n(1) * n(2) + 5.31126 * n(1) * n(3)

R3 = -4.81 * n(1) * n(4) + -0.021003 * n(2) * n(3) + -0.00540019 * n(2) * n(4) + -0.70453 * n(3) * n(4) + - 0.00140765 * n(0) ^ 2 + 9.04074 * n(1) ^ 2

R4 = 0.00000723861 * n(2) ^ 2 + 0.11459 * n(3) ^ 2 + -1.78504 * n(4) ^ 2 + -0.0000819342 * n(0) * n(1) * n(2) + -0.021891 * n(0) * n(1) * n(3) + 0.013961 * n(0) * n(1) * n(4)

R5 = -0.0000529987 * n(0) * n(2) * n(3) + 0.0000474044 * n(0) * n(2) * n(4) + 0.000387712 * n(0) ^ 2 * n(1) + 0.000000947006 * n(0) ^ 2 * n(2) + 0.0000959058 * n(0) ^ 2 * n(3) + -0.000237815 * n(0) ^ 2 * n(4)

R6 = -0.04276 * n(0) * n(1) ^ 2 + -0.000000154351 * n(0) * n(2) ^ 2 + 0.00112266 * n(0) * n(4) ^ 2 + - 2.45185 * n(1) ^ 2 * n(3) + 2.50057 * n(1) ^ 2 * n(4) + 0.0000161108 * n(1) * n(2) ^ 2

R7 = 0.40533 * n(1) * n(4) ^ 2 + 0.00000632871 * n(2) ^ 2 * n(3) + 0.00083054 * n(2) * n(4) ^ 2 + - 0.45553 * n(3) ^ 2 * n(4) + 0.87222 * n(3) * n(4) ^ 2 + -0.0000031075 * n(0) ^ 3

R8 = -2.84358 * n(1) ^ 3 + 0.14201 * n(3) ^ 3 + -0.38692 * n(4) ^ 3 + 0 + 0 + 0 Model_3_Clear = R1 + R2 + R3 + R4 + R5 + R6 + R7 + R8

### Model_2_Dark

n(0) = density – levels mean (organism level); n(1) = SD – levels mean (organism level); n(2) = percent persistence (well level); n(3) = SD – levels mean (well level); n(4) = pinch mean (organism level); n(5) = model_1 (organism level)

R1 = -361.12162 + 0.56347 * n(0) + 1.91376 * n(1) + -4.00323 * n(2) + -0.013169 * n(3) + -139.20978 * n(4) + 218.65312 * n(5) + -0.00153984 * n(0) * n(1) + 0.0052126 * n(0) * n(2) + -0.000454827 * n(0) * n(3) + -0.050456 * n(0) * n(4) + -0.18053 * n(0) * n(5) + -0.024777 * n(1) * n(2) + -0.00141908 * n(1) * n(3) + -0.12855 * n(1) * n(4) + -0.15031 * n(1) * n(5) + 0.011171 * n(2) * n(3) + 2.32431 * n(2) * n(4) + - 2.09687 * n(2) * n(5) + 0.53271 * n(3) * n(4) + -0.027043 * n(3) * n(5)

R2 = 12.99119 * n(4) * n(5) + -0.000309284 * n(0) ^ 2 + -0.000849861 * n(1) ^ 2 + 0.043616 * n(2) ^ 2 + -0.000056373 * n(3) ^ 2 + 124.14887 * n(4) ^ 2 + -56.25437 * n(5) ^ 2 + 0.0000128757 * n(0) * n(1) * n(2) + 0.00000098784 * n(0) * n(1) * n(3) + 0.000182684 * n(0) * n(1) * n(4) + 0.00002946 * n(0) * n(1) * n(5) + -0.00000513663 * n(0) * n(2) * n(3) + -0.000177285 * n(0) * n(2) * n(4) + 0.0000739509 * n(0) * n(2) * n(5) + -0.000231917 * n(0) * n(3) * n(4) + 0.0000143359 * n(0) * n(3) * n(5) + 0.051313 * n(0) * n(4) * n(5) + 0.00000505626 * n(1) * n(2) * n(3) + -0.000114908 * n(1) * n(2) * n(4) + 0.0036413 * n(1) * n(2) * n(5) + -0.000463003 * n(1) * n(3) * n(4)

R3 = 0.000195403 * n(1) * n(3) * n(5) + 0.027341 * n(1) * n(4) * n(5) + -0.0057082 * n(2) * n(3) * n(4) + - 0.00151504 * n(2) * n(3) * n(5) + -0.37303 * n(2) * n(4) * n(5) + -0.032101 * n(3) * n(4) * n(5) + 0.000000363092 * n(0) ^ 2 * n(1) + -0.00000126177 * n(0) ^ 2 * n(2) + 0.000000335253 * n(0) ^ 2 * n(3) + 0.00000930206 * n(0) ^ 2 * n(4) + 0.0000630895 * n(0) ^ 2 * n(5) + 0.000000625327 * n(0) * n(1) ^ 2 + -0.0000534651 * n(0) * n(2) ^ 2 + -0.0000000239318 * n(0) * n(3) ^ 2 + -0.00396581 * n(0) * n(4) ^ 2 + 0.023233 * n(0) * n(5) ^ 2 + 0.00000989143 * n(1) ^ 2 * n(2) + 0.00000105728 * n(1) ^ 2 * n(3) + - 0.000160825 * n(1) ^ 2 * n(4) + 0.0000106839 * n(1) ^ 2 * n(5) + 0.0000984379 * n(1) * n(2) ^ 2

R4 = 0.000000294665 * n(1) * n(3) ^ 2 + 0.026272 * n(1) * n(4) ^ 2 + -0.021385 * n(1) * n(5) ^ 2 + - 0.00000220276 * n(2) ^ 2 * n(3) + 0.00809653 * n(2) ^ 2 * n(4) + 0.018287 * n(2) ^ 2 * n(5) + - 0.00000561739 * n(2) * n(3) ^ 2 + -1.65978 * n(2) * n(4) ^ 2 + 0.50979 * n(2) * n(5) ^ 2 + 0.000142765 * n(3) ^ 2 * n(4) + 0.0000247584 * n(3) ^ 2 * n(5) + -0.10679 * n(3) * n(4) ^ 2 + 0.030073 * n(3) * n(5) ^ 2 + -54.71951 * n(4) ^ 2 * n(5) + 5.80519 * n(4) * n(5) ^ 2 + 0.000000067095 * n(0) ^ 3 + - 0.000000310388 * n(1) ^ 3 + -0.000236629 * n(2) ^ 3 + 0.000000169864 * n(3) ^ 3 + -6.52428 * n(4) ^ 3 + 5.28688 * n(5) ^ 3

R5 = 0.000000128977 * n(0) * n(1) * n(3) * n(4) + -0.0000000405581 * n(0) * n(1) * n(3) * n(5) + - 0.0000552203 * n(0) * n(1) * n(4) * n(5) + 0.000000890618 * n(0) * n(2) * n(3) * n(4) + 0.000000325049 * n(0) * n(2) * n(3) * n(5) + 0.00000169698 * n(1) * n(2) * n(3) * n(4) + -0.000000758555 * n(1) * n(2) * n(3) * n(5) + 0.00057653 * n(2) * n(3) * n(4) * n(5) + -6.57659E-11 * n(0) ^ 2 * n(1) ^ 2 + - 0.00000000245433 * n(0) ^ 2 * n(1) * n(2) + -0.00000000022752 * n(0) ^ 2 * n(1) * n(3) + 0.0000000214646 * n(0) ^ 2 * n(1) * n(4) + 0.0000000125412 * n(0) ^ 2 * n(2) ^ 2 + 0.000000000904972 * n(0) ^ 2 * n(2) * n(3) + 4.11355E-11 * n(0) ^ 2 * n(3) ^ 2 + 0.0000000265477 * n(0) ^ 2 * n(3) * n(4) + -0.0000104834 * n(0) ^ 2 * n(4) * n(5) + -0.00000366468 * n(0) ^ 2 * n(5) ^ 2 + - 0.00000000218385 * n(0) * n(1) ^ 2 * n(2) + -0.000000000175384 * n(0) * n(1) ^ 2 * n(3) + - 0.0000000678748 * n(0) * n(1) ^ 2 * n(4)

R6 = -0.0000000209727 * n(0) * n(1) * n(2) ^ 2 + -0.000000000109807 * n(0) * n(1) * n(3) ^ 2 + - 0.0000800513 * n(0) * n(1) * n(4) ^ 2 + 0.0000145427 * n(0) * n(1) * n(5) ^ 2 + -0.00000000641016 * n(0) * n(2) ^ 2 * n(3) + -0.00000476213 * n(0) * n(2) ^ 2 * n(4) + 0.00000000101544 * n(0) * n(2) * n(3) ^ 2 + 0.000304876 * n(0) * n(2) * n(4) ^ 2 + -0.0000663645 * n(0) * n(2) * n(5) ^ 2 + -0.0000000404017 * n(0) * n(3) ^ 2 * n(4) + 0.0000394658 * n(0) * n(3) * n(4) ^ 2 + -0.00000946448 * n(0) * n(3) * n(5) ^ 2 + 0.013504 * n(0) * n(4) ^ 2 * n(5) + -0.00662779 * n(0) * n(4) * n(5) ^ 2 + -0.000000035197 * n(1) ^ 2 * n(2) ^ 2 + -0.0000000101636 * n(1) ^ 2 * n(2) * n(3) + 0.000000190077 * n(1) ^ 2 * n(3) * n(4) + 0.0000734302 * n(1) ^ 2 * n(4) ^ 2 + 0.0000483842 * n(1) ^ 2 * n(4) * n(5) + -0.0000000182885 * n(1) * n(2) ^ 2 * n(3) + -0.0000227993 * n(1) * n(2) ^ 2 * n(5)

R7 = 0.00000000324656 * n(1) * n(2) * n(3) ^ 2 + -0.0000941191 * n(1) * n(2) * n(5) ^ 2 + - 0.0000000755622 * n(1) * n(3) ^ 2 * n(5) + 0.010924 * n(1) * n(4) * n(5) ^ 2 + 0.0000000126476 * n(2) ^ 2 * n(3) ^ 2 + 0.0000114672 * n(2) ^ 2 * n(3) * n(4) + 0.00286912 * n(2) ^ 2 * n(4) ^ 2 + -0.00348402 * n(2) ^ 2 * n(4) * n(5) + -0.00275424 * n(2) ^ 2 * n(5) ^ 2 + 0.000000519982 * n(2) * n(3) ^ 2 * n(5) + 0.000450946 * n(2) * n(3) * n(4) ^ 2 + 0.31246 * n(2) * n(4) ^ 2 * n(5) + 3.02457 * n(4) ^ 2 * n(5) ^ 2 + - 2.00237E-11 * n(0) ^ 3 * n(1) + -0.000000000070125 * n(0) ^ 3 * n(3) + -0.00000000793741 * n(0) ^ 3 * n(5) + -0.000000000158606 * n(0) * n(1) ^ 3 + 0.0000000785055 * n(0) * n(2) ^ 3 + -9.21427E-11 * n(0) * n(3) ^ 3 + -0.011845 * n(0) * n(4) ^ 3 + 0.00000000297869 * n(1) ^ 3 * n(2)

R8 = -0.000000000243674 * n(1) ^ 3 * n(3) + -0.0000000759836 * n(1) ^ 3 * n(5) + -0.00000000015798 * n(1) * n(3) ^ 3 + 0.031905 * n(1) * n(4) ^ 3 + -0.00308156 * n(1) * n(5) ^ 3 + -0.0000000397407 * n(3) ^ 3 * n(4) + -0.0000000277537 * n(3) ^ 3 * n(5) + -0.0022742 * n(3) * n(5) ^ 3 + -0.72285 * n(4) * n(5) ^ 3 + -3.17195E-12 * n(0) ^ 4 + 0.000000000456645 * n(1) ^ 4 + 0.000000336111 * n(2) ^ 4 + 7.93096E- 11 * n(3) ^ 4 + 6.09015 * n(4) ^ 4 + -0.31393 * n(5) ^ 4 + 0 + 0 + 0 + 0 + 0 + 0

Model_2_Dark = R1 + R2 + R3 + R4 + R5 + R6 + R7 + R8

### Model_3_Dark

n(0) = percent persistence (organism level); n(1) = x-y position stdev (organism level); n(2) = area mean (organism level); n(3) = area stdev (organism level); n(4) = SD – levels stdev (organism level); n(5) model_1 (organism level); n(6) model_2 (organism level);

R1 = -4.61017 + 0.092707 * n(0) + 0.35653 * n(1) + 0.000750986 * n(2) + -0.00452111 * n(3) + 0.011807 * n(4) + 5.71269 * n(5) + -1.49563 * n(6) + -0.00976856 * n(0) * n(1) + -0.0000263884 * n(0) * n(2) + 0.000135389 * n(0) * n(3) + 0.000208516 * n(0) * n(4) + -0.08961 * n(0) * n(5) + 0.013908 * n(0) * n(6)

R2 = 0.000111107 * n(1) * n(2) + -0.000521866 * n(1) * n(3) + -0.02206 * n(1) * n(4) + -0.39836 * n(1) * n(5) + -0.32854 * n(1) * n(6) + -0.0000000992238 * n(2) * n(3) + 0.0000185838 * n(2) * n(4) + - 0.000983153 * n(2) * n(5) + 0.000152417 * n(2) * n(6) + 0.0000626892 * n(3) * n(4) + 0.000877242 * n(3) * n(5) + 0.00418134 * n(3) * n(6) + 0.00787532 * n(4) * n(5) + 0.045654 * n(4) * n(6)

R3 = 0.55274 * n(5) * n(6) + -0.00027944 * n(0) ^ 2 + -0.013183 * n(1) ^ 2 + 0.0000000794713 * n(2) ^ 2 + 0.00000436954 * n(3) ^ 2 + -0.000737969 * n(4) ^ 2 + -1.60087 * n(5) ^ 2 + 1.28554 * n(6) ^ 2 + 0.00000121589 * n(0) * n(1) * n(2) + 0.000000680117 * n(0) * n(1) * n(3) + 0.000063997 * n(0) * n(1) * n(4) + 0.00737386 * n(0) * n(1) * n(5) + 0.00629009 * n(0) * n(1) * n(6) + -0.0000000203623 * n(0) * n(2) * n(3)

R4 = 0.000000283108 * n(0) * n(2) * n(4) + 0.0000214193 * n(0) * n(2) * n(5) + -0.00000950404 * n(0) * n(2) * n(6) + 0.000000257292 * n(0) * n(3) * n(4) + -0.0000845602 * n(0) * n(3) * n(5) + -0.0000444008 * n(0) * n(3) * n(6) + 0.00015789 * n(0) * n(4) * n(5) + -0.000189701 * n(0) * n(4) * n(6) + 0.0000000668764 * n(1) * n(2) * n(3) + -0.00000226437 * n(1) * n(2) * n(4) + -0.0000495035 * n(1) * n(2) * n(5) + -0.0000570377 * n(1) * n(2) * n(6) + -0.0000110973 * n(1) * n(3) * n(4) + -0.0000984269 * n(1) * n(3) * n(5)

R5 = -0.000547909 * n(1) * n(3) * n(6) + 0.000826802 * n(1) * n(4) * n(5) + -0.00174602 * n(1) * n(4) * n(6) + -0.034571 * n(1) * n(5) * n(6) + -0.0000000183872 * n(2) * n(3) * n(4) + 0.00000080816 * n(2) * n(3) * n(5) + 0.0000000300624 * n(2) * n(3) * n(6) + -0.0000272219 * n(2) * n(4) * n(5) + -0.0000091387 * n(2) * n(4) * n(6) + -0.000119087 * n(2) * n(5) * n(6) + 0.0000690982 * n(3) * n(4) * n(5) + 0.0000172096 * n(3) * n(4) * n(6) + -0.000273843 * n(3) * n(5) * n(6) + 0.00861831 * n(4) * n(5) * n(6)

R6 = 0.0000303932 * n(0) ^ 2 * n(1) + 0.000000156693 * n(0) ^ 2 * n(2) + -0.000000170271 * n(0) ^ 2 * n(3) + -0.0000168954 * n(0) ^ 2 * n(4) + 0.000243014 * n(0) ^ 2 * n(5) + -0.0000186158 * n(0) ^ 2 * n(6) + -0.000473015 * n(0) * n(1) ^ 2 + -0.000000000917335 * n(0) * n(2) ^ 2 + 0.0000000098126 * n(0) * n(3) ^ 2 + 0.00000251944 * n(0) * n(4) ^ 2 + 0.017146 * n(0) * n(5) ^ 2 + -0.000841143 * n(0) * n(6) ^ 2 + 0.0000110572 * n(1) ^ 2 * n(2) + 0.0000145286 * n(1) ^ 2 * n(3)

R7 = 0.00205469 * n(1) ^ 2 * n(4) + 0.040184 * n(1) ^ 2 * n(5) + 0.060707 * n(1) ^ 2 * n(6) + - 0.0000000343855 * n(1) * n(2) ^ 2 + 0.000000215682 * n(1) * n(3) ^ 2 + 0.000447475 * n(1) * n(4) ^ 2 +

0.241 * n(1) * n(5) ^ 2 + 0.28481 * n(1) * n(6) ^ 2 + 0.000000000152796 * n(2) ^ 2 * n(3) + - 0.000000000242684 * n(2) ^ 2 * n(4) + 0.00000000363359 * n(2) ^ 2 * n(5) + 0.00000012185 * n(2) ^ 2 * n(6) + -0.000000000959079 * n(2) * n(3) ^ 2 + 0.00000018851 * n(2) * n(4) ^ 2

R8 = 0.000173728 * n(2) * n(5) ^ 2 + -0.000284603 * n(2) * n(6) ^ 2 + 0.0000000102328 * n(3) ^ 2 * n(4) + -0.0000022087 * n(3) ^ 2 * n(5) + -0.0000000934567 * n(3) ^ 2 * n(6) + -0.00000105153 * n(3) * n(4) ^ 2 + -0.0000934181 * n(3) * n(5) ^ 2 + -0.00295776 * n(3) * n(6) ^ 2 + 0.000423315 * n(4) ^ 2 * n(5) + -0.0000799979 * n(4) ^ 2 * n(6) + -0.00450556 * n(4) * n(5) ^ 2 + -0.02169 * n(4) * n(6) ^ 2 + 0.26604 * n(5) * n(6) ^ 2 + -0.00000126558 * n(0) ^ 3

R9 = -0.0039262 * n(1) ^ 3 + -4.15924E-12 * n(2) ^ 3 + -5.47209E-11 * n(3) ^ 3 + -0.0000191162 * n(4) ^ 3 + 0.059443 * n(5) ^ 3 + -0.49754 * n(6) ^ 3 + -0.000000000383815 * n(0) * n(1) * n(2) * n(3) + - 0.000000750809 * n(0) * n(1) * n(2) * n(5) + -0.00000073033 * n(0) * n(1) * n(2) * n(6) + 0.00000000066314 * n(0) * n(1) * n(3) * n(4) + 0.00000157802 * n(0) * n(1) * n(3) * n(5) + 0.00000032997 * n(0) * n(1) * n(3) * n(6) + 0.0000444528 * n(0) * n(1) * n(4) * n(5) + 0.0000138956 * n(0) * n(1) * n(4) * n(6)

R10 = 7.76253E-11 * n(0) * n(2) * n(3) * n(4) + 0.00000000959345 * n(0) * n(2) * n(3) * n(5) + 0.00000000542978 * n(0) * n(2) * n(3) * n(6) + -0.000000138988 * n(0) * n(2) * n(4) * n(5) + - 0.000000255355 * n(0) * n(3) * n(4) * n(5) + 0.000000000366051 * n(1) * n(2) * n(3) * n(4) + - 0.0000000198672 * n(1) * n(2) * n(3) * n(5) + 0.0000000271726 * n(1) * n(2) * n(3) * n(6) + 0.000000965152 * n(1) * n(2) * n(4) * n(5) + 0.00000090072 * n(1) * n(2) * n(4) * n(6) + 0.00000812757 * n(1) * n(3) * n(4) * n(5) + 0.00000797685 * n(1) * n(3) * n(4) * n(6) + 0.000103226 * n(1) * n(3) * n(5) * n(6) + -0.0000207396 * n(3) * n(4) * n(5) * n(6)

R11 = -0.00000195856 * n(0) ^ 2 * n(1) ^ 2 + 0.0000000269254 * n(0) ^ 2 * n(1) * n(3) + -0.0000508801 * n(0) ^ 2 * n(1) * n(5) + 2.49496E-11 * n(0) ^ 2 * n(2) * n(3) + -0.0000000742936 * n(0) ^ 2 * n(2) * n(5) + 0.0000000306992 * n(0) ^ 2 * n(2) * n(6) + -0.000000000135004 * n(0) ^ 2 * n(3) ^ 2 + - 0.00000000292662 * n(0) ^ 2 * n(3) * n(4) + 0.000000278337 * n(0) ^ 2 * n(3) * n(5) + 0.00000997994 * n(0) ^ 2 * n(4) * n(5) + -0.0000639369 * n(0) ^ 2 * n(6) ^ 2 + 0.0000000558002 * n(0) * n(1) ^ 2 * n(2) + - 0.000000143373 * n(0) * n(1) ^ 2 * n(3) + -0.00000414178 * n(0) * n(1) ^ 2 * n(4)

R12 = -0.000000000741061 * n(0) * n(1) * n(3) ^ 2 + -0.00000126359 * n(0) * n(1) * n(4) ^ 2 + - 0.00203358 * n(0) * n(1) * n(6) ^ 2 + -2.37659E-13 * n(0) * n(2) ^ 2 * n(3) + 0.00000000033799 * n(0) * n(2) ^ 2 * n(6) + 1.08851E-12 * n(0) * n(2) * n(3) ^ 2 + -0.00000000142941 * n(0) * n(2) * n(4) ^ 2 + - 0.00000293071 * n(0) * n(2) * n(5) ^ 2 + 0.00000237668 * n(0) * n(2) * n(6) ^ 2 + -8.93754E-11 * n(0) * n(3) ^ 2 * n(4) + 0.00000000419135 * n(0) * n(3) * n(4) ^ 2 + 0.00000995675 * n(0) * n(3) * n(6) ^ 2 + - 0.000271624 * n(0) * n(4) * n(5) ^ 2 + -0.0000000015755 * n(1) ^ 2 * n(2) * n(3)

R13 = -0.0000076574 * n(1) ^ 2 * n(2) * n(5) + -0.00000993414 * n(1) ^ 2 * n(2) * n(6) + - 0.00000000914334 * n(1) ^ 2 * n(3) ^ 2 + 0.00000238695 * n(1) ^ 2 * n(3) * n(5) + -0.000688278 * n(1) ^ 2 * n(4) * n(5) + -0.000619047 * n(1) ^ 2 * n(4) * n(6) + -0.018618 * n(1) ^ 2 * n(6) ^ 2 + 0.0000000176265 * n(1) * n(2) ^ 2 * n(5) + 0.0000000156328 * n(1) * n(2) ^ 2 * n(6) + 0.000019277 * n(1) * n(2) * n(6) ^ 2 + -0.00000000110944 * n(1) * n(3) ^ 2 * n(4) + -0.0000000105332 * n(1) * n(3) ^ 2 * n(5) + -0.0000000252387 * n(1) * n(3) ^ 2 * n(6) + -0.0000000702263 * n(1) * n(3) * n(4) ^ 2

R14 = 0.00012233 * n(1) * n(3) * n(6) ^ 2 + -0.000128546 * n(1) * n(4) ^ 2 * n(5) + -0.000106207 * n(1) * n(4) ^ 2 * n(6) + 0.0030741 * n(1) * n(4) * n(6) ^ 2 + -0.087995 * n(1) * n(5) * n(6) ^ 2 + 7.37175E-13 * n(2) ^ 2 * n(3) * n(4) + -9.59323E-11 * n(2) ^ 2 * n(3) * n(5) + -6.02571E-11 * n(2) ^ 2 * n(3) * n(6) + 2.50502E-12 * n(2) * n(3) ^ 2 * n(4) + 0.000000000442336 * n(2) * n(3) ^ 2 * n(5) + 9.93414E-11 * n(2) * n(3) ^ 2 * n(6) + -6.40403E-11 * n(2) * n(3) * n(4) ^ 2 + -0.000000292517 * n(2) * n(3) * n(5) ^ 2 + 0.00000955825 * n(2) * n(4) * n(5) ^ 2

R15 = 0.000000000350581 * n(3) ^ 2 * n(4) ^ 2 + -0.0000000168097 * n(3) ^ 2 * n(4) * n(5) + - 0.0000000161471 * n(3) ^ 2 * n(4) * n(6) + -0.0000000692311 * n(3) ^ 2 * n(5) * n(6) + -0.000020757 * n(3) * n(4) * n(5) ^ 2 + 0.000617916 * n(3) * n(5) * n(6) ^ 2 + 0.00021377 * n(4) ^ 2 * n(5) * n(6) + 0.000000339072 * n(0) ^ 3 * n(1) + -0.00000000207837 * n(0) ^ 3 * n(3) + 0.000019645 * n(0) * n(1) ^ 3 + 4.45966E-14 * n(0) * n(2) ^ 3 + 9.04952E-13 * n(0) * n(3) ^ 3 + 0.0000000763313 * n(0) * n(4) ^ 3 + 0.00283387 * n(0) * n(6) ^ 3

R16 = 0.000000219559 * n(1) ^ 3 * n(2) + 0.000000953974 * n(1) ^ 3 * n(3) + -0.0000133042 * n(1) ^ 3 * n(4) + 0.000882434 * n(1) ^ 3 * n(6) + -0.067623 * n(1) * n(5) ^ 3 + 2.58842E-15 * n(2) ^ 3 * n(3) + - 9.82343E-12 * n(2) ^ 3 * n(6) + 0.0000000054835 * n(3) * n(4) ^ 3 + 0.000463001 * n(3) * n(5) ^ 3 + 0.000345934 * n(3) * n(6) ^ 3 + 0.00000320697 * n(4) ^ 3 * n(5) + 0.00000317484 * n(4) ^ 3 * n(6) + - 0.2124 * n(5) * n(6) ^ 3 + -7.87648E-17 * n(3) ^ 4

R17 = 0.12069 * n(6) ^ 4

Model_3_Dark = R1 + R2 + R3 + R4 + R5 + R6 + R7 + R8 + R9 + R10 + R11 + R12 + R13 + R14 + R15 + R16 + R17

### MySQL Database

The experiment is defined in the database (Figure 6) by updating the “project”, “assay”, “version”, “plate”, “well”, “compound”, “lot”, and “session” tables. In the database, “version” is the version of the assay, “lot” is the version of the compound, and “session” stores the location of data files (data CSV, image acquisition XDCE), the date of the start of the iteration, and the status of the data processing. Next, the timestamp of every image is parsed and loaded into the “time” table. Then the file from data pre-processing is imported into the “raw” table and linked to the “time” table. The “time” table becomes the route to travel to different points in time within the same well whereas “session” is useful in separating the campaign into experimental iterations. The “time” table also stores flags indicating a well should not be considered for analysis. From the “raw” table, the data is processed and loaded into “frag”, “worm”, “result”, and “effect”.

Data in the “raw” table were reorganized into “fragments”. Organisms in each time frame were linked to organisms in subsequent time frames using the method described in “Object Classification” steps 1 through 5. There is a chance that a link cannot be found or a link will be found a later time point. These gaps in the linking operation produce fragments or varying size. If the fragments are too small (< 4 time points) the fragments are not used for analysis. Four or more time points per fragment ensures 3 or more data points to estimate rate which is the mean amount of change per time frame.

Long enough fragments in the “fragment” table were reorganized into “worms”. The “static” mean and standard deviation of each feature for all worm time points were calculated. The “rate” mean and standard deviation of each feature’s absolute change between time points were calculated. The “frequency” of each feature is determined by measuring the number of directional changes per time. Changes in value that are below the system noise were carried over if the change continues in the same direction. (**Table 2**).

The worms in the “worm” table are analyzed at the well level to provide results to the “results” table. The mean and standard deviation for “static”, “rate”, and frequency modes were calculated.

Results in the “statistic” table are used to calculate effect sizes and Mahalanobis Distance in the “effects” table and “Mahalanobis” table. The Glass effect size is used to calculate effect sizes. The Mahalanobis Distance is calculated for the “static”, “rate” and “frequency” categories separately, the combination of “static” and “rate” categories, and the full combination of “static”, “rate”, and “frequency” categories. Degeneracy data is located in the “degeneracy” table. Degeneracy is the number of dark worms divided by the total worms per well.

**Figure 6.**
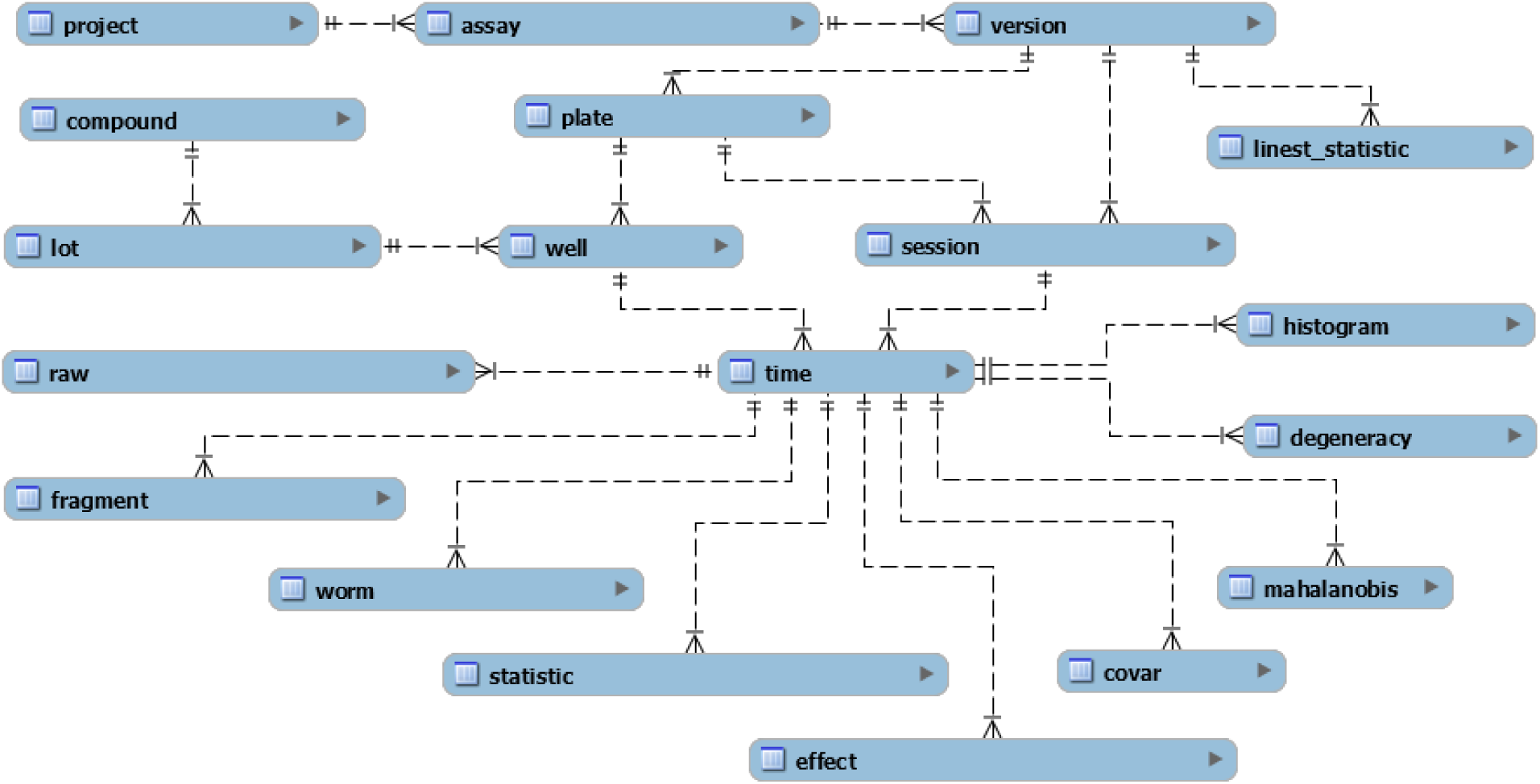
Best mask to represent the object. Image shows the best mask to represent the object from the set of 12 possible masks. Blue outlines are derived from the clear body workflow, green outlines are from the tegument outliner, and red outlines are from the dark body workflow.

**Table 2.** A measure of system noise per feature. The table shows values for system noise per feature. The system noise values are used to detect changes in motion that can be used to calculate frequency.

| List of Features | Cutoff for "Clear" | Cutoff for "Opaque" |
| --- | --- | --- |
| area | 52.75 | 152 |
| weighted_position_x | 1.6 | 2.2 |
| weighted_position_y | 1.2 | 2.2 |
| median_diameter | 1.456 | 2.11 |
| length | 1.686 | 3.058 |
| form_factor | 0.0125 | 0.0155 |
| perimeter | 3.4485 | 5.569 |
| straight_chord | 1.659 | 2.8585 |
| curved_chord | 2.8105 | 4.372 |
| custom_bend | 0.0205 | 0.024 |
| custom_pinch | 0.025 | 0.0315 |
| custom_wave | 0.0225 | 0.0225 |
| mass | 7475.5 | 26395.5 |
| weighted_moment_of_inertia | 1.654 | 3.49 |
| density_levels | 4.445 | 2.3275 |
| standard_deviation_of_density_levels | 3.4245 | 1.8145 |
| angle | 2.059 | 3.232 |

### Graphical User Interface (GUI) of SchistoView

The GUI of SchistoView (**Figure 7**) allows hierarchical navigation of the experimental results. A Mahalanobis Distance or percent degeneracy heatmap of the assay plate provides a high-level summary. Selecting on a well in the heatmap updates the effect size heatmap which provides insight into which features contribute to the well result. An effect size for a given feature may then be selected to update the histogram of the well, the effect size plot over time, and the effect size dose response plot for that feature. Frequency is visualized with an estimate of the waveform and displays the wavelength and amplitude versus negative control DMSO. An image of the well at the indicated time point is displayed.

The user can toggle the campaigns, iterations within the campaign, data grouping, type of Mahalanobis Distance (or percent degeneracy), effect category, and feature.

SchistoView was created using Excel VBA forms and a MySQL connector to query the database in real-time.

**Figure 7.**
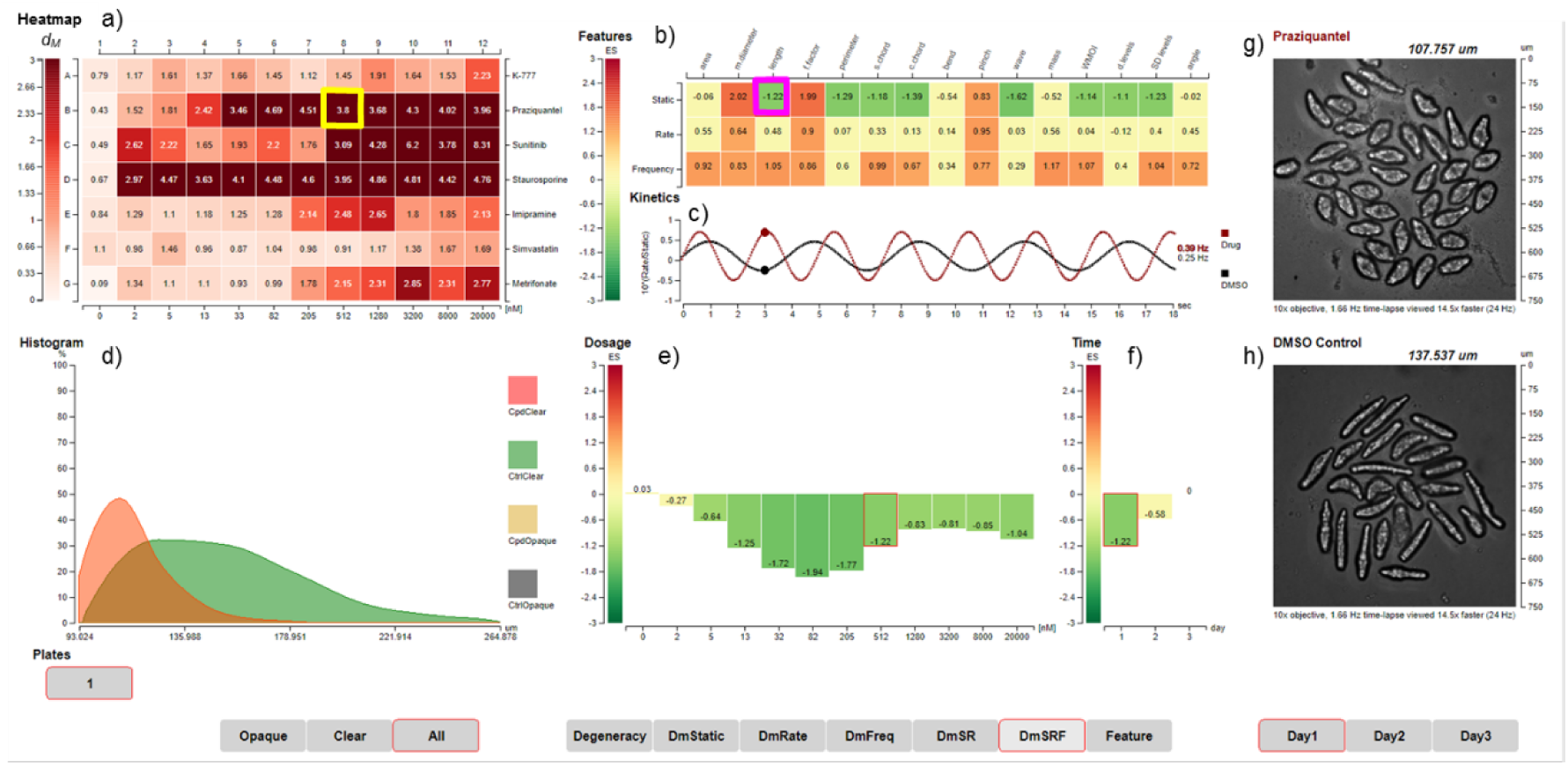
Screenshot of the SchistoView graphical user interface. Selected data are shown to illustrate the hierarchical approach to visualization. **(a)** Heat map of Mahalanobis distances (*dM*) for seven test drugs arrayed over an 11-point 2.5-fold dilution series from 2 nM in column 2 to 20 μM in column 12. Drugs, from top to bottom, are, K11777, PZQ, sunitinib, staurosporine, imipramine, simvastatin and metrifonate. DMSO controls are arrayed in column 1 and are shown as the average *dM* (0.77) for all DMSO controls. A *dM* of 1.61 is significantly different (3 SD) from control. Clicking on coordinate B8 (identified by the yellow square: 512 nM PZQ) populates panels (**b**) and (**g**) (see below). **(b)** Heat map showing the effect sizes (ES) for static, rate and frequency, after exposure to 512 nM PZQ for 2 h, *i.e*., the selected well from (**a**). Three sets of 15 features are arrayed in rows and columns, respectively. Clicking on the intersection of the length feature and static mode (magenta box) in (**b**) populates panels (**c**) through (**f**) and the underlying data. (**c**) Calculated waveforms defined by the range of length (amplitude) and frequency of length contraction (frequency). DMSO control worms are slower moving (lower frequency) than those treated with 512 nM PZQ (red line). (**d**) Histogram displaying the distribution of static length for DMSO control worms (green) and PZQ-treated worms (orange). (**e**) Bar graph depicting the ES for static length after PZQ treatment across 11 concentrations (second row in (**a**)). (**f**) Bar graph depicting the ES for static length in the 512 nM PZQ treatment across the three days of measurement. **(g)** First image from time-lapsed movie of the well highlighted in (**a**); in the live SchistoView, the 30-frame movie is looped. (**h**) as for (**g**) except for the DMSO control.

